# Two different pathways for initiation of *Trichoderma reesei* Rad51-only meiotic recombination

**DOI:** 10.1101/644443

**Authors:** Wan-Chen Li, Yu-Chien Chuang, Chia-Ling Chen, Ljudmilla Timofejeva, Wen-Li Pong, Yu-Jie Chen, Chih-Li Wang, Ting-Fang Wang

**Affiliations:** Taiwan International Graduate Program in Molecular and Cellular Biology, Academia Sinica, Taipei 115, Taiwan; Institute of Life Sciences, National Defense Medical Center, Taipei 115, Taiwan; Institute of Molecular Biology, Academia Sinica, Taipei 115, Taiwan; Department of Plant Pathology, National Chung-Hsing University, Taichung 402, Taiwan

**Keywords:** Meiosis, Homologous recombination, Double strand break (DSB), Spo11, Topoisomerase II, Rad51, Dmc1, Synaptonemal complex, genetic interference, crossover homeostasis, *Trichoderma reesei*

## Abstract

**Background:** Meiotic recombination is mainly, but not exclusively, initiated by Spo11-induced double strand breaks (DSBs) in some sexual eukaryotes. DSBs are repaired by one or two RecA-like recombinases (ubiquitous Rad51 and meiosis-specific Dmc1). In yeast and mammals, Dmc1 is superior to Rad51 in tolerating mismatched sequences during highly polymorphic hybrid meiosis. The mechanisms underlying Rad51-only meiotic recombination remain less studied.

**Results:** The Rad51-only filamentous fungus *Trichoderma reesei* has only one *spo11* gene. Removal of *spo11* from *T. reesei* genome does not affect normal sexual development, meiosis or chromosome synapsis, but results in decrease of interhomolog recombination products to 70%, crossover homeostasis and lower genetic interference. Our results also suggest that *T. reesei* Rad51, like yeast and mammalian Dmc1 (but not Rad51), can tolerate mismatched sequences during meiotic recombination. Moreover, Topoisomerase II might act redundantly (and predominantly) with Spo11 to initiate meiotic recombination.

**Conclusion:** We suggest that *T. reesei* is an emerging model for studying Spo11-independent and Rad51-only meiosis.

## Background

Homologous interactions (e.g., pairing, recombination and synapsis) between maternal and paternal homologous chromosomes are the central theme of meiosis because they generate genetic diversity in haploid gametes and, upon fertilization, in the offspring. The interhomolog crossover (CO) products are also required for proper segregation of parental chromosomes during the first meiotic division (reviewed in [1]).

In almost all studied model eukaryotes, Spo11-induced DNA double-strand breaks (DSBs) are responsible for initiation of meiotic recombination. Spo11 is a meiosis-specific Topoisomerase VII subunit A endonuclease [2, 3]. There is considerable evidence indicating that Spo11-independent DSBs and/or meiotic recombination proceed in some sexual eukaryotes [4–7]. Uniquely, as far as is known among sexual eukaryotes, dictyostelids (i.e., social amoebae) lack a *spo11* gene [8]. The molecular mechanisms underlying Spo11-independent DSBs remain less studied than for Spo11-dependent ones. A primary reason for this situation is the lack of an experimentally tractable model system in which normal meiosis can proceed in the absence of *spo11* to produce viable haploid gametes (e.g., the ascospores or sexual spores in fungi).

Sexual eukaryotes fall into two groups with respect to their RecA-like recombinases, which catalyze homologous pairing and strand-exchange reactions during homologous recombination. The first group (referred to as “Dual-RecA eukaryotes”) includes budding and fission yeast, higher plants and mammals. They all possess Rad51 and Dmc1, which cooperate during meiotic recombination. In addition to meiotic recombination Rad51 (as the only RecA strand-exchange protein) is responsible for mitotic recombination, whereas Dmc1 is meiosis-specific (reviewed in [9]). Recent single-molecular imaging experiments revealed that budding yeast and mammalian Dmc1 are superior to Rad51 in tolerating mismatched sequences during their strand exchange reaction [10, 11]. Consistent with this hypothesis, Dmc1-mediated recombination is more efficient than Rad51-mediated recombination in highly polymorphic diploid hybrid yeasts, e.g., SK1/S228c and YJM/S228c [12]. The “Rad51-only” eukaryotes include *D. melanogaster, C. elegans* and Pezizomycotina filamentous fungi (e.g., *Neurospora crassa*). The absence of Dmc1 in the “Rad51-only” organisms raises the intriguing question as to how interhomolog recombination is possible among highly diversified zygotes.

Like *Neurospora crassa, Trichoderma reesei* (teleomorph *Hypocrea jacorina*) is a Pezizomycotina filamentous fungus. QM6a is the ancestor of all *T. reesei* workhorse strains currently used for industrial production of lignocellulosic biomass-degrading enzymes and recombinant proteins (see review in [13]). QM6a was for a long time thought to be an asexual filamentous fungus. A milestone in this respect was the finding that QM6a has a *MAT1-2* locus and that it can be readily crossed with a female fertile CBS999.97(*MAT1-1*) strain [14]. Crosses of CBS999.97(*MAT1-1*) with QM6a or CBS999.97(*MAT1-2*) can induce rapid sexual development and generate reproductive structures. The round-shaped stromata (fruiting bodies) are composed of a thallus, with multiple flask-shaped perithecia inside. Each perithecium contains a bouquet of linear asci. The 16 ascospores in each ascus are generated from meiosis, followed by two rounds of post-meiotic mitosis [15].

QM6a has a small genome (~35 Mbp) and possseses only one copy of a RecA-like recombinase gene (*rad51* but not *dmc1*) [16]. In this study, we report that the industrial workhorse filamentous fungi *Trichoderma reesei* (teleomorph *Hypocrea jacorina*) is an ideal sexual eukaryote for studying the molecular mechanisms underlying Spo11-independent and Rad51-only meiotic recombination.

## Results

### Highest quality genome sequences yet of CBS999.97(*MAT1-1*) and CBS999.97(*MAT1-2*)

QM6a was originally isolated from one of the Solomon Islands, whereas *Hypocrea jacorina* CBS999.97 was sampled from French Guiana (see review in [17]). Due to geographical isolation, QM6a and CBS999.97 might harbor high levels of sequence variation. We had determined the complete genome sequences of QM6a [16]. In this study, both PacBio long reads and Illumina paired-end reads were applied to assemble the complete genome sequences of CBS999.97(*MAT1-1*) and CBS999.97(*MAT1-2*) (Table S1). Like QM6a, CBS999.97(*MAT1-1*) and CBS999.97(*MAT1-2*) have eight reference chromosomes (seven nuclear chromosomes plus the mitochondrial DNA). Of these, all seven nuclear chromosomes were essentially returned as a single, complete unitig (Table S1 and Fig. 1). Except for the left terminus of chromosome VI, all other chromosomal ends in CBS999.97(*MAT1-1*) and CBS999.97(*MAT1-2*) contain 8-13 telomeric repeats (i.e., TTAGGG at 3′-termini and the reverse complement CCCTAA at 5′-termini) (Table S2) [16]. Due to the occurrence of repeat-induced point mutation (RIP) in *T. reesei* [16], both CBS999.97 haploid genome sequences contain only 62 transposable elements (Table S3) but 2349 AT-rich blocks with length ≥ 500 base pairs (bp) (Table S4).

**Fig. 1.**
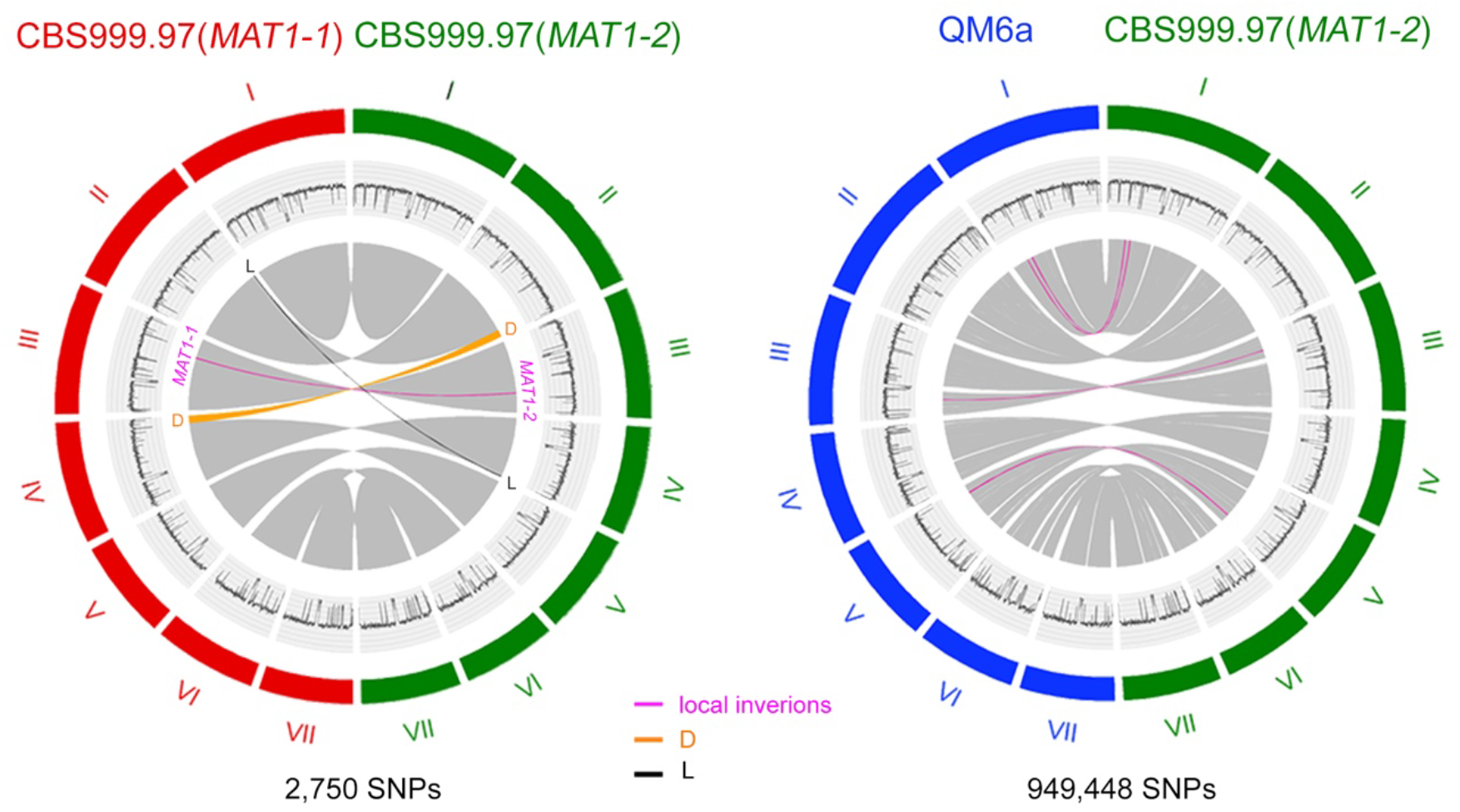
**Conserved synteny, inversions and translocations between** CBS999.97(*MAT1-1*) and CBS999.97(*MAT1-2*) (**left panel**) and between QM6a and CBS999.97(*MAT1-2*) (**right panel**). Collinearity of *Trichoderma reesei* genomes is depicted in grey in the inner circles of the diagrams. The D segment is indicated in orange, the L segment in black, and local inversions in magenta. The outer circles indicate the seven chromosomes (I-VII) of CBS999.97(*MAT1-1*) (in red), CBS999.97(*MAT1-2*) (in green) and QM6a (in blue). The GC contents (window size 5000 bp) of the seven chromosomes are shown in the middle traces.

### Reciprocal translocation in CBS999.97(*MAT1-1*)

We reported previously that sexual crosses of CBS999.97(*MAT1-1*) with QM6a or CBS999.97(*MAT1-2*) resulted in ~10% asci with 16 euploid ascospores and ~90% asci with two different types of segmentally aneuploid (SAN) ascospores [15]. The euploid progeny, as for QM6a and the two CBS999.97 parental strains, germinate to form mycelia with dark-green conidia (i.e., asexual spores). The first type of SAN ascospores can germinate but fail to undergo vegetative growth due to loss of the D segment (~0.5 Mb). The second type of SAN ascospores lack the L segment (~30 kb) but possess two D segments. Our high-quality genome sequences readily confirm the translocation event between the D segment and the L segment (Fig. 1). The D segment is located at the right terminus of chromosome II in QM6a and CBS999.97(*MAT1-2*) or at the right terminus of chromosome IV in CBS999.97(*MAT1-1*). In contrast, the L segment is located at the right terminus of chromosome IV in QM6a and CBS999.97(*MAT1-2*) or at the right terminus of chromosome II in CBS999.97(*MAT1-1*). Accordingly, CBS999.97(*MAT1-1*) and CBS999.97(*MAT1-2*) are referred to as CBS999.97(*MAT1-1*, II_L, IV_D) and CBS999.97(*MAT1-2*, II_D, IV_L), respectively [15].

Next, we applied pulsed field gel electrophoresis (PFGE) to separate the seven chromosomes of CBS999.97(*MAT1-1*, II_L, IV_D), CBS999.97(*MAT1-2*, II_D, IV_L) and two type II SAN progeny strains (SAN1 and SAN2; Fig. S1a). Southern hybridization with two DNA probes in the D segment and a DNA probe in the *pks4* gene confirmed that CBS999.97(*MAT1-1*) and CBS999.97(*MAT1-2*) contain D and L segments in the corresponding chromosomes. In contrast, in the two type II SAN strains, both chromosome II and chromosome IV contain a D segment but no L segment (Fig. S1b-S1d).

### Single nucleotide polymorphism (SNP) markers

The three high-quality genome sequences allow efficient SNP calling for genetic and comparative genomic studies. Using the high stringency SNP detection pipeline from the MUMmer alignment tools (http://mummer.sourceforge.net/), we identified SNP markers between CBS999.97(*MAT1-1*) and CBS999.97(*MAT1-2*) (Fig. 1 and Table S5). There are 935,999 SNPs between QM6a and CBS999.97(*MAT1-1*) (Table S6) and 949,448 SNPs between QM6a and CBS999.97(*MAT1-2*) (Fig. 1 and Table S7). Since all SNP markers are evenly distributed throughout the seven chromosomes in QM6a and CBS999.97(*MAT1-1*), these two strains are suitable for determining the genome-wide meiotic recombination landscape (see below).

It is worth noting that the degree of sequence heterozygosity (≥37 bps/SNP) between QM6a and CBS999.97(*MAT1-1*) is higher than that (≥200 bps/SNP) of the budding yeast SK1/S288c and YJM/S288c diploid hybrids [12]. Since *T. reesei* is a “Rad51-only” sexual eukaryote and crosses of CBS999.97(*MAT1-1*) with QM6a or CBS999.97(*MAT1-2*) can generate viable progeny, our results raise the intriguing possibility that *T. reesei* Rad51, like budding yeast Dmc1, might be superior to budding yeast Rad51 in tolerating mismatched sequences during the strand exchange reaction. However, a phylogenetic analysis revealed that all Rad51 proteins in six different “Rad51-only” eukaryotes are more similar in amino acid sequence to the Rad51 proteins than to the Dmc1 proteins in nine different “Dual-RecA” eukaryotes (Fig. S2).

### *T. reesei spo11*, unlike *rad51* or *sae2*, is dispensable for normal meiosis

Our high-quality genome sequences confirm that there is only one *spo11, rad51* (but not *dmc1*) and *sae2* in each of QM6a [16], CBS999.97(*MAT1-1*) and CBS999.97(*MAT1-2*) (this study). The conserved primary structures and well-aligned protein sequences with other fungal orthologs strongly support identification of these important meiotic genes in *T. reesei* (Fig. S3-S5). We then generated *rad51*Δ, *sae2*Δ and *spo11*Δ mutants by gene replacement based on homologous recombination and confirmed mutation by antibiotic selection, Southern hybridization or genomic polymerase chain reactions (Fig. S6).

*T. reesei* Rad51 is indispensable for repairing DSBs and/or DNA lesions during meiosis in *T. reesei*. Deletion of *rad51* resulted in no apparent effect on formation of fruiting bodies, perithecia or asci. Almost all *rad51*Δ asci exhibit meiotic prophase arrest and have only one nucleus (Fig. 2b). This meiotic prophase arrest phenotype is similar to that of *S. cerevisiae dmc1*Δ or *rad51*Δ *dmc1*Δ lines in which unrepaired ssDNA accumulates, leading to activation of Mec1^ATR^, an evolutionarily conserved DNA damage checkpoint kinase [18].

**Fig. 2.**
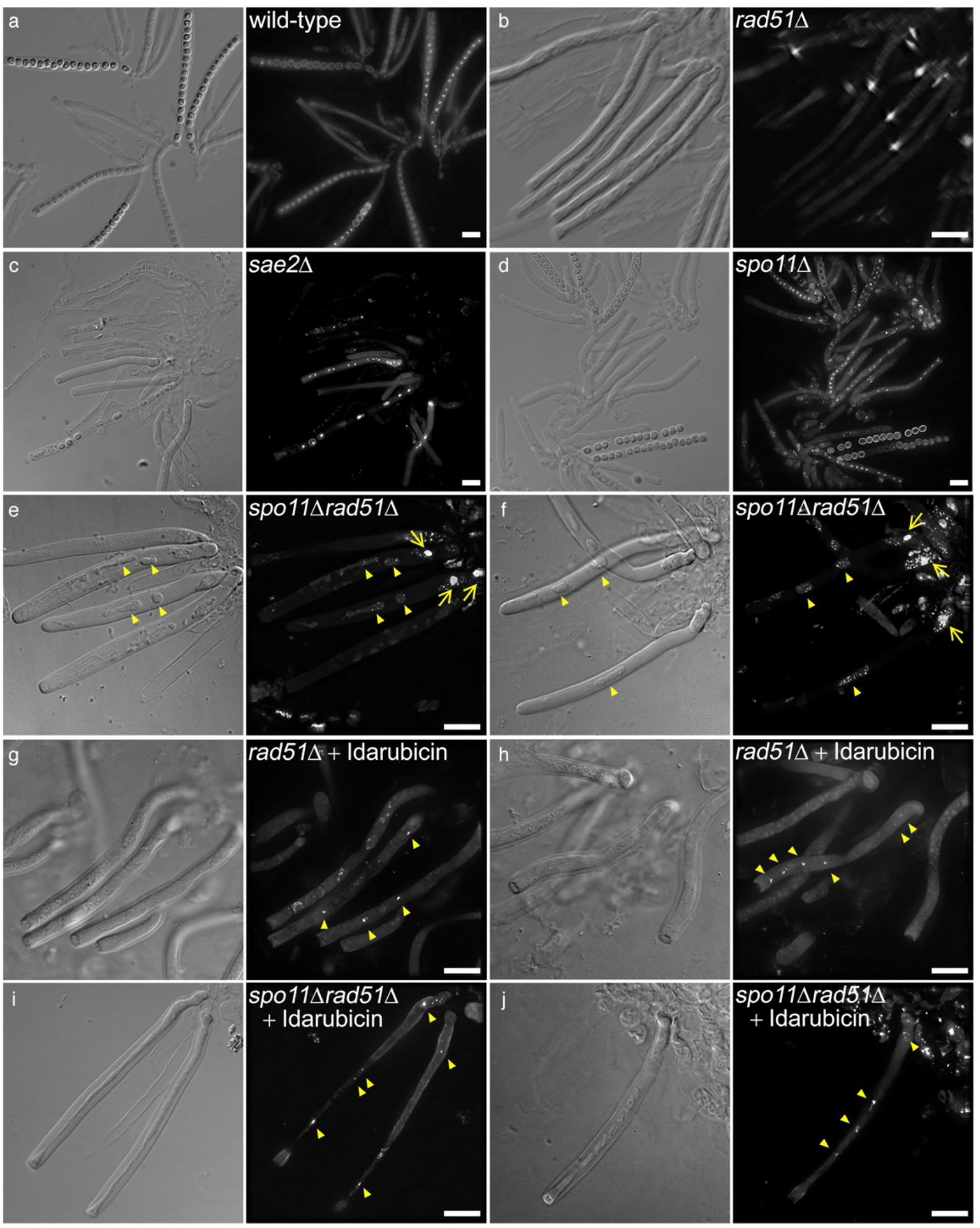
Visualization of asci from wild-type and mutant homozygous zygotes. Rosettes of asci were dissected from developing fruiting bodies, stained with 4’,6-diamidino-2-phenylindole (DAPI), and then visualized by fluorescence microscopy. Differential interference contrast (DIC) and DAPI fluorescent images are shown. Scale bar is 10 μm. (**a**) wild-type, (**b**) *rad51*Δ, (**c**) *sae2*Δ, (**d**) *spo11*Δ, (**e** and **f**) *spo11*Δ *rad51*Δ. The meiotic prophase arrest phenotypes of the *rad51*Δ asci (**g** & **h**) and the *spo111*Δ *rad51*Δ asci (**i** & **j**) were rescued (at least partly) upon addition of idarubicin, an effective inhibitor of fungal Top2. Yellow arrows indicate the single strong DAPI-staining focus in the *spo11*Δ *rad51*Δ asci (**e** & **f**). Yellow triangles indicate the DAPI signals in the *spo11*Δ *rad51*Δ asci (**e** & **f**) and upon addition of idarubicin (100 μM) to the *rad51*Δ asci (**g** & **h**) and the *spo11*Δ *rad51*Δ asci (**i** & **j**).

Sae2 (CtIP in mammals or Ctp1 in fission yeast) is a DNA endonuclease that collaborates with the Mre11-Rad50-Xrs/Nbs1 complex to remove a small oligo nucleotide(s) from free DSB ends, Spo11-conjugated DSBs and Topoisomerase II (Top2)-conjugated DSB ends [19–22]. *T. reesei* Sae2, like Rad51, is dispensable for the formation of fruiting bodies or perithecia. However, the *sae2*Δ perithecia contain more underdeveloped or short asci than those of wild-type and *rad51*Δ. Some *T. reesei sae2*Δ asci can still undergo the first or even second meiotic nuclear divisions. Though few *sae2*Δ asci formed mature ascospores, there was always fewer than sixteen ascospores (Fig. 2c). We then applied yeast tetrad dissection microscopy to isolate single *sae2*Δ ascospores. The majority of the *sae2*Δ ascospores (>57%) failed to germinate and form vegetative colonies on a malt extract agar (MEA) plate. Therefore, the *T. reesei sae2*Δ mutant is phenotypically similar to *S. cerevisiae sae2* null mutants, with unprocessed Spo11-conjugated DSBs in these latter activating another evolutionarily conserved DNA damage checkpoint kinase Tel1^ATM^ (but not Mec1^ATR^). Since the kinase activity of Tel1^ATM^ in budding yeast meiotic cells is much lower than that of Mec1^ATR^, the *sae2* null mutants exhibit much weaker meiotic prophase arrest phenotypes compared to *dmc1*Δ or the *rad51*Δ *dmc1*Δ double mutant. Accordingly, the majority of *S. cerevisiae sae2* null mutant cells (50-70%) can execute the first or even the second meiotic nuclear divisions with reasonable efficiency but hardly produce any viable spores [20].

The *T. reesei* Spo11 protein exhibits significant homology (33.8% identity in amino acid sequence) with the fission yeast *S. pombe* Rec12 protein. It contains five conserved motifs (motifs 1-5) and three evolutionarily conserved amino acid residues equivalent to those of *S. cerevisiae* Spo11: Tyr-135 (in motif 1), Glu-233 (in motif 3) and Asp-288 in motif 5 (Fig. S3). These three evolutionarily conserved amino acid residues are known to be essential for meiotic recombination in *S. cerevisiae* [23]. Surprisingly, deletion of *spo11* resulted in no apparent effect on the entire sexual development process, including formation of fruiting bodies, perithecia or asci, as well as the number of ascospores per ascus (Fig. 2d). We then sequentially isolated all 16 ascospores from an ascus. Our results reveal that spore viability, colony morphology and colony color were identical regardless of the presence or absence of *spo11* in the individually cultured ascospores (Table S8).

### Genome-wide detection of meiotic recombination products in *T. reesei*

Next, we explored genome-wide meiotic recombination profiles in the presence and absence of *spo11*. An in-house bioinformatics pipeline was established for SNP position genotyping and recombination analysis (see “Methods”). For efficacy assessment, the pipeline was first used to reanalyze the NGS data of budding yeast tetrads from a diploid SK1/S288c hybrid [12]. We could detect all non-crossover (NCO) and crossover (CO) recombination products (Table S9) using the ReCombine (v2.1) [24] and GroupEvents programs [25]. NCO is a gene conversion (GC) without exchange of flanking markers, whereas CO involves the exchange of flanking markers and possible GC. Median CO length, estimated by a cumulative frequency plot, is ~ 1 kb (see below). The long GC tracts observed in budding yeast meiotic COs arises from clustered Spo11-induced DSBs within local chromosome regions [26].

Next, we determined the genome-wide meiotic recombination landscapes of three asci (#1-#3) generated by QM6a and CBS999.97(*MAT1-1*) hybrid meiosis (Table 1 and Table S10). All 48 ascospores from these three asci could germinate and formed mycelia with dark-green conidial pigments, indicating that they are all euploid. The genomic DNA of the 48 F1 progeny was isolated and genotyped to detect *mat1-1, mat1-2* and *actin* as described previously [15]. The PCR results revealed that the 16 ascospores in each ascus can be classified into four genetically identical groups. One representative ascospore from each genetically identical group was then selected for whole genome sequencing using an Illumina-Miseq sequencer. We used PlotTetred, a component of the “ReCombine” programs [24], to create graphical representations of all seven chromosomes of QM6a and CBS999.97(*MAT1-1*), as well as the four representative progeny from each ascus (Fig. 3b and Fig. S7a-S7c). Remarkably, all the interhomolog recombination products in these three *T. reesei* asci (#1-#3) are “simple CO” or “simple NCO”, without any other genotype switches within 5 kb (Table 1 and Table S11-S12). This feature differs from budding yeast meiosis, which often generates COs with discontinuous GC, NCOs with two GC tracts, or even ambiguous recombination products that can arise from more than one pathway [24].

**Fig. 3.**
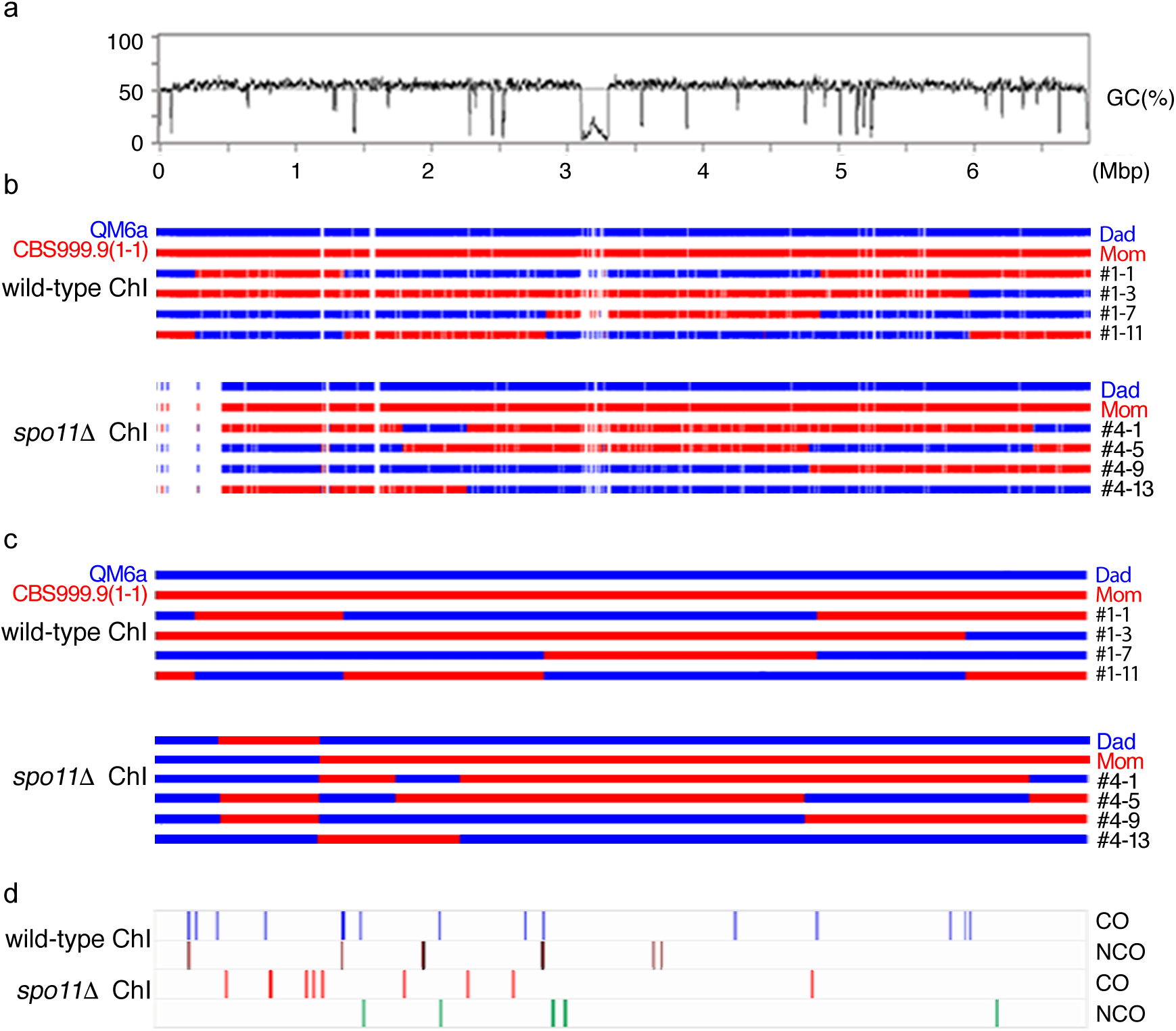
*Trichoderma reesei* meiosis generates interhomolog recombination products regardless of the presence or absence of *spo11*. (**a**) Traces represent graphs of GC content (window size 5000 bp) of the telomere-to-telomere sequence of the first QM6a chromosome (ChI). (**b**) The NGS short-read sequences identical to QM6a ChI are depicted by blue bars, those identical to CBS999.97(*MAT1-1*) ChI are represented by red bars. (**c**) The PacBio long-read sequences identical to QM6a ChI are depicted by blue bars, those identical to CBS999.97(*MAT1-1*) ChI are represented by red bars. (**d**) Overview of all ChI interhomolog recombination products detected in the three QM6a/CBS999.97(*MAT1-1*) asci (#1-#3) and the six QM6a *spo11*Δ/CBS999.97(*MAT1-1*) *spo11*Δ asci (#4-#9). The positions of COs and NCOs on each chromosome are indicated by vertical lines.

**Table 1.** Hybrid crossing of QM6a with CBS999.97(*MAT1-1*) generates both CO and NCO products regardless of the presence or absence of *spo11*

|  |  | QM6a<br>X<br>CBS99.97(*MAT1-1*) |  |  | QM6a *spo11*Δ<br>X<br>CBS99.97(*MAT1-1 spo11*Δ |  |  | QM6a *spo11*Δ<br>X<br>CBS99.97(*MAT1-1 spo11*Δ |  |  |
| --- | --- | --- | --- | --- | --- | --- | --- | --- | --- | --- |
| SNP coverage |  | 100% |  |  | 94.64% |  |  | ²50.51% |  |  |
| Ascus # |  | #1 | #2 | #3 | #4 | #5 | #6 | #7 | #8 | #9 |
| CO# |  | 24 | 24 | 27 | 19 | 12 | 21 | 5 | 7 | 19 |
| NCO# |  | 8 | 13 | 15 | 7 | 1 | 13 | 6 | 4 | 5 |
| Total |  | 75 COs & 36 NCOs |  |  | 52 COs & 21 NCOs |  |  | 31 COs & 15 NCOs |  |  |
| Normalized<br>interhomolog<br>recombination rate | CO1 | 1250 ± 86.6 cM¹ |  |  | 916 ± 250 cM |  |  | Not determined² |  |  |
|  | NCO | 600 ± 180 cM |  |  | 370 ± 317 cM |  |  |  |  |  |
|  | Both | 1850 ± 250 cM |  |  | 1286 ± 560 cM |  |  |  |  |  |
| Intragenic COs<br>& NCOs |  | 65 (59%) |  |  | 49 (47%) |  |  |  |  |  |
| Intergenic COs<br>& NCOs |  | 46 (41%) |  |  | 24 (33%) |  |  |  |  |  |
| COs & NCOs<br>at 5' or 3' regions³ |  | 5'-region |  | 3'-region | 5'-region |  | 3'-region | Not determined² |  |  |
|  |  | 40 (36%) |  | 42 (38%) | 17 (23%) |  | 42 (57%) |  |  |  |

1. One CO or NCO implies a genetic map length of 50 centiMorgans (cMs). 2. The sequence heterogeneity between the second pair of QM6a *spo11*Δ and CBS99.97(*MAT1-1*) *spo11*Δ strains is too low to accurately determine the interhomolog recombination rate. 3. The 5'-region is operationally defined from 1000 bp upstream and 1000bp downstream to the 'start' codon (ATG), whereas the 3'-region from 1000 bp upstream and 1000 bp downstream to the 'stop' codons, respectively.

Due to the short lengths of Illumina paired-end reads, we failed to accurately determine the recombination profiles of chromosomal regions hosting long AT-rich blocks (Fig. 3b and Fig. S7a-S7c). Accordingly, we applied PacBio SMRT technology to sequence and assemble the complete genomes of the four representative F1 progeny in ascus #1 (Table S13). By comparing the completed genome sequences of the two parental genomes, we were able to infer accurate recombination landscapes for all seven chromosomes (Fig. 3c-3d). These results not only confirm all interhomolog recombination products detected by the Illumina paired-end reads, but also reveal no new interhomolog recombination products.

We conclude that *Trichoderma reesei* Rad51 is capable of catalyzing interhomolog recombination to form both CO and NCO recombination products, though there is a high level of sequence heterogeneity between the genomes of QM6a and CBS999.97(*MAT1-1*). The average CO rate per meiosis is 1,250 ± 86.6 centiMorgans (cMs) or 28 ± 2 kb/cM (Table 1 and Table S14).

### *T. reesei* exhibits *Spo11*-independent interhomolog recombination

Next, we generated two pairs of QM6a *spo11*Δ and CBS999.97(*MAT1-1*) *spo11*Δ mutants by backcrossing the *spo11*Δ mutants to QM6a and CBS999.97(*MAT1-1*), respectively (Table S10). Based on SNP calling, we found that their genomes exhibited 94.64% and 58.32% sequence heterogeneity to each other. Next, we applied NGS to determine genome-wide meiotic recombination profiles of three representative asci (#4-#6) generated by the first *spo11*Δ mutant pair, as well as three representative asci (#7-#9) generated by the second *spo11*Δ mutant pair (Table 1, Table S14, Table S17-S18, Fig. 3b and Fig. S7). All six *spo11*Δ asci (#4-#9) we examined in this study produced both CO and NCO interhomolog recombination products.

We also applied PacBio SMRT technology to sequence and assemble the complete genomes of the four representative F1 progeny in ascus #4 (Fig. 3c, Fig. S8 and Table S13), and the results are consistent with those from whole-genome sequencing using an Illumina-MiSeq sequencer.

For the three asci (#4-6) generated from crossing the first pair of *spo11*Δ mutants, the average interhomolog CO rate per meiosis (after normalization to the percentage of overall sequence heterozygosity; 94.64%) is 916 ± 250 cMs (40 ± 13 kb/cM), which is ~70% that of the wild-type QM6a and wild-type CBS999.97(*MAT1-1*) cross (Table 1 and Table S14). Accordingly, we infer that the *spo11*-independent pathway is about two-fold more active than the *spo11*-dependent pathway.

This inference is consistent with our cytological observations that the *rad51*Δ asci exhibited stronger meiotic prophase arrest phenotypes than the *spo11*Δ *rad51*Δ asci. Presumably, the *rad51*Δ asci accumulate unrepaired ssDNAs generated from both Spo11-dependent DSBs and Spo11-independent DNA lesions, whereas the *spo11*Δ *rad51*Δ asci contain only Spo11-independent unrepaired ssDNA. We found that almost all *rad51*Δ asci exhibited a single strongly DAPI-stained focus (Figure 2B). The majority of *spo11*Δ *rad51*Δ asci (>90%) also contained one strongly DAPI-stained focus. Notably, most *spo11*Δ *rad51*Δ asci often (>50%) contained two or more additional micronuclei with no or few much weaker DAPI-stained signals (Fig. 2e and 2f).

### High fidelity of postmeiotic DNA replication in the presence or absence of *spo11*

To rule out the possibility that unexpected sequence variations or even interhomolog recombination products might be generated during the two rounds of postmeiotic mitosis, we isolated the genomic DNA from all 16 progeny generated from the #1 wild-type asci and the #4 *spo11*Δ asci for whole-genome sequencing using an Illumina-MiSeq sequencer (Table S10). The results of genome-wide sequence mapping confirm that all sixteen F1 progeny of these two asci can be classified into four genetically identical groups (Fig. S9). The nucleotide sequences of each genetically identical group possess very few mismatched sequences (≤1 per 10,000 bp) (Tables S19-S22). Moreover, there are no chromosomal abnormalities (deletions, inversions or translocations) in any of the 32 sexual progeny examined here. We infer that the two rounds of postmeiotic DNA replication are highly faithful, regardless of the presence or absence of *spo11*.

### *T. reesei spo11* is dispensable for chromosome synapsis during meiosis

In many sexual eukaryotes, homologous chromosome synapsis is mediated by the formation of synaptonemal complex (SC) during meiotic prophase. The SC is a tripartite proteinaceous structure consisting of two parallel lateral/axial filaments formed along sister chromatids of each homolog and numerous transverse filaments between two lateral elements forming the SC central element. Arrays of multiple chromatin loops are anchored to the lateral elements. The SC functions primarily as a scaffold to allow interacting chromatids of homologs to complete their homologous recombination activities. The SC transverse filament proteins of most sexual eukaryotes share no sequence similarity but strong structural homology; all comprise long internal coiled-coil domains (with sizes correlated with the width of the SC; ~100 nm) flanked by globular N- and C-terminal domains (see review in [27]). The formation of SC between homologs can occur independently of recombination in some organisms (e.g. *C. elegans* and *D. melanogaster*). It is also true that for many sexual eukaryotes, SC assembly is coupled to formation of recombination intermediates, such as *S. cereveisae, S. macrospora*, and mouse (see review in [27]). *T. reesei*, like *S. macrospora*, possesses the *sme4* homologous gene encoding a conserved traverse filament protein [28], but it is still unclear if SC occurs in *T. reesei* and whether *T. reesei spo11* is required for the formation of SC.

To investigate this topic, we applied transmission electron microscopy (TEM) to visualize SC in *T. reesei* meiotic cells at pachytene stage. In both wild-type (Fig. 4a) and *spo11*Δ (Fig. 4b), the two lateral elements lie about 100 nm apart and are interconnected by transverse filaments. Using TEM tomography, we were able to reconstruct several fully developed SCs in representative *spo11*Δ meiotic cells (Fig. 4d-4e and Movie S1). In contrast, only unsynapsed axial elements were detected in the *rad51*Δ asci (Fig. 4c, Fig. 4f and Movie S2). We conclude that assembly of SC is coupled to meiotic recombination in *T. reesei*. Notably, unlike *S. cerevisiae spo11* [3] and *S. marcospora spo11* [28], *T. reesei spo11* is dispensable for promoting meiotic chromosome synapsis.

**Fig. 4.**
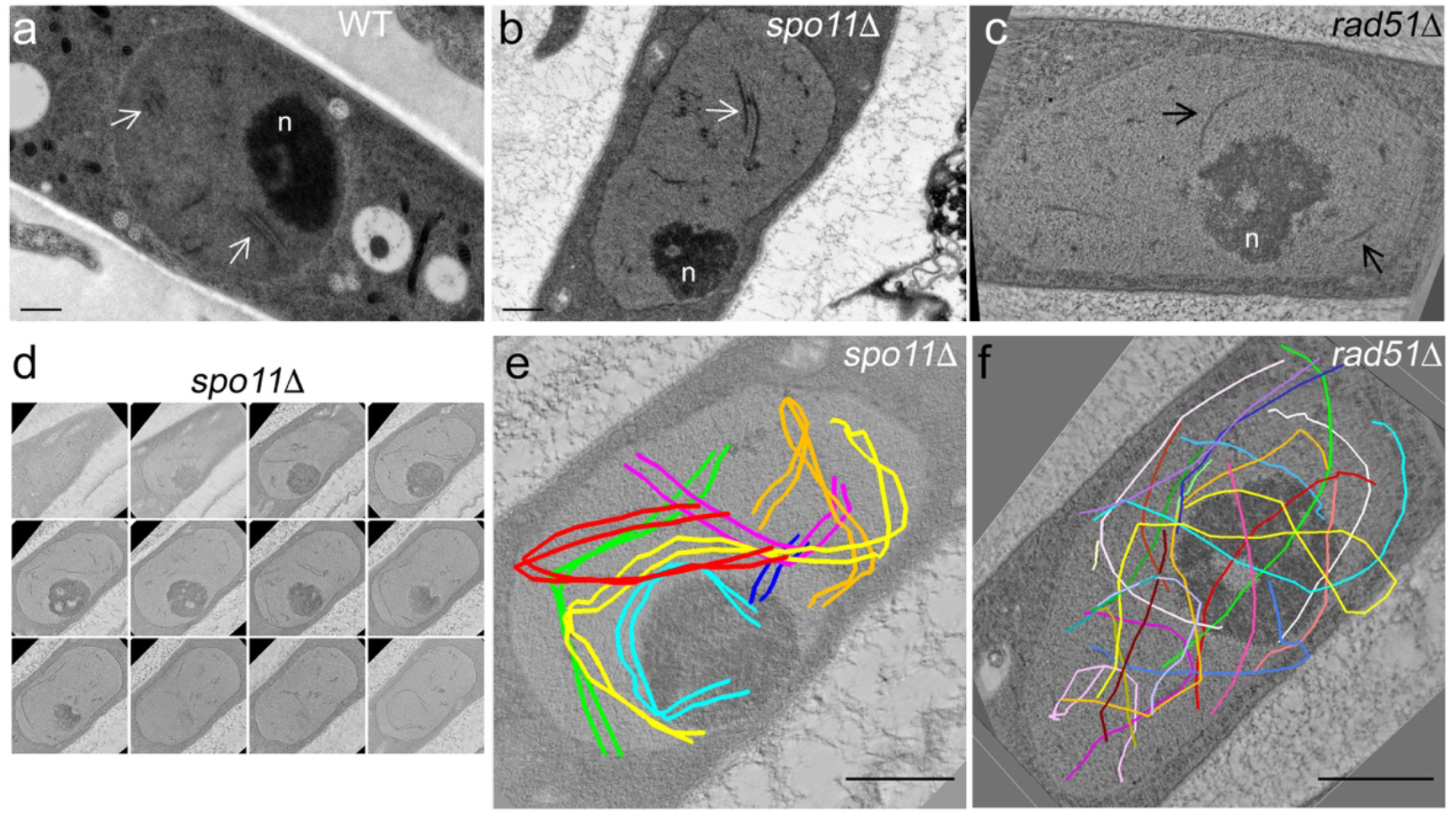
The synaptonemal complex (SC) in wild-type and *spo11*Δ and the axial elements in *rad51*Δ. Representative TEM images of wild-type (**a**), *spo11*Δ (**b**) and *rad51*Δ (**c**) meiotic cells at the pachytene stages. The SCs in wild-type and *spo11*Δ are marked by white arrows, whereas axial elements in *rad51*Δ are marked by black arrows. The enlarged nucleolus (n) is a hallmark of meiotic prophase nuclei. (**d**) Twelve sequential representative TEM images of a *spo11*Δ meiotic cell. (**e & f**) TEM tomography. The seven pairs of lateral elements in *spo11*Δ (**e**) and the fourteen axial elements in *rad51*Δ (**f**) are highlighted as colored lines. Black bar: 0.5 μm.

### Comparative analysis of COs and NCOs in the presence or absence of *spo11*

The three wild-type asci (#1-#3) generated 75 COs and 36 NCOs, whereas 83 COs and 36 NCOs were produced by the six *spo11*Δ asci (#4-#9) (Table 1 and Table S14). Using our nucleotide sequences, we could map their positions, and compare the distances between adjacent COs and/or NCOs, as well as their distances to neighboring AT-rich blocks. Remarkably, none of the interhomolog events in the nine asci examined here overlap (Fig. 3c and Table S23-S24).

Positive CO interference apparently exists in *T. reesei* regardless of the presence or absence of *spo11* (Fig. 5a). The inter-CO distances in wild-type and *spo11*Δ meiosis fit a gamma distribution [29]. The strength of interference is determined by the value of the shape parameter γ of the best-fit distribution (γ = 1 indicates a random distribution and γ > 1 indicates positive interference). We found that γ = 4.9 in wild-type and γ = 3.2 in *spo11*Δ meiosis.

**Fig. 5.**
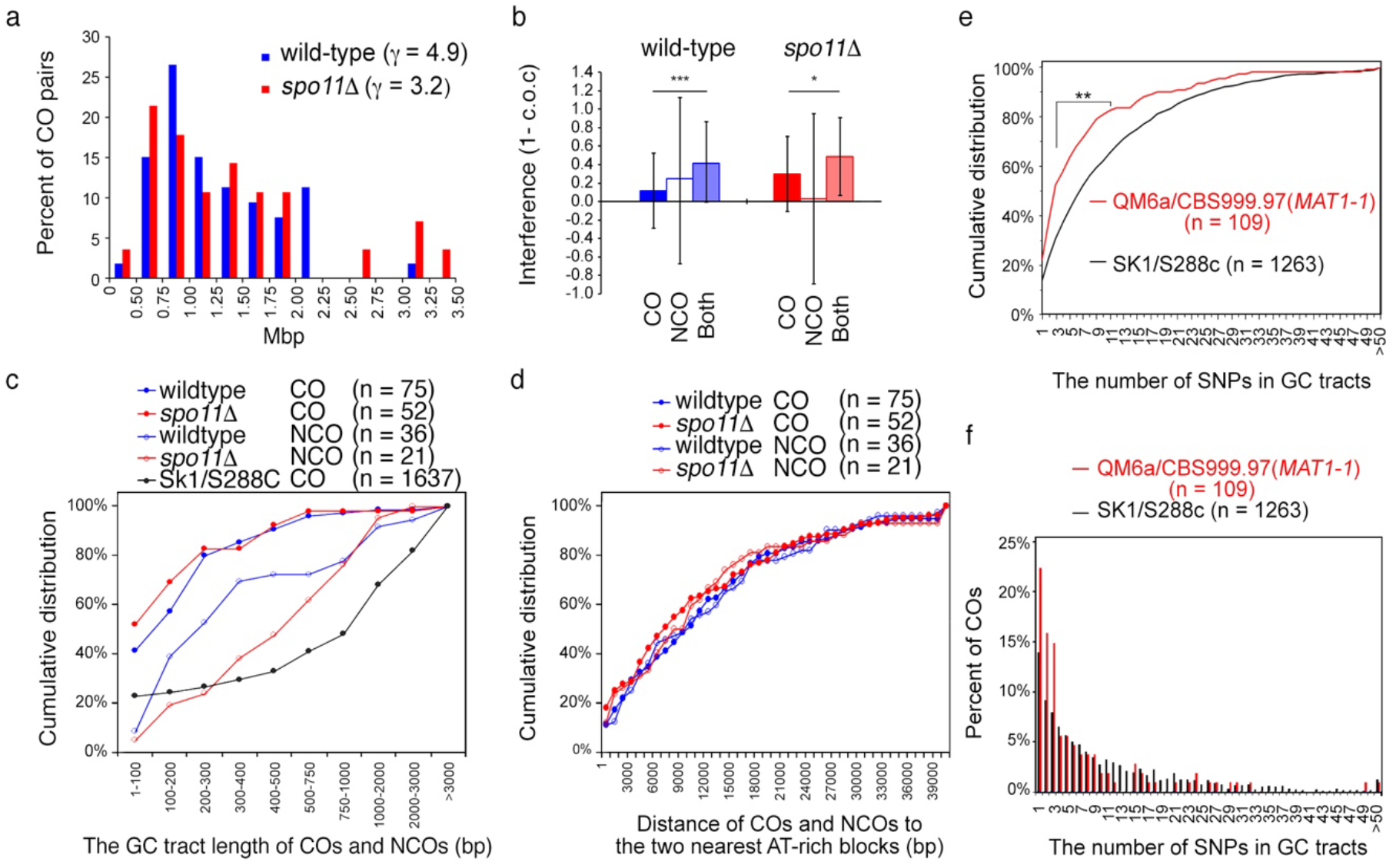
Comparative analysis of COs and NCOs in wild-type and *spo11*Δ asci. (**a**) Crossover interference. Distribution of distances between adjacent COs among the detected COs. Values for the shape parameter γ of the best-fit gamma distribution are indicated. (**b**) Interference between interhomolog recombination products. Interference calculated as 1 - c.o.c. (coefficient of coincidence) for COs only, NCOs only, and all events combined from whole-genome recombination profiling data. (**c**) Cumulative distribution of the lengths of COs and/or NCOs during hybrid meiosis of QM6a/CBS999.97(*MAT1-1*), QM6a *spo11*Δ/CBS999.97(*MAT1-1*) *spo11*Δ, and SK1/S288c. (**d**) Cumulative distribution of all COs and NCOs to the two adjacent AT-rich blocks. The NGS results of 20 different SK1/S288c tetrads published previously were reanalyzed here. (**e-f**) *T. reesei* Rad51, like budding yeast Dmc1, can tolerate mismatch sequences during the strand exchange reaction. (**e**) Cumulative distribution of the SNP number in the CO-associated gene conversion tracts. Results from nine QM6a/CBS999.97(*MAT1-1*) asci (#1-#9) and twenty SK1/S288c tetrads (Callender et al., 2016) are shown. (f) The two-sample Kolmogorov-Smirnov test (two-sided) confirmed that the two cumulative distributions are different (** *p* < 0.01). Relative distribution of SNP number in the CO-associated gene conversion tracts.

Previous study of budding yeast meiosis revealed that the genome-wide distribution of detectable interhomolog events (COs + NCOs) reflects the underlying Spo11-induced DSB distribution [30]. Moreover, DSB interference among NCOs is weaker than that among COs [31]. We found that, regardless of the presence or absence of *T. reesei spo11*, the distributions of COs, NCOs and a combination of interhomolog recombination products all display positive interference (Tables S25-S26). Unexpectedly, genetic interference among NCOs is stronger than that among COs for the wild-type, but weaker for *spo11*Δ (Fig. 5b; p < 0.01). We hypothesize that the spacing between Spo11-induced DSBs and/or Spo11-independent DNA lesions in *T. reesei* are tightly regulated by a novel interference mechanism(s), and that this mechanism(s) might be different from that in budding yeast.

GC tracts associated with interhomolog recombination events have been used for CO and NCO length measurements [32]. Compared to the median CO length (~1 kb) of SK1/S228c hybrid meiosis (Table S9), both *T. reesei* wild-type and *spo11*Δ meiosis generated COs with much shorter median lengths (<200 bp). The median lengths (400-500 bp) of NCOs in *T. reesei spo11*Δ are slightly longer than those (200-300 bp) of NCOs in wild-type (Fig. 5c and Table S11-S18). Since the longer GC tracts in budding yeast might be due to clustered Spo11-induced DSBs within the same chromosomal loops [26], we infer that both Spo11-induced DSBs and Spo11-independent DNA lesions in *T. reesei* might be tightly controlled to prevent clustering.

We also found that the median SNP number per CO in QM6a/CBS999.97(*MAT1-1*) hybrid meiosis is ~3.5, which is significantly lower than that (~6.0) in SK1/S288c hybrid meiosis (Fig. 5e). Intriguingly, in both QM6a/CBS999.97(*MAT1-1*) and SK1/S288c hybrid meiosis, there are few long GC tracts containing a high number (up to 50) of SNPs (Fig. 5f). These results support the notion that *T. reesei* Rad51, like *S. cerevisiae* and mammalian Dmc1 (but not Rad51), can tolerate mismatched sequences during the strand exchange reaction.

### Interhomolog recombination products tend to be located at 3′-regions of protein-encoding genes in the *spo11*Δ mutant line

Previous studies revealed that meiotic DSBs are not randomly distributed along chromosomes. In *S. cerevisiae*, Spo11-induced DSBs form preferentially on chromosome arms, and preferentially form in GC-rich chromatin loop regions rather than AT-rich axis-associated DNA [33]. Locally, there are DSB hotspots (typically ~200 bp) in which Spo11 cleaves preferentially. The majority (88%) of DSB hotspots in *S. cerevisiae* are within nucleosome-depleted regions (NDRs) in gene promoters [34–36].

Our results revealed that, at the chromosome level, the majority of interhomolog events in *T. reesei* preferentially occur within chromosomal loops rather than at the chromosomal axes. Of the 127 CO and 57 NCO products we detected in the six asci (#1-#6) (Table 1), only 7 COs and 5 NCOs overlap with short AT-rich blocks (either 500 or 1000 bp in length; Tables S23-S24). Therefore, both Spo11-induced DSBs and Spo11-independent DNA lesions preferentially occur within chromosomal loops in *T. reesei*. No interhomolog recombination product was detected within centromeres, telomeres or subtelomeric regions, with these long AT-rich blocks perhaps forming constitutive heterochromatin and displaying a reduced frequency of meiotic recombination.

Locally, regardless of the presence or absence of *spo11*, two-thirds of the GC tracts associated with interhomolog recombination products overlap with intragenic regions, whereas one-third overlap with intergenic regions (Table 1 and Table S27). In the presence of *spo11*, there is no bias in the occurrence of interhomolog recombination products at either the 5′ or 3′ regions of protein-encoding genes. The 5′ region is operationally defined as being from 1000 bp upstream and 1000 bp downstream of the ‘start’ codon (ATG), whereas the 3′ region is from 1000 bp upstream and 1000 bp downstream to the ‘stop’ codons. Remarkably, in *spo11*Δ, >57% of interhomolog recombination products overlap with 3′ regions and only ~ 23% overlap with 5′ regions (Table 1).

### Top2 might be responsible for Spo11-independent DSBs during *T. reesei* meiosis

Three lines of evidence suggest that Top2 likely acts redundantly with Spo11 to initiate interhomolog recombination in *T. reesei*. First, *T. reesei sae2* is required for both *spo11*-dependent and *spo11*-independent interhomolog recombination in *T. reesei*. Sae2 and the Mre11-Rad50-Nbs1/Xrs2 complex not only remove the 5′ protein adducts of Spo11 during initiation of meiotic recombination [3], they also function in Top2 removal from the 5′ termini that arise during poisoned topoisomerase reactions [37–39]; Second, the twin-supercoiled-domain model [40] can explain why the majority of Spo11-independent interhomolog products overlap with the 3′ regions of protein-encoding genes. During transcription, RNA polymerase (RNAP) tracks the helical groove of DNA, overtwisting downstream DNA and undertwisting upstream DNA. Therefore, positively supercoiled DNA is generated in front of RNAP and negatively supercoiled DNA forms behind it. Like *S. cerevisiae* [41] and *Drosophila* [42], *T. reesei* has only one isoform of Top2 (this study). Humans express two closely related isoforms, Top2α and Top2β. It has been reported previously that *S. cerevisiae* Top2, *Drosophila* Top2 and human Top2α preferentially relaxes positively supercoiled DNA [43]. These Top2 proteins likely function ahead of RNAP. Consistent with this supposition, a genome-wide mapping study has revealed that human Top2α cleavage sites preferentially cluster in more distal regions of protein-encoding and/or non-coding RNA genes [44]. Third, we found that addition of idarubicin, an effective inhibitor of fungal Top2 [45], to the developing stromata could partly rescue the meiotic prophase arrest phenotype of both *rad51*Δ (Fig. 2g and 2h) and *rad51*Δ *spo11*Δ asci (Fig. 2i and 2j).

## Discussion

Meiotic recombination promotes proper genetic diversity and chromosome segregation. Accurate estimates of recombination rates are of great importance for our understanding of the molecular mechanisms governing meiosis and evolution. A milestone in this respect is the development of NGS-based analyses of hybrid *S. cerevisiae* genomes bearing thousands of heterozygous SNP markers (see review in [46]). Here, in this study, we combined both NGS and PacBio RSII sequencing technology to whole genome sequence data from hybrid crosses of two highly polymorphic *T. reesei* strains, QM6a and CBS999.97(*MAT1-1*). Our recombination maps of *T. reesei* hybrid meiosis provide novel insights into the mechanisms of meiotic recombination in this economically important fungus.

### *T. reesei* is an ideal model for studying Rad51-only meiosis

The first key finding of this study is that *T. reesei* Rad51 is essential and sufficient for promoting interhomolog recombination and chromosome synapsis in this Rad51-only eukaryote. Our genome-wide recombination mapping data indicate that *T. reesei* Rad51 can tolerate mismatched sequences in the strand exchange reaction of a highly polymorphic hybrid meiosis. This property of *T. reesei* Rad51 is similar to that of Dmc1 (but not Rad51) in budding yeast and mammals [10–12].

It was reported previously that the eukaryotic *rad51* and *dmc1* genes formed two separate monophyletic groups when archaeal RecA-like recombinase genes (*RADA*) were used as an outgroup. The *dmc1* genes were lost independently in Rad51-only eukaryotic organisms during evolution. Moreover, *D. melanogaster* and *C. elegans* possess rapidly evolving *rad51* genes [47]. Rapid evolution might be a general property of all *rad51* genes since they are typically non-interchangeable within genera and even within species. For example, fission yeast, mouse and human *rad51* genes all failed to complement the DNA repair defect of *S. cerevisiae rad51* mutants [48]. Further studies are needed to identify the structural motif(s) in Dmc1 and *T. reesei* Rad51 responsible for tolerating mismatched sequences during meiotic recombination. It will also be of great interest to determine if and how the *rad51* genes in “Rad51-only” eukaryotes independently acquired the capability to tolerate mismatched sequences after (or before) their *dmc1* genes were lost.

### *spo11*-independent interhomolog recombination prevails during *T. reesei meiosis*

The second important finding of our study is that removal of *T. reesei spo11* does not affect the formation of SC and mature asci with 16 viable ascospores. On average, the frequency of Spo11-independent interhomolog recombination is about two-fold greater than that of Spo11-induced interhomolog recombination. Both Sae2 and Rad51 are required for repair of Spo11-induced and Spo11-independent DSBs. Accordingly, *T. reesei spo11*Δ mutants represent the best model yet for studying *spo11*-independent meiotic recombination for a number of reasons. First, all other studied sexual eukaryotes (except social amoebae) require their *spo11* genes to generate sufficient CO products between homologous chromosomes (reviewed in [8, 49–51]). Second, compared to *T. reesei*, sexual reproduction was rarely observed in the best-studied model organisms of social amoebae, *D. discoideum* [52]. Third, advancements in genetic transformation tools (reviewed in [13]) and the availability of *T. reesei* haploid genome sequences ([53] and this study) make *T. reesei* amenable to multiple experimental approaches, including genetics, reverse genetics and genomics.

### Does Top2 induce Spo11-independent DSBs?

We present three lines of evidence in this study to support the notion that Top2 might act redundantly (and predominantly) to initiate meiotic recombination in *T. reesei*. Top2 has the capability to make DSBs. Top2-dependent post-meiotic DSBs in *Tetrahymena thermophila* occur in conjunction with the transition from a heterochromatic to euchromatic chromatin structure in the haploid pronucleus [54]. In budding yeast, inactivation of the DNA damage checkpoint kinase Mec1, a homolog of mammalian ataxia telangiectasia and Rad3-related (ATR) protein kinase, leads to Top2-dependent DSBs at fragile sites (referred to as replication slow zones, RSZs). RSZs are homologous to mammalian common fragile sites (CFSs), stability of which is regulated by ATR [55, 56]. In human K562 leukemia cells, Top2α cleaves functionally conserved local sequences at cleavage cluster regions (CCRs). Top2α CCRs are biased toward the distal regions of gene bodies [44]. Consistent with this scenario, we found that CO and NCO products arising from *spo11*Δ meiosis were biased toward the 3’ region of protein-encoding genes. Genome-wide detection of Spo11-independent DSB sites will reveal whether they share conserved local sequences with yeast RSZs and mammalian CFSs or CCRs. Further investigations are also needed to determine whether Top2 is involved in the initiation of Spo11-independent meiotic DSBs or recombination in other sexual eukaryotes.

### Genetic interference and crossover homeostasis

Genetic interference is a common characteristic of sexual eukaryotes, with very few known exceptions (e.g., fission yeast) in which recombination proceeds with no interference (see review of [57]). In this study, we show that both CO and NCO products proceed with positive interference regardless of the presence or absence of *spo11*. In addition, *T. reesei* preferentially produces “simple” CO and “simple” NCO products (Table 1 and Table S11-S12). Both Spo11-induced and Spo11-independent DSBs are tightly regulated to prevent clustering within the same or even nearby chromosome loops.

Our data also reveal that loss of *spo11* resulted in a higher CO/NCO ratio, i.e., 2.1 in wild-type (asci #1-#3) and 2.5 in *spo11*Δ (asci #4-#6) *T. reesei* (Table 1). One potential explanation for this outcome is that a buffer system operates in *T. reesei* meiosis to maintain CO levels at the expense of NCOs when there are no Spo11-dependent DSBs. This buffer system is functionally analogous to “CO homeostasis”, which ensures a stable CO number at the expense of NCOs even if the number of Spo11-induced DSBs varies. The “CO homeostasis” phenomenon was first discovered in *S. cerevisiae* meiosis [58] and subsequently found in *D. melanogaster, C. elegans* and mouse meiosis (reviewed in [59]). A second potential explanation is simply that Spo11-independent DSBs have a greater tendency to generate CO products as compared to Spo11-dependent DSBs. It will be important to further investigate these two hypotheses and decipher the underlying molecular mechanism. The findings of that investigative effort will also provide new insights into whether the CO/NCO decision is made before or after initiation of DSBs, given that both Spo11-independent DSBs and Spo11-induced DSBs share the same molecular mechanism for DSB end processing (i.e., Sae2) and Rad51-only DSB repair.

### SC assembly in *T. reesei* is Spo11-independent but requires Rad51

*T. reesei* possesses the gene homologous to the S. marcospora *sme4* encoding a conserved traverse filament protein [28]. We show for the first time that the seven pairs of homologous chromosomes in *T. reesei* wild-type and *spo11*Δ meiotic cells form SCs. In contrast, the *rad51*Δ mutant only forms unsynapsed axial elements. Our results confirm that the *T. reesei* SC shares strong structural homology with that of other sexual eukaryotes and that the assembly of SC in *T. reesei* is coupled to meiotic recombination regardless of the presence or absence of Spo11. It is currently thought that the function of the SC is to shape the chromosome-wide landscape of interhomolog recombination products via cooperative assembly, CO promotion and NCO inhibition. Further analyses of corresponding *T. reesei* mutants will provide insights into the functional interactions between SC assembly, recombinosome localization, genetic interference and CO homeostasis.

## Conclusion

In conclusion, *T. reesei* represents an excellent model organism for studying Spo11-independent (or Top2-dependent) and Rad51-only meiosis. The three *T. reesei* genome sequences, we report here, are the highest quality yet generated and provide a better framework for genetic and genomic analyses of this industrially important fungus. Further investigations of the underlying mechanisms of *T. reesei* meiosis can provide new information on the molecular mechanism(s) and evolution of meiosis, as well as revealing innovative avenues for industrial strain improvements.

## Methods

### Miscellaneous

Fungal growth, culture media, sexual crossing, single ascospore isolation, preparation of genomic DNA, PCR genotyping, Southern hybridization, DAPI staining and cytological analysis have been described in our previous studies [15, 60]. Whole genome sequencing and assembly were carried out by the PacBio Single Molecule Real-Time (SMRT) method and the Illumina-MiSeq paired-end method [16]. Asci were stained with DAPI (4’,6-diamidino-2-phenylindole) and visualized by the DeltaVision Core Imaging System (Applied Precision, LLC, USA).

### Pulsed field gel electrophoresis (PFGE)

Intact chromosomal DNA was prepared using the agarose spheroplasting method with two modifications. The conidia were isolated and germinated in potato dextrose agar (PDA) at 25 °C, 200 rpm for 6 hr before being embedded into the agarose plug. *Trichoderma harziaum* lysing enzymes (Sigma-Aldrich Co., St. Louis, MO) were used to digest the cell walls during spheroplasting. Electrophoresis conditions for karyotype analysis were described previously [61].

### SNP genotyping and recombination analysis

All bioinformatics experiments were performed using the same approaches described previously for recombination analysis of the SK1/S288c tetrads [12]. The complete high quality genome sequences of QM6a [16], CBS999.97(*MAT1-1*) and CBS999.97(*MAT1-2*) were used as the references for SNP calling. For each sample, either the PacBio-SMRT long reads or the Illumina paired-end reads were mapped to the reference genomes using Burrows-Wheeler Aligner (BWA v0.6.2) [62]. The recombination events were detected with an inhouse pipeline incorporating a similar algorithm to the CrossOver (v6.3) algorithm in the ReCombine programs (v2.1) [24]. Genotype data were formatted according to Anderson et al. (2011) and the program was run with a 0 bp threshold. The output data were then processed using the GroupEvents program to merge closely spaced events into single classes [25]. To validate the efficacy of our bioinformatics methods, we used them to reanalyze the NGS data of two previously published SK1/S288c tetrads [12]. The results confirmed that we could identify almost all previously reported recombination products (Table S9).

### TEM tomography

Developing fruiting bodies were collected and dissected into thin sections (0.2 mm thick) using a Vibratome 1000 PLUS. Pre-fixation was performed in 2.5% glutaraldehyde/0.1M sodium cacodylate (pH 7.2) for 3 hr at room temperature, and then shifted to 4 °C for overnight. The specimens were washed three times in 0.1M sodium cacodylate (pH 7.2) for 15 min in each change. Post-fixation was performed by 2% OsO4 solution/0.1M sodium cacodylate buffer at room temperature for 1 hr. Next, the specimens were washed three times with 0.1M sodium cacodylate for 15 minutes. We used 2% uranyl acetate for in block-staining at room temperature for 1 hr. For dehydration, a series of ethanol solutions with 10% increments (from 10% to 100%) were used. The specimens were immersed twice into 100% 1,2-propylene oxide (PO) for 20 minutes. Infiltration was performed for 4 hr in a mixture of PO with 10% Spurr’s resin, and we then increased the concentration of Spurr’s resin by 10% for 8-12 hr each change till 100%. The specimens were immersed twice every 12 hours into the Spurr’s resin with an accelerator (DMAE) for 12 hours, and then transferred to the embedding molds. Polymerization proceeded at 70 °C for 12 hr. Serial 200 nm-thick sections were collected and scanned with a Thermo Scientific™ Talos L120C TEM system (4×4k CMOS Ceta camera) operating at 120 kV with 0 and 90 degrees. The TEM tomography images were taken at 6700× magnification. We used the Inspect 3D 4.3 program for image stack alignment, reconstruction and axis registration for each section. The Amira Software 6.0 was used to align and concatenate all image sections. Three-dimensional models were generated by inputting the concatenated images into Imaris9.1.2 software.

## Supporting information

Movie 1 (WCL-TFW)

Movie 2 (WCL-TFW)

## Data access

The complete genome sequences of QM6a [16], CBS999.97(*MAT1-1*) and CBS999.97(*MAT1-2*) have been submitted to the National Center for Biotechnology Information (NCBI), and the accession numbers are CP017983-CP017984, CP020875-CP020879, CP020724-CP020730, CP040187-CP040228 and SRR6884626-SRR6884713, respectively. The raw dataset of the PacBio long reads and Illumina-Miseq paired-end reads from this study have been submitted to the Genomes (WGS) and Sequence Read Archive (SRA) of NCBI, and the accession numbers are BioProjects: PRJNA352653, PRJNA382020, PRJNA386077 and PRJNA433292 (Table S10).

## Acknowledgements

TFW was supported in this work by Academia Sinica (AS-105-TP-B07 and AS108-TP-B07) and the Ministry of Science and Technology (MOST163-2311-B-001-016-MY3), Taiwan, Republic of China. CLW, YCC and YJC were supported by postdoctoral fellowships from Academia Sinica. LT was supported by a visiting specialist fellowship from Academia Sinica. We thank Shu-Yun Tung (IMB Genomic Core) for NGS sequencing service, Kun-Hai Ye and Yi-Ning Chen (IMB Bioinformatics Core) for statistical help and bioinformatics consulting service, John O’Brien for English editing, Chin-Fu Cheng and John Kung (IMB Animal Facility) for programming help with the in-house bioinformatics pipeline, Sue-Ping Li (IMB, Imaging Core) for help with TEM tomography, and Yu-Tang Huang (IMB Computer Room) for maintaining the computer workstation.

## Authors’ contributions

WCL, YCC, CLC, CLW, LT, WLP and YJC performed the experiments and analyzed the data. WCL and TFW conceived and designed the experiments and wrote the paper. All of the authors read and approved the manuscript.

