## Supplemental Tables and Figures for "Two different pathways for initiation of *Trichoderma reesei* Rad51-only meiotic recombination"

#### List of supplemental Tables and Figures

| File name | Title | Page # |
|---|---|---|
| Supplemental_Fig_S1 | Karyotyping by pulsed field gel electrophoresis (PFGE). | 1 |
| Supplemental_Fig_S2 | Phylogenetic tree of Rad51 and Dmc1 in 15 different eukaryotic organisms. | 2 |
| Supplemental_Fig_S3 | The primary structures of Spo11 proteins in six fungi. | 3 |
| Supplemental_Fig_S4 | Rad51 homolog sequence alignment. | 4 |
| Supplemental_Fig_S5 | Sae2 homolog sequence alignment. | 5 |
| Supplemental_Fig_S6 | Generation of *T. reesei* mutant strains | 6 |
| Supplemental_Fig_S7 | NGS analyses of *T. reesei* meiotic interhomolog recombination products | 7 |
| Supplemental_Fig_S8 | *Trichoderma reesei* meiosis generates interhomolog recombination products regardless of the presence or absence of *spo11* | 8 |
| Supplemental_Fig_S9 | High fidelity of the two rounds of postmeiotic DNA replication regardless of the presence or absence of *spo11*. | 9 |

| | | |
|---|---|---|
| Supplemental_Table_S1 | Summary of PacBio RSII sequencing and assembly results. | 10 |
| Supplemental_Table_S2 | Characteristics and assembly of the seven chromosomes in CBS999.97(*MAT1-1*) and CBS999.97(*MAT1-2*) | 11-12 |
| Supplemental_Table_S3 | Transposable elements in QM6a, CBS999.97(*MAT1-1*) and CBS999.97(*MAT1-2*). | 13 |
| Supplemental_Table_S4 | Size distribution of AT-rich blocks in QM6a, CBS999.97(*MAT1-1*) and CBS999.97(*MAT1-2*) | 14 |
| Supplemental_Table_S5 | SNPs between CBS999.97(*MAT1-1*) and CBS999.97(*MAT1-2*) | 15 |
| Supplemental_Table_S6 | SNPs between QM6a and CBS999.97(*MAT1-1*) | 16 |
| Supplemental_Table_S7 | SNPs between QM6a and CBS999.97(*MAT1-2*) | 17 |
| Supplemental_Table_S8 | *T. reesei spo11* is dispensable for meiosis and meiosis-driven segmental aneuploidy (SAN) | 18 |
| Supplemental_Table_S9 | Summary of COs and NCOs (with ≥ 2 SNP) in two representative SK1/S288c tetrads | 19-24 |
| Supplemental_Table_S10 | List of all *Trichoderma reesei* strains analyzed by whole genome sequencing technology | 25-36 |

| | | |
|---|---|---|
| Supplemental_Table_S11 | Summary of all CO products from the three asci (#1-#3) generated by crossing QM6a and CBS999.97(*MAT1-1*) | 37-40 |
| Supplemental_Table_S12 | Summary of all NCO products from the three asci (#1-#3) generated by crossing QM6a and CBS999.97(*MAT1-1*) | 41-42 |
| Supplemental_Table_S13 | PacBio RSII sequencing and assembly results of the four representative F1 progeny in the #1 ascus and the #4 ascus | 43-45 |
| Supplemental_Table_S14 | Distribution of CO and NCO products in the seven homologous chromosomes | 46 |
| Supplemental_Table_S15 | Summary of all CO products from the three asci (#4-#6) generated by crossing the first pair of QM6a *spo11*Δ and CBS999.97(*MAT1-1*) *spo11*Δ mutants | 47-49 |
| Supplemental_Table_S16 | Summary of all NCO products from the three asci (#4-#6) generated by crossing the first pair of QM6a *spo11*Δ and CBS999.97(*MAT1-1*) *spo11*Δ mutants | 50-51 |
| Supplemental_Table_S17 | Summary of all CO products from the three asci (#7-#9) generated by crossing the second pair of QM6a *spo11*Δ and CBS999.97(*MAT1-1*) *spo11*Δ mutants | 52-53 |
| Supplemental_Table_S18 | Summary of all NCO products from the three asci (#7-#9) generated by crossing the second pair of QM6a *spo11*Δ and CBS999.97(*MAT1-1*) *spo11*Δ mutants | 54 |
| Supplemental_Table_S19 | Summary of pairwise SNP calling between the 5th chromosomes in the 16 F1 progeny generated by QM6a and CBS999.97(*MAT1-1*) ascus #1 | 55 |
| Supplemental_Table_S20 | Summary of pairwise SNP calling between the 6th chromosomes in the 16 F1 progeny generated by QM6a and CBS999.97(*MAT1-1*) ascus #1 | 56 |

| | | |
|---|---|---|
| Supplemental_Table_S21 | Summary of pairwise SNP calling between the 5th chromosomes in all 16 F1 progeny of the #4 ascus generated by crossing QM6a *spo11*Δ and CBS999.97(*MAT1-1*) *spo11*Δ | 57 |
| Supplemental_Table_S22 | Summary of pairwise SNP calling between the 6th chromosomes in all 16 F1 progeny of the #4 ascus generated by mating QM6a *spo11*Δ and CBS999.97(*MAT1-1*) *spo11*Δ | 58 |
| Supplemental_Table_S23 | Distances in the 6 asci for all COs to the two neighboring AT-rich blocks | 59-65 |
| Supplemental_Table_S24 | Distances in the 6 asci for all NCOs to their two neighboring AT-rich blocks | 66-68 |
| Supplemental_Table_S25 | The coefficient of coincidence and interference of all neighboring interhomolog products in the three asci (#1-#3) generated by crossing QM6a and CBS999.97(*MAT1-1*) | 69-73 |
| Supplemental_Table_S26 | Coefficient of coincidence and interference for all neighboring interhomolog products (COs and NCOs) in the three asci (#4-#6) generated by crossing the first pair of QM6a *spo11* Δ and CBS999.97(*MAT1-1*) *spo11*Δ mutants | 74-76 |
| Supplemental_Table_S27 | Locations of interhomolog recombination products in the six asci (#1-#6). | 77-85 |

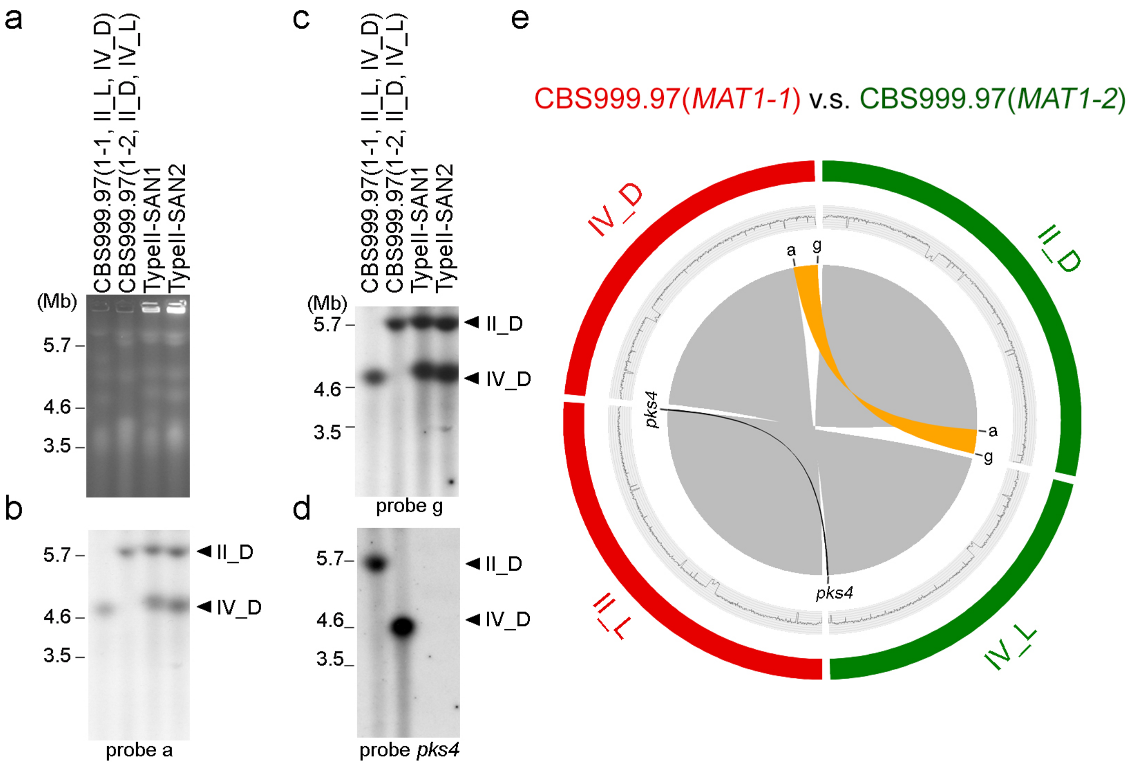

**Fig. S1. Karyotyping by pulsed field gel electrophoresis (PFGE). (a)** Electrophoretic karyotyping and (**b-d**) Southern hybridization of CBS999.97(*MAT1-1*, II_L, IV_D), CBS999.97(*MAT1-2*, II_D, IV_L) and the two SAN progeny generated by crossing CBS999.97(*MAT1-1*, II_L, IV_D) and CBS999.97(*MAT1-2*, II_D, IV_L). (**e**) The locations of the three DNA probes (a, g, *pks4*) for Southern hybridization. The D segment is indicated in orange, and the L segment in black. The outer circles indicate the two rearranged chromosomes (II and IV) in CBS999.97(*MAT1-1*) (in red) and CBS999.97(*MAT1-2*) (in green). GC contents (window size 5000 bp) of the seven chromosomes are shown in the middle traces.

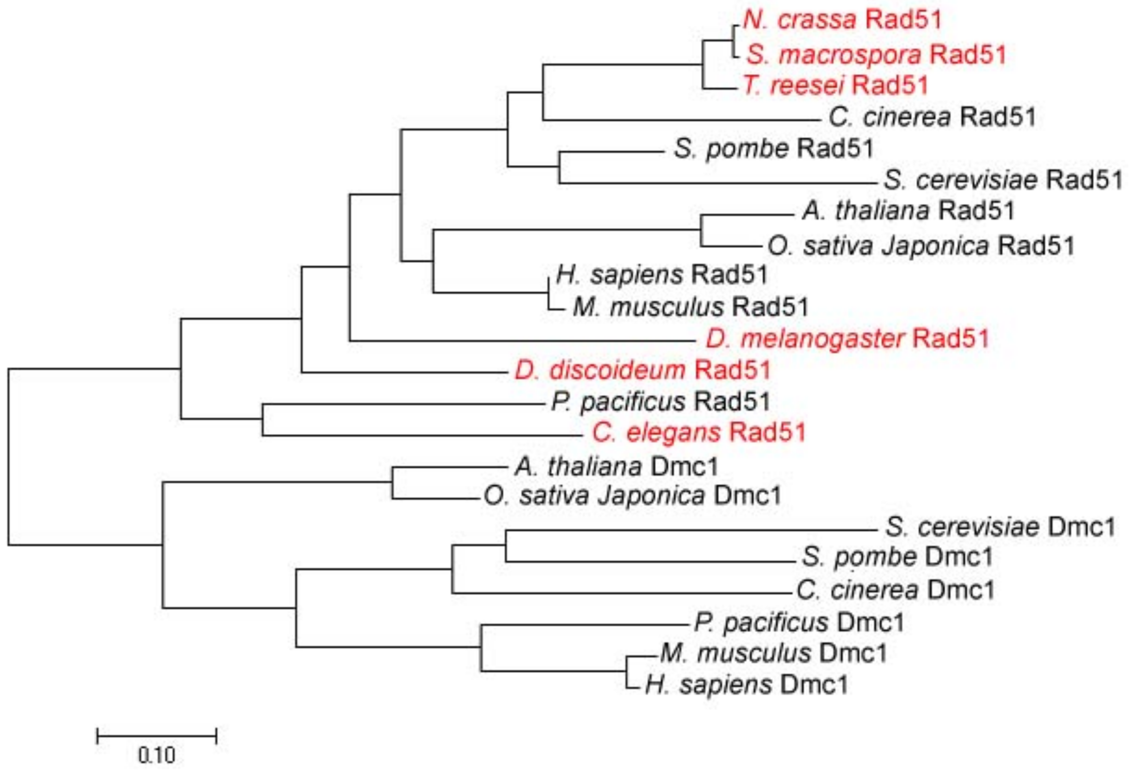

**Fig. S2. Phylogenetic tree of Rad51 and Dmc1 in 15 different eukaryotic organisms.** Dmc1 has been lost from six Rad51-only eukaryotic organisms (in red). The evolutionary history was inferred by using a Maximum Likelihood approach based on the JTT matrix-based model. The tree is drawn to scale, with branch lengths equating to the number of substitutions per site. The analysis involved 20 sequences of 323 amino acids. All positions containing gaps and missing data were eliminated. Evolutionary analyses were conducted in MEGA7.

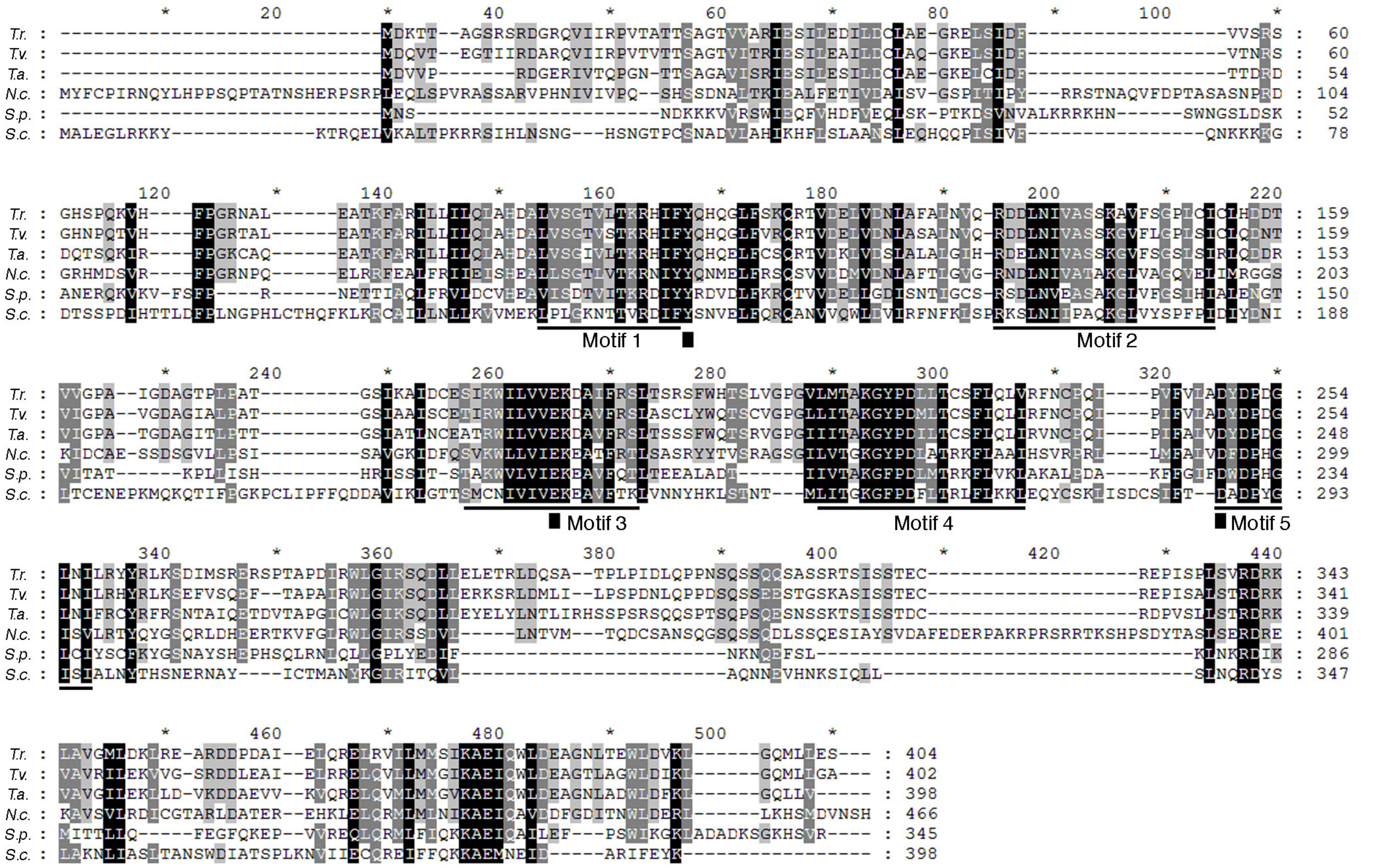

**Fig. S3. The primary structures of Spo11 proteins in six fungi.** The amino acid sequences of Spo11 protein sequences of *Trichoderma reesei* (*T.r.*), *Trichoderma atroviride* (*T.a.*) and *Trichoderma virens* (*T.v.*) were aligned with those of *Neurospora crassa* (*N.c.*), *Schizosaccharomyces pombe* (*S.p.*) and *Saccharomyces cerevisiae* (*S.c.*). Five conserved motif regions are denoted by black lines below the sequence blocks. The three amino acids (Tyr-135, Glu-233 and Asp-288) essential for the initiation of meiotic recombination in *Saccharomyces cerevisiae* Spo11 are indicated by black squares.

```
                    *        20         *        40         *        60         *        80         *       100         *
T.r. : ----------------------------M--------------------------------SDEYDEENQVAEEGGMTGPGAPTPLSALEGVAGLTKRDIQLVVDGGFNT :  50
N.c. : ----------------------------MSN------------------------------NEEYDESANMEDDSGIPNPGAPTPLSALEGVAGLTKRDIQLIVDGGFNT :  52
S.p. : ---------------------MADTEVEMQVSAADMNNN-------------ENGQAQSNYEYDVNVQDEEDEAA-----AGPMPLQMLEG-NGITASDIKKIHEAGYYT :  70
H.s. : ---------------------MA-----MQM------------------------------QLEANADTSVEEES-----FGPQPISRLEQ-CGINANDVKKLEEAGFHT :  48
S.c. : MSQVQEQHISESQLQYGNGSLMSTVPADLSQSVVDGNGNGSSEDIEATNGSGDGGGLQEQAEAQGEMEDEAYDEAAL---GSFVPIEKLQV-NGITMADVKKLRESGLHT : 106

                  120         *       140         *       160         *       180         *       200         *       220
T.r. : VESVAYTPRRVLEQIKGISEQKATKILAEASKLVPMGFTTATEMHQRRSELISITTGSKNLDTLLAGGIETGSVTELFGEFRTGKSQICHTLAVTCQLPFDMGGGEGKCL : 160
N.c. : VESVAYTPRRVLEQIKGISEQKAGKILAEASKLVPMGFTTATEMHQRRSELISITTGSKNLDTLLAGGIETGSVTEIFGEFRTGKSQICHTLAVTCQLPFDMGGGEGKCL : 162
S.p. : VESIAYTPKRQLLLIKGISEAKADKLLGEASKLVPMGFTTATEYHIRRSELITITTGSKQLDTLLQGGVETGSITELFGEFRTGKSQICHTLAVTCQLPIDMGGGEGKCL : 180
H.s. : VEAVAYAPKKELINIKGISEAKADKILAEAAKLVPMGFTTATEFHQRRSEIIQITTGSKELDKLLQGGIETGSITEMFGEFRTGKTQICHTLAVTCQLPIDRGGGEGKAM : 158
S.c. : AEAVAYAPRKDLLEIKGISEAKADKLLNEAARLVPMGFVTAADFHMRRSELICLTTGSKNLDTLLGGGVETGSITELFGEFRTGKSQLCHTLAVTCQIPLDIGGGEGKCL : 216

                    *       240         *       260         *       280         *       300         *       320         *
T.r. : YIDTEGTFRPVRLLAVANRFGLSGEEVLDNVAYARAYNSDHQLQLLNQAAAMMCETRFSLLIVDSATSLYRTDFTGRGELSNRQTHLAKFMRTLQRLADEFGIAVVITNQ : 270
N.c. : YIDTEGTFRPVRLLAVANRYGLSGEEVLDNVAYARAYNSDHQLQLLNQAAAMMCETRFSLLIVDSATSLYRTDFLGRGELSSRQTHLAKFMRTLQRLADEFGIAVVITNQ : 272
S.p. : YIDTEGTFRPVRLLAVADRYGLNGEEVLDNVAYARAYNADHQLELLQQAANMMSESRFSLLVVDSCTALYRTDFSGRGELSARQMHLARFMRTLQRLADEFGIAVVITNQ : 290
H.s. : YIDTEGTFRPERLLAVAERYGLSGSDVLDNVAYARAFNTDHQTQLLYQASAMMVESRYALLIVDSATALYRTDYSGRGELSARQMHLARFLRMLLRLADEFGVAVVITNQ : 268
S.c. : YIDTEGTFRPVRLVSIAQRFGLDPDDALNNVAYARAYNADHQLRLLDAAAQMMSESRFSLIVVDSVMALYRTDFSGRGELSARQMHLAKFMRALQRLADQFGVAVVVTNQ : 326

                  340         *       360         *       380         *       400         *
T.r. : VVAQVDGGPSAMFNPDPKKPIGGNIIAHASTTRISLKKGRGETRIAKIYDSPCLPESDTLFAIGEDGIGDPAPKDLEKEKD : 351
N.c. : VVAQVDGGPSAMFNPDPKKPIGGNIIAHASTTRISLKKGRGETRIAKIYDSPCLPESDCLFAINEDGIGDPSPKDMEKMNQ : 353
S.p. : VVAQVDG--IS-FNPDPKKPIGGNILAHSSTTRLSLRKGRGEQRICKIYDSPCLPESEAIFAINSDGVGDPKEIIAPV--- : 365
H.s. : VVAQVDG--AAMFAADPKKPIGGNIIAHASTTRLYLRKGRGETRICKIYDSPCLPEAEAMFAINADGVGDAKD-------- : 339
S.c. : VVAQVDGG-MA-FNPDPKKPIGGNIMAHSSTTRLGFKKGKGCQRLCKVVDSPCLPEAECVFAIYEDGVGDPREEDE----- : 400
```

**Fig. S4. Rad51 homolog sequence alignment**. The amino acid sequence of *Trichoderma reesei* (*T.r.*) Rad51 protein aligned with those of *Neurospora crassa* (*N.c.*), *Schizosaccharomyces pombe* (*S.p.*), *Homo sapiens* (*H.s*) and *Saccharomyces cerevisiae* (*S.c.*).

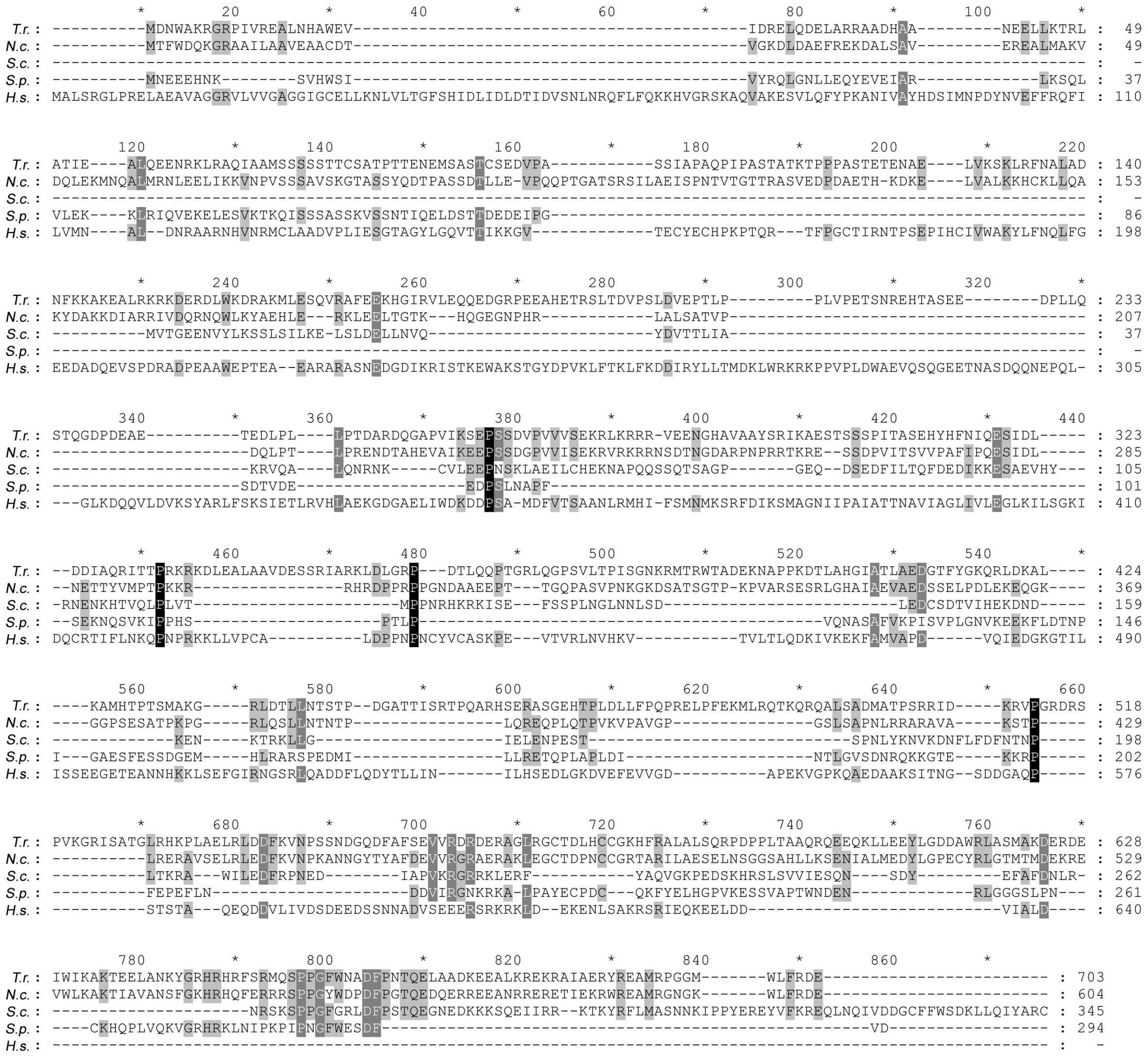

**Fig. S5. Sae2 homolog sequence alignment.** The amino acid sequence of *Trichoderma reesei* (*T.r.*) Sae2 protein aligned with those of *Neurospora crassa* (*N.c.*), *Schizosaccharomyces pombe* (*S.p.*), *Homo sapiens* (*H.s*) and *Saccharomyces cerevisiae* (*S.c.*).

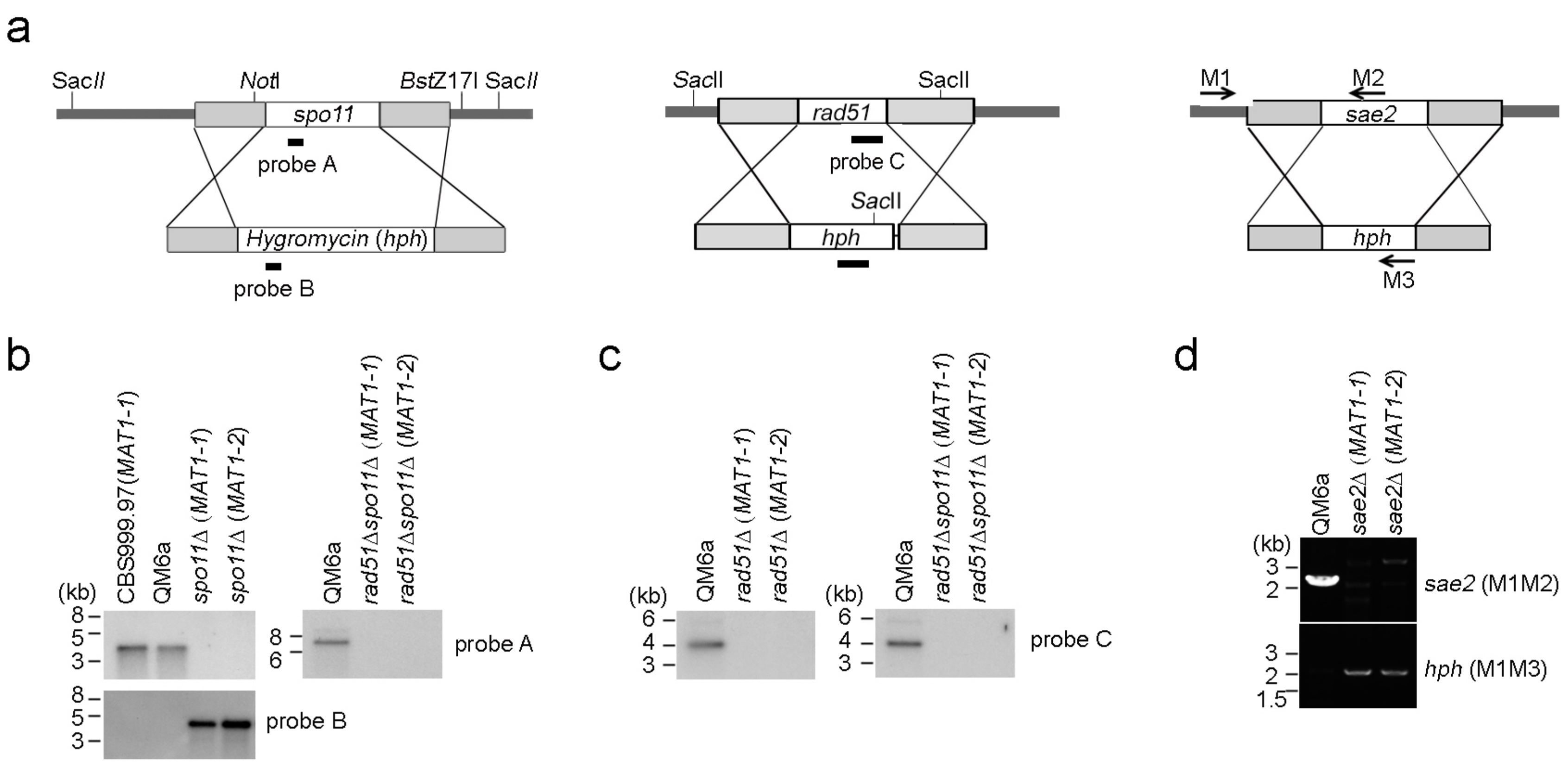

**Fig. S6. Generation of *T. reesei* mutant strains.** (**a**) The cassettes for the removal of *spo11* (upper panel), *rad51* (middle panel), and *sae2* (bottom panel), respectively. The protein-encoding regions and the hydromycin selectable marker (*hph*) are indicated. Shaded boxes represent the upstream and downstream sequences of the protein-encoding genes. The restriction enzyme sites are indicated by italics. The three DNA probes for Southern hybridization are indicated by black boxes. (**b-c**) Southern hybridization. Genomic DNA was isolated, digested by the indicated restriction enzyme(s) and then visualized by Southern blotting. (**d**) Genomic PCR. The genomic DNA was isolated and then PCR-amplified by a pair of indicated oligonucleotide primers (M1-M3). The nucleotide sequences of these PCR primers will be made available upon request.

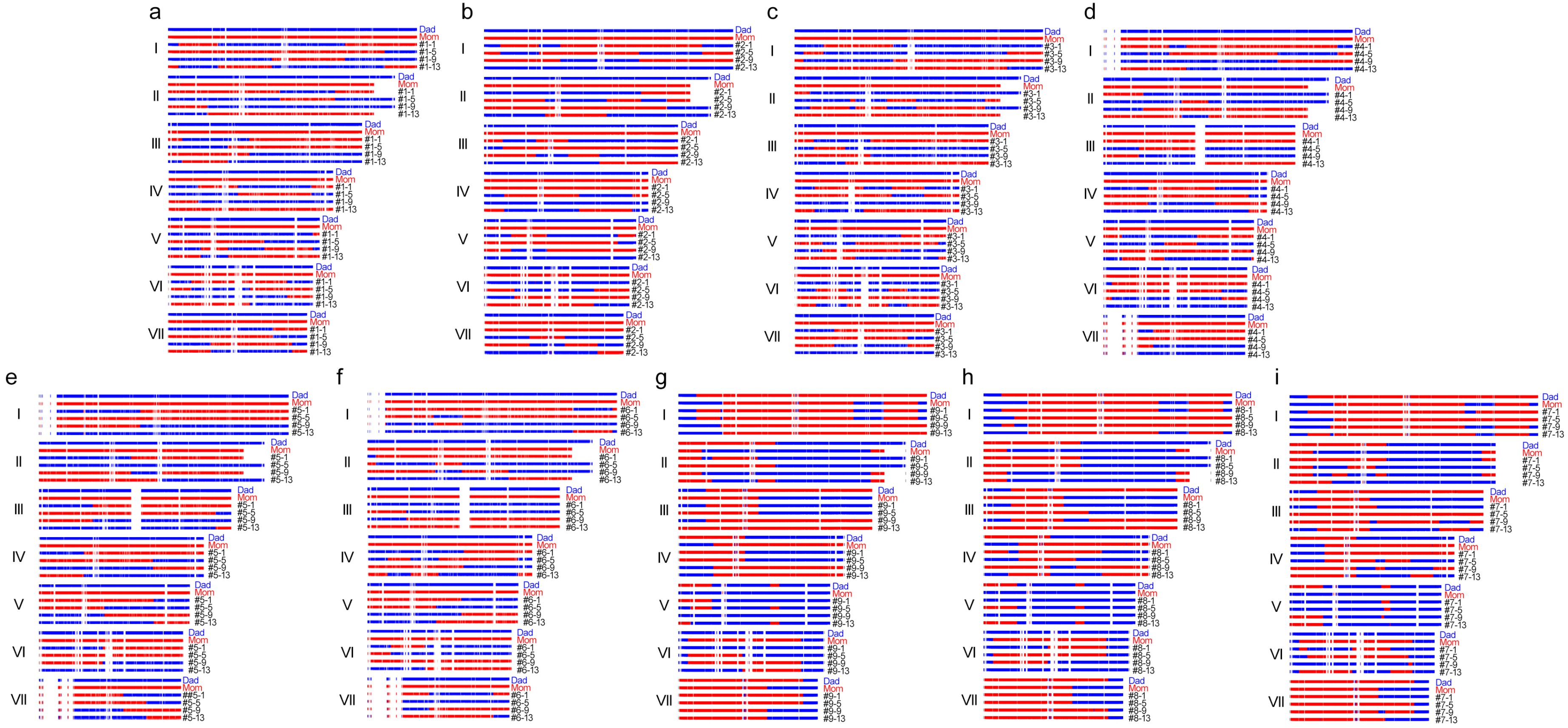

**Fig. S7. NGS analyses of *T. reesei* meiotic interhomolog recombination products.** Sequences identical to QM6a are depicted with blue bars and those identical to CBS999.97(*MAT1-1*) are represented by red bars. Empty areas are highly AT-rich sequences lacking read coverage or SNPs. The Illumina MiSeq paired ends were used to generate the genome-wide recombination profiles of (**a-c**) the three QM6a/CBS999.97(*MAT1-1*) asci (#1-#3; A-C), (**d-i**) the six QM6a *spo11*Δ/CBS999.97(*MAT1-1*) *spo11*Δ asci (#4-#9).

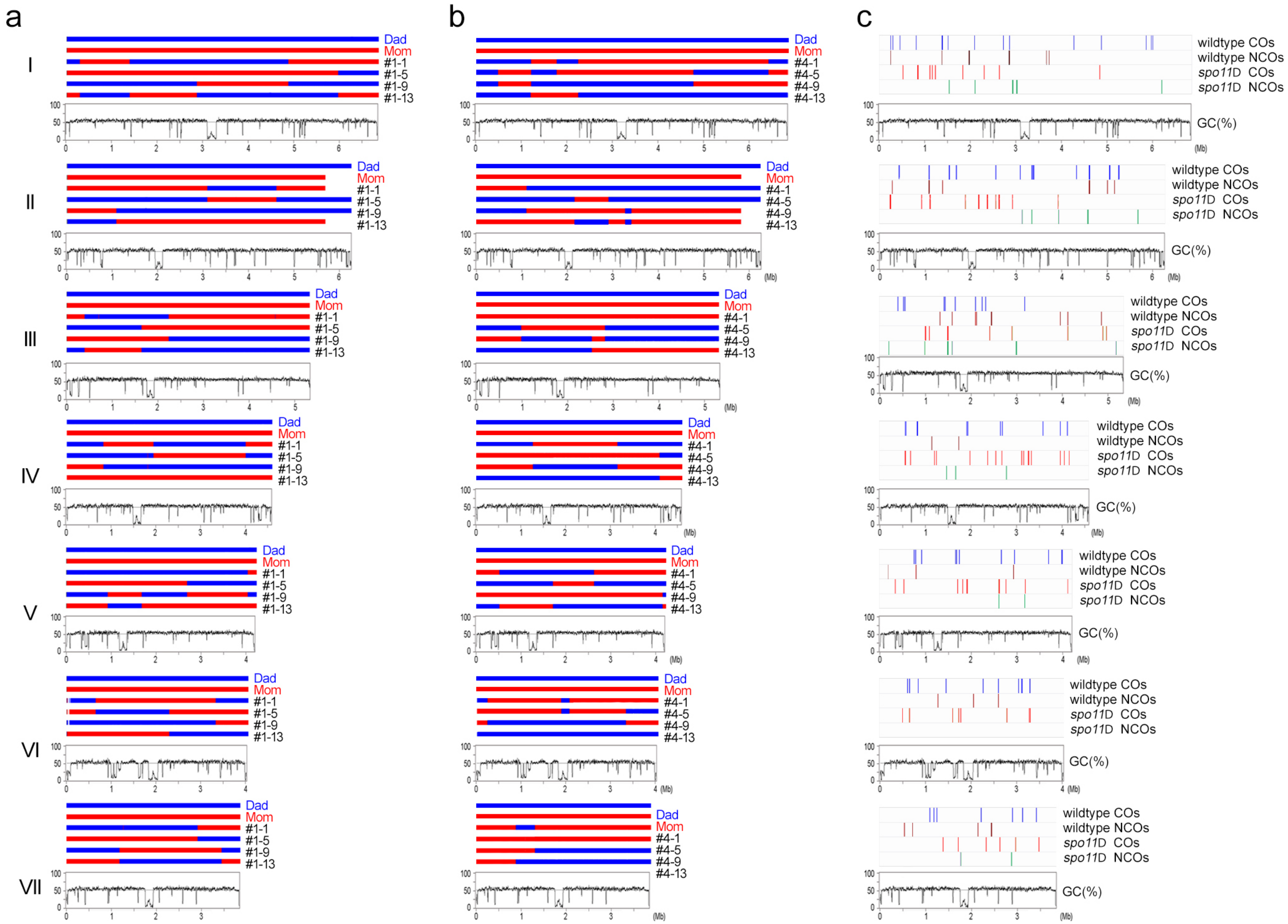

**Fig. S8.** ***Trichoderma reesei*** **meiosis generates interhomolog recombination products regardless of the presence or absence of** ***spo11*****.** Sequences identical to QM6a are depicted by blue bars, those identical to CBS999.97(*MAT1-1*) are represented by red bars. Traces represent graphs of GC content (window size 5000 bp) of the seven telomere-to-telomere chromosomes (I-VII) in QM6a. The PacBio long reads were used to generate the genome-wide recombination profiles of (**a**) QM6a/CBS999.97(*MAT1-1*) asci #1 and (**b**) QM6a *spo11*Δ/CBS999.97(*MAT1-1*) *spo11*Δ asci #4. (**c**) Overview of all interhomolog recombination products detected in the three QM6a/CBS999.97(*MAT1-1*) asci (#1-#3) and the six QM6a *spo11*Δ/CBS999.97(*MAT1-1*) *spo11*Δ asci (#4-#9). The positions of COs and NCOs on each chromosome are indicated by vertical lines.

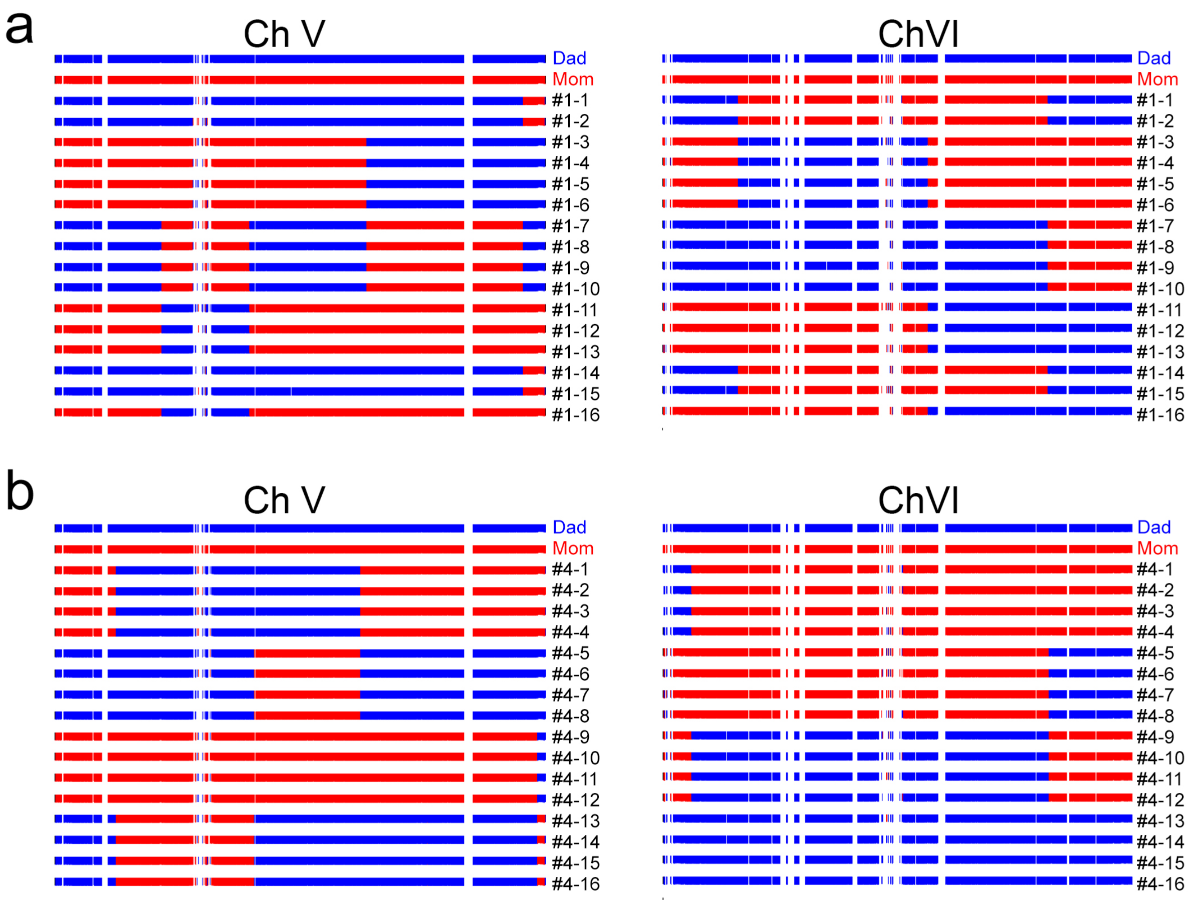

**Fig. S9. High fidelity of the two rounds of postmeiotic DNA replication regardless of the presence or absence of *spo11*.** Sequences identical to QM6a are shown as blue bars and those identical to CBS999.97(*MAT1-1*) are represented by red bars. Empty areas are AT-rich sequences lacking read coverage or SNPs. Whole genome sequencing profiles of the two chromosomes (V and VI) of all 16 F1 progeny from the QM6a/CBS999.97(*MAT1-1*) ascus #1 (**a**) and from the six QM6a *spo11*Δ/CBS999.97(*MAT1-1*) *spo11*Δ ascus #4 (**b**) were determined from Illumina MiSeq paired-end reads.

**Table S1. Summary of PacBio RSII sequencing and assembly results**

| Strain | QM6a | CBS999.97(*MAT1-1*) | CBS999.97(*MAT1-2*) |
|---|---|---|---|
| Total sequenced bases | 3,397,762,180 bp | 4,810,795,008 bp | 6,202,644,396 bp |
| Maximum of all assembled unitigs | 6,835,650 bp | 6,822,671 bp | 6,835,650 bp |
| $N_{50}$ of all assembled unitigs | 5,311,312 bp | 5,258,125 bp | 5,262,578 bp |
| Number reads | 263,312 | 370,345 | 636,376 |
| $N_{50}$ reads | 18,236 bp | 18,599 bp | 14,406 bp |
| Phred Quality Score | 48.8 | 48.1 | 48.7 |
| Unitigs | 7 linear chromosomes | 7 linear chromosomes | 7 linear chromosomes |
| Coverage | 81X | 93X | 127X |
| Genome size | 34,922,528 bp | 34,319,199 bp | 34,324,311 bp |
| Unidentified bases (N) | 0 bp | 0 bp | 0 bp |
| GC content | 51.1% | 51.6% | 51.6% |

#### Table S2. Characteristics and assembly of the seven CBS999.97 chromosomes

| Strain | Chromosome | Size (bp) | GC (%)[1] | N[2] | Centromere[3] | [4]Telomeric repeats | | L telomere + subtelomere[5] |
|---|---|---|---|---|---|---|---|---|
| | | | | | | L | R | R telomere + subtelomere[5] |
| CBS999.97(*MAT1-1*) | I | 6,822,680 | 52.09 | 0 | 3184500- 3359500<br>(~175 kb) | 12 | 10 | 1-19500<br>6812500-6822680 |
| | II | 5,559,498 | 52.40 | 0 | 1892000-2063500<br>(~172 kb) | 10 | 10 | 1-4000<br>5558500-5559498 |
| | III | 5,258,134 | 52.11 | 0 | 1677500-1830500<br>(~153 kb) | 13 | 10 | 1-4000<br>5256000-5258134 |
| | IV | 4,872,985 | 52.06 | 0 | 1453500- 1617500<br>(~164 kb) | 13 | 9 | 1-1500<br>4842500-4872985 |
| | V | 4,096,940 | 51.81 | 0 | 1102500-1259500<br>(~157 kb) | 8 | 13 | 1-17500<br>4092500-4096940 |
| | VI | 3,741,771 | 50.24 | 0 | 1636500-1810000<br>(~174 kb) | -- | 9 | 1-32500<br>3741000-3741771 |
| | VII | 3,967,191 | 49.80 | 0 | 1858000-2018500<br>(~161 kb) | 12 | 9 | 1-30000<br>3956000-3967191 |
| | Overall | 34,319,199 | 51.6 | 0 | 1071500 bp | 68 | 70 | ~157 kb |
| CBS[illegible] | I | 6,822,679 | 52.09 | 0 | 3184500- 3359500 | 14 | 7 | 1-19500 |

| | | | | | | | | |
|---|---|---|---|---|---|---|---|---|
| | | | | | (~175 kb) | | | 6814000-6822679 |
| | II | 6,061,978 | 52.37 | 0 | 1892000-2063500 (~172 kb) | 12 | 12 | 1-4000 |
| | | | | | | | | 6042500-6061978 |
| | III | 5,262,578 | 52.11 | 0 | 1677500-1830500 (~153 kb) | 14 | 7 | 1-4000 |
| | | | | | | | | 5256000-5262578 |
| | IV | 4,370,647 | 52.07 | 0 | 1453500- 1617500 (~164 kb) | 8 | 13 | 1-1500 |
| | | | | | | | | 4369500-4370647 |
| | V | 4,096,893 | 51.81 | 0 | 1102500-1259500 (~157 kb) | 10 | 7 | 1-17500 |
| | | | | | | | | 4092500-4096893 |
| | VI | 3,742,327 | 50.23 | 0 | 1637000-1810500 (~174 kb) | - | 13 | 1-33000 |
| | | | | | | | | 3741500-3742327 |
| | VII | 3,927,209 | 49.80 | 0 | 1858000-2018500 (~161 kb) | 8 | 14 | 1-30000 |
| | | | | | | | | 3956000-3967209 |
| | Overall | 34,324,311 | 51.6 | 0 | 1153000 bp | 66 | 73 | ~157 kb |

1. The EMBOSS geecee tool (http://www.bioinformatics.nl/cgi-bin/emboss/geecee) was used to calculate the fractional GC content using a 500 bp sliding window.
2. Number of unresolved bases (N).
3. Centromeres were manually identified as the longest AT-rich islands in each chromosome.
4. The telomeric repeats were identified as $(TTAGGG)_n$ at 3′ termini and the complementary sequence $(CCCTAA)_n$ at 5′ termini.
5. Subtelomeres were manually identified as the AT-rich island right next to the telomeric repeats.

**Table S3. Transposable elements in QM6a, CBS999.97(*MAT1-1*) and CBS999.97(*MAT1-2*)**

| Strain | Genome size (bp) | Repetitive sequences (bp) | Overall | Class I (retrotransposons) | | | | | | | Class II (transposons) | | | | | |
|---|---|---|---|---|---|---|---|---|---|---|---|---|---|---|---|---|
| | | | | *Tad1*-LINE | *I*-LINE | *Jockey*-LINE | other LINEs | *Copia*-LTR | *Gypsy*-LTR | other LTRs | *CMC-EnSpm* | *hAT* | *MULE-MuDR* | *TcMar -Fot1* | *PIF-Harbinger-like* | Others |
| | | | copy number | | | | | | | | | | | | | |
| QM6a | 34922528 | 42035 | 70 | 0 | 0 | 4 | 14 | 8 | 10 | 4 | 6 | 0 | 21 | 0 | 0 | 3 |
| CBS999.97 (*MAT1-1*) | 34319199 | 20756 | 62 | 0 | 0 | 11 | 11 | 5 | 2 | 6 | 4 | 1 | 17 | 0 | 0 | 5 |
| CBS999.97 (*MAT1-2*) | 34324311 | 20756 | 62 | 0 | 0 | 11 | 11 | 5 | 2 | 6 | 4 | 1 | 17 | 0 | 0 | 5 |

**Table S4. Size distribution of AT-rich blocks in QM6a, CBS999.97(*MAT1-1*) and CBS999.97(*MAT1-2*)**

| Strain | Genome (Mb) | Chromosome number | GC (%) | Number of AT-rich blocks of different lengths (L in kb)[1] | | | | | | | | | |
|---|---|---|---|---|---|---|---|---|---|---|---|---|---|
| | | | | 0.5 ≤ L < 1 | 1 ≤ L < 3 | 3 ≤ L < 5 | 5 ≤ L < 10 | 10 ≤ L < 15 | 15 ≤ L < 20 | 20 ≤ L < 50 | 50 ≤ L < 100 | L<100 | Total |
| QM6a | 34.9 | 7 | 51.1 ± 11.6 | 1845 | 337 | 69 | 38 | 26 | 13 | 9 | 6 | 6 | 2249 |
| CBS999.97(*MAT1-1*) | 34.3 | 7 | 51.6 ± 10.7 | 1861 | 272 | 51 | 23 | 20 | 10 | 10 | 7 | 5 | 2259 |
| CBS999.97(*MAT1-2*) | 34.3 | 7 | 51.6 ± 10.8 | 1826 | 301 | 50 | 21 | 20 | 9 | 10 | 8 | 5 | 2250 |

1. AT-rich interspersed islands were identified by their AT contents being ≥6% higher than that of average DNA in each fungal genome.

**Table S5. SNPs between CBS999.97(*MAT1-1*) and CBS999.97(*MAT1-2*)**

| Chromosome | Total SNPs | Nucleotide variation (point mutation) | Insertion / Deletion (Indel) |
|---|---|---|---|
| I | 404 | 73 | 331 |
| II | 325 | 116 | 209 |
| III | 616 | 301 | 315 |
| IV | 265 | 43 | 222 |
| V | 271 | 79 | 192 |
| VI | 474 | 147 | 327 |
| VII | 350 | 180 | 170 |
| Genome | 2705 | 939 | 1766 |

**Table S6. SNPs between QM6a and CBS999.97(*MAT1-1*)**

| Chromosome | Total SNPs | Nucleotide variation (point mutation) | Insertion / Deletion (Indel) |
|---|---|---|---|
| I | 188,729 | 120,810 | 67,919 |
| II | 159,995 | 102,547 | 57,448 |
| III | 136,910 | 88,503 | 48,407 |
| IV | 122867 | 78,085 | 44,782 |
| V | 115,253 | 72,582 | 42,671 |
| VI | 104,773 | 70,501 | 34,272 |
| VII | 107,472 | 71,550 | 35,992 |
| Genome | 935,999 | 604,578 | 331,491 |

**Table S7. SNPs between QM6a and CBS999.97(*MAT1-2*)**

| Chromosome | Total SNPs | Nucleotide variation (point mutation) | Insertion / Deletion (Indel) |
|---|---|---|---|
| I | 188,658 | 120,800 | 67,858 |
| II | 173,152 | 110,865 | 62,287 |
| III | 136,994 | 88,518 | 48,476 |
| IV | 123,284 | 78,370 | 44,914 |
| V | 115,260 | 72,579 | 42,681 |
| VI | 104,620 | 70,258 | 34,362 |
| VII | 107,480 | 71,557 | 35,923 |
| Genome | 949,448 | 612,947 | 336,501 |

**Table S8.** ***Trichoderma reesei spo11*** **is dispensable for meiosis and meiosis-driven segmental aneuploidy (SAN)**

| Sexual crossing | Number of asci dissected | Number of asci with 4 or 8 viable SAN ascospores |
|---|---|---|
| CBS999.97(*MAT1-1*, II_D, IV_L) X CBS999.97(*MAT1-2*, II_D, IV_L) | 20 | 0 |
| CBS999.97(*MAT1-1*, II_L, IV_D) X CBS999.97(*MAT1-2*, II_L, IV_D) | 20 | 0 |
| CBS999.97(*MAT1-1*, II_L, IV_D) X CBS999.97(*MAT1-2*, II_D, IV_L) | 20 | 19 |
| CBS999.97(*MAT1-1*, II_D, IV_L) X CBS999.97(*MAT1-2*, II_L, IV_D) | 21 | 19 |
| *spo11*Δ (*MAT1-1*, II_D, IV_L) X *spo11*Δ (*MAT1-2*, II_D, IV_L) | 26 | 0 |
| *spo11*Δ (*MAT1-1*, II_L, IV_D) X *spo11*Δ (*MAT1-2*, II_D, IV_L) | 22 | 19 |

**Table S9. Summary of COs and NCOs (with ≥ 2 SNP) in a representative SK1/S288c tetrad.**

| Tetrad number | Interhomolog recombination | Chromosome number | Chromatids | SNP_start_out | SNP_start_in | SNP_end_in | SNP_end_out | Position of CO | GC tract associated with CO | GC tract length (bp) | CO length (bp) |
|---|---|---|---|---|---|---|---|---|---|---|---|
| 11 | CO | 1 | 2,4 | 175318 | 175409 | 173966 | 174674 | 174842 | yes | 1044 | 1044 |
| 11 | CO | 1 | 2,4 | 137336 | 138499 | 138650 | 139272 | 138439 | yes | 1044 | 1044 |
| 11 | CO | 2 | 2,4 | 319306 | 320491 | 319306 | 320491 | 319899 | no | 0 | 1185 |
| 11 | CO | 2 | 2,4 | 406115 | 406787 | 406787 | 407477 | 406792 | yes | 681 | 681 |
| 11 | CO | 2 | 1,3 | 194226 | 196145 | 196745 | 197035 | 196038 | yes | 1705 | 1705 |
| 11 | CO | 2 | 1,4 | 651616 | 652362 | 651616 | 652362 | 651989 | no | 0 | 746 |
| 11 | CO | 3 | 1,3 | 63611 | 63845 | 63611 | 63845 | 63728 | no | 0 | 234 |
| 11 | CO | 3 | 3,4 | 242917 | 246581 | 242917 | 246581 | 244749 | no | 0 | 3664 |
| 11 | CO | 3 | 1,4 | 164277 | 165283 | 162442 | 162686 | 163672 | yes | 2216 | 2216 |
| 11 | CO | 4 | 1,4 | 83841 | 84443 | 83841 | 84443 | 84142 | no | 0 | 602 |
| 11 | CO | 4 | 1,4 | 1056448 | 1056586 | 1056739 | 1057249 | 1056756 | yes | 477 | 477 |
| 11 | CO | 4 | 1,2 | 615816 | 615959 | 617533 | 617915 | 616806 | yes | 1837 | 1837 |

| | | | | | | | | | | | |
|---|---|---|---|---|---|---|---|---|---|---|---|
| 11 | CO | 4 | 2,4 | 682954 | 683240 | 682303 | 682954 | 682863 | yes | 469 | 469 |
| 11 | CO | 4 | 3,4 | 176132 | 176206 | 174998 | 175763 | 175775 | yes | 789 | 789 |
| 11 | CO | 4 | 2,4 | 1293368 | 1293471 | 1296525 | 1296732 | 1295024 | yes | 3209 | 3209 |
| 11 | CO | 4 | 1,4 | 555425 | 555588 | 555425 | 555588 | 555507 | no | 0 | 163 |
| 11 | CO | 4 | 1,3 | 418485 | 418803 | 418803 | 419073 | 418791 | yes | 294 | 294 |
| 11 | CO | 4 | 2,3 | 1179516 | 1179751 | 1176974 | 1177557 | 1178450 | yes | 2368 | 2368 |
| 11 | CO | 5 | 1,4 | 309648 | 309961 | 309648 | 309961 | 309805 | no | 0 | 313 |
| 11 | CO | 5 | 3,4 | 483903 | 485006 | 482316 | 483903 | 483782 | yes | 1345 | 1345 |
| 11 | CO | 5 | 3,4 | 463625 | 464078 | 460411 | 461582 | 462424 | yes | 2855 | 2855 |
| 11 | CO | 5 | 2,3 | 63585 | 64340 | 66291 | 67131 | 65337 | yes | 2749 | 2749 |
| 11 | CO | 6 | 2,4 | 226036 | 226370 | 223531 | 223558 | 224874 | yes | 2659 | 2659 |
| 11 | CO | 6 | 1,4 | 77013 | 77181 | 75475 | 76053 | 76431 | yes | 1333 | 1333 |
| 11 | CO | 6 | 1,2 | 227587 | 228299 | 226615 | 226776 | 227319 | yes | 1248 | 1248 |
| 11 | CO | 7 | 3,4 | 1010428 | 1010860 | 1011203 | 1011474 | 1010991 | yes | 695 | 695 |
| 11 | CO | 7 | 1,3 | 763517 | 763747 | 763036 | 763187 | 763372 | yes | 521 | 521 |
| 11 | CO | 7 | 1,3 | 610674 | 611145 | 610674 | 611145 | 610910 | no | 0 | 471 |
| 11 | CO | 7 | 3,4 | 226711 | 228148 | 223151 | 224665 | 225669 | yes | 3522 | 3522 |
| 11 | CO | 7 | 2,3 | 431198 | 431867 | 433234 | 433416 | 432429 | yes | 1793 | 1793 |
| 11 | CO | 7 | 1,4 | 913000 | 913222 | 912779 | 913000 | 913000 | yes | 222 | 222 |
| 11 | CO | 7 | 3,4 | 245686 | 247666 | 241708 | 244757 | 244954 | yes | 3444 | 3444 |
| 11 | CO | 8 | 3,4 | 248808 | 250465 | 248808 | 250465 | 249637 | no | 0 | 1657 |
| 11 | CO | 8 | 1,3 | 318584 | 319046 | 319904 | 320927 | 319615 | yes | 1601 | 1601 |

| | | | | | | | | | | | |
|---|---|---|---|---|---|---|---|---|---|---|---|
| 11 | CO | 8 | 2,4 | 278461 | 278994 | 277301 | 278461 | 278304 | yes | 847 | 847 |
| 11 | CO | 8 | 2,3 | 62846 | 62950 | 59801 | 59916 | 61378 | yes | 3040 | 3040 |
| 11 | CO | 8 | 2,3 | 120445 | 121477 | 121987 | 122655 | 121641 | yes | 1360 | 1360 |
| 11 | CO | 9 | 3,4 | 133668 | 135075 | 132756 | 133353 | 133713 | yes | 1317 | 1317 |
| 11 | CO | 9 | 1,3 | 329125 | 332459 | 333073 | 333345 | 332001 | yes | 2417 | 2417 |
| 11 | CO | 9 | 3,4 | 179673 | 180009 | 181236 | 181677 | 180429 | yes | 1176 | 1176 |
| 11 | CO | 10 | 1,3 | 455458 | 456781 | 453740 | 454963 | 455236 | yes | 1768 | 1768 |
| 11 | CO | 10 | 1,3 | 238135 | 238262 | 235976 | 236393 | 237192 | yes | 2014 | 2014 |
| 11 | CO | 10 | 2,4 | 574799 | 576464 | 577732 | 578943 | 576985 | yes | 2706 | 2706 |
| 11 | CO | 10 | 1,4 | 334204 | 334468 | 334468 | 334858 | 334500 | yes | 327 | 327 |
| 11 | CO | 10 | 3,4 | 315022 | 315237 | 315888 | 316551 | 315675 | yes | 1090 | 1090 |
| 11 | CO | 11 | 1,3 | 519020 | 519321 | 506614 | 506935 | 512973 | yes | 12396 | 12396 |
| 11 | CO | 11 | 2,4 | 362705 | 363248 | 364384 | 364596 | 363733 | yes | 1514 | 1514 |
| 11 | CO | 11 | 1,3 | 578230 | 578669 | 576209 | 576947 | 577514 | yes | 1872 | 1872 |
| 11 | CO | 12 | 2,4 | 254321 | 254610 | 254773 | 255134 | 254710 | yes | 488 | 488 |
| 11 | CO | 12 | 2,3 | 126014 | 126853 | 126014 | 126853 | 126434 | no | 0 | 839 |
| 11 | CO | 12 | 1,2 | 397870 | 398071 | 397049 | 397180 | 397543 | yes | 856 | 856 |
| 11 | CO | 12 | 3,4 | 446902 | 447260 | 446652 | 446902 | 446929 | yes | 304 | 304 |
| 11 | CO | 12 | 2,4 | 827928 | 828244 | 827928 | 828244 | 828086 | no | 0 | 316 |
| 11 | CO | 13 | 1,4 | 160146 | 160280 | 162123 | 162309 | 161215 | yes | 2003 | 2003 |
| 11 | CO | 13 | 1,3 | 99367 | 99579 | 98310 | 98939 | 99049 | yes | 849 | 849 |
| 11 | CO | 13 | 1,4 | 428833 | 429465 | 434958 | 435813 | 432046 | yes | 5794 | 5794 |

| | | | | | | | | | | | |
|---|---|---|---|---|---|---|---|---|---|---|---|
| 11 | CO | 13 | 2,3 | 772812 | 772883 | 775426 | 775628 | 774187 | yes | 2680 | 2680 |
| 11 | CO | 13 | 3,4 | 825884 | 826148 | 828384 | 828518 | 827234 | yes | 2435 | 2435 |
| 11 | CO | 13 | 2,3 | 193159 | 194053 | 194821 | 195330 | 194341 | yes | 1470 | 1470 |
| 11 | CO | 13 | 1,4 | 760322 | 760768 | 761015 | 761300 | 760851 | yes | 613 | 613 |
| 11 | CO | 13 | 2,4 | 20622 | 21082 | 18729 | 20392 | 20206 | yes | 1292 | 1292 |
| 11 | CO | 14 | 1,2 | 413686 | 414010 | 412369 | 412378 | 413111 | yes | 1475 | 1475 |
| 11 | CO | 14 | 1,4 | 439088 | 440622 | 442806 | 443263 | 441445 | yes | 3180 | 3180 |
| 11 | CO | 14 | 3,4 | 762860 | 763381 | 766008 | 766895 | 764786 | yes | 3331 | 3331 |
| 11 | CO | 14 | 1,3 | 689582 | 689944 | 691115 | 692612 | 690813 | yes | 2101 | 2101 |
| 11 | CO | 14 | 3,4 | 255192 | 255432 | 252203 | 254039 | 254217 | yes | 2191 | 2191 |
| 11 | CO | 15 | 2,3 | 164383 | 164556 | 159369 | 159827 | 162034 | yes | 4872 | 4872 |
| 11 | CO | 15 | 2,3 | 698870 | 700016 | 698870 | 700016 | 699443 | no | 0 | 1146 |
| 11 | CO | 15 | 2,4 | 701852 | 702223 | 700551 | 700853 | 701370 | yes | 1336 | 1336 |
| 11 | CO | 15 | 1,4 | 212987 | 213206 | 213775 | 214699 | 213667 | yes | 1141 | 1141 |
| 11 | CO | 15 | 2,3 | 128123 | 128405 | 127669 | 128123 | 128080 | yes | 368 | 368 |
| 11 | CO | 15 | 3,4 | 780631 | 780750 | 781222 | 781811 | 781104 | yes | 826 | 826 |
| 11 | CO | 15 | 2,3 | 128581 | 128721 | 128405 | 128530 | 128559 | yes | 184 | 184 |
| 11 | CO | 15 | 1,4 | 486511 | 487028 | 486511 | 487028 | 486770 | no | 0 | 517 |
| 11 | CO | 16 | 1,4 | 432288 | 432416 | 429951 | 430119 | 431194 | yes | 2317 | 2317 |
| 11 | CO | 16 | 1,4 | 890080 | 890467 | 887709 | 888159 | 889104 | yes | 2340 | 2340 |
| 11 | CO | 16 | 3,4 | 680837 | 681090 | 680837 | 681090 | 680964 | no | 0 | 253 |
| 11 | CO | 16 | 3,4 | 339114 | 339687 | 339114 | 339687 | 339401 | no | 0 | 573 |

| | | | | | | | | | | | |
|---|---|---|---|---|---|---|---|---|---|---|---|
| 11 | CO | 16 | 2,4 | 677025 | 677287 | 680750 | 680822 | 678971 | yes | 3630 | 3630 |

| **Asci number** | **Interhomolog recombination** | **Chromosome number** | **Chromatids** | **SNP_start** | **SNP_end** | **Position of NCO (bp)** | **SNP number** | **NCO length (bp)** |
|---|---|---|---|---|---|---|---|---|
| 11 | NCO | 1 | D | 130292 | 130362 | 130362 | 2 | 70 |
| 11 | NCO | 2 | C | 171433 | 172681 | 172057 | 15 | 1248 |
| 11 | NCO | 2 | B | 255127 | 255743 | 255435 | 4 | 616 |
| 11 | NCO | 2 | A | 321309 | 321309 | 321309 | 1 | 0 |
| 11 | NCO | 2 | A | 322823 | 322823 | 322823 | 1 | 0 |
| 11 | NCO | 2 | B | 502305 | 502627 | 502466 | 3 | 322 |
| 11 | NCO | 3 | C | 51152 | 51650 | 51401 | 5 | 498 |
| 11 | NCO | 4 | C | 1057267 | 1057601 | 1057434 | 4 | 334 |
| 11 | NCO | 4 | D | 1257108 | 1257111 | 1257110 | 2 | 3 |
| 11 | NCO | 5 | D | 128140 | 129628 | 128884 | 11 | 1488 |
| 11 | NCO | 5 | D | 262462 | 262462 | 262462 | 1 | 0 |
| 11 | NCO | 5 | D | 310175 | 310540 | 310357.5 | 2 | 365 |
| 11 | NCO | 5 | A | 399399 | 400915 | 400157 | 7 | 1516 |
| 11 | NCO | 6 | A | 77256 | 77256 | 77256 | 1 | 0 |
| 11 | NCO | 7 | C | 54723 | 56199 | 55461 | 12 | 1476 |
| 11 | NCO | 7 | D | 393263 | 394661 | 393962 | 10 | 1398 |

| | | | | | | | | |
|---|---|---|---|---|---|---|---|---|
| 11 | NCO | 7 | D | 849134 | 849797 | 849465.5 | 9 | 663 |
| 11 | NCO | 7 | A | 914517 | 916059 | 915288 | 6 | 1542 |
| 11 | NCO | 7 | B | 1009812 | 1009812 | 1009812 | 1 | 0 |
| 11 | NCO | 8 | C | 114144 | 114162 | 114153 | 3 | 18 |
| 11 | NCO | 8 | C | 114255 | 422992 | 268623.5 | 6 | 308737 |
| 11 | NCO | 10 | B | 464252 | 466454 | 465353 | 7 | 2202 |
| 11 | NCO | 10 | A | 690731 | 691098 | 690914.5 | 4 | 367 |
| 11 | NCO | 11 | C | 78581 | 80353 | 79467 | 11 | 1772 |
| 11 | NCO | 11 | D | 225480 | 228339 | 226909.5 | 12 | 2859 |
| 11 | NCO | 11 | C | 361983 | 361983 | 361983 | 1 | 0 |
| 11 | NCO | 11 | D | 367554 | 367554 | 367554 | 1 | 0 |
| 11 | NCO | 12 | A | 604090 | 604115 | 604102.5 | 2 | 25 |
| 11 | NCO | 12 | C | 967075 | 967302 | 967188.5 | 2 | 227 |
| 11 | NCO | 13 | D | 275405 | 275405 | 275405 | 1 | 0 |
| 11 | NCO | 13 | A | 674192 | 677544 | 675868 | 21 | 3352 |
| 11 | NCO | 13 | B | 829431 | 831576 | 830503.5 | 11 | 2145 |
| 11 | NCO | 15 | B | 988081 | 989690 | 988885.5 | 9 | 1609 |
| 11 | NCO | 16 | C | 899206 | 900649 | 899927.5 | 17 | 1443 |

**Table S10. List of all *Trichoderma reesei* strains analyzed by whole genome sequencing technology**

| Strain | Description | Sequencing platform | BioProject accession | WGS/SRA accession |
|---|---|---|---|---|
| CBS999.97(*MAT1-1*) | Wild type (*MAT1-1*) | PacBio RSII | PRJNA352653 | CP017983-CP017984, CP020875-CP020879 |
| CBS999.97(*MAT1-2*) | Wild type (*MAT1-2*) | PacBio RSII | PRJNA382020 | CP020724-CP020730 |
| #1 QM6a *spo11*Δ | The parental strain of #4-#6 asci | PacBio Sequel | PRJNA433292 | CP040222-CP040228 |
| #1 CBS999.97(MAT1-1) *spo11*Δ | The parental strain of #4-#6 asci | PacBio Sequel | PRJNA433292 | CP040215-CP040221 |
| #1 QM6a *spo11*Δ | The parental strain of #4-#6 asci | ILLUMINA NextSeq 500 | PRJNA433292 | SRR6884639 |
| #1 CBS999.97(MAT1-1) *spo11*Δ | The parental strain of #4-#6 asci | ILLUMINA NextSeq 500 | PRJNA433292 | SRR6884638 |
| #2 QM6a *spo11*Δ | The parental strain of #7-#9 asci | ILLUMINA NextSeq 500 | PRJNA433292 | SRR6884679 |
| #2 CBS999.97(MAT1-1) *spo11*Δ | The parental strain of #7-#9 asci | ILLUMINA NextSeq 500 | PRJNA433292 | SRR6884678 |

| | | | | |
|---|---|---|---|---|
| #1-1 | The #1 ascospore in the #1 QM6a/CBS999.97(MAT1-1) ascus | ILLUMINA NextSeq 500 | PRJNA433292 | SRR6884669 |
| #1-1 | The #1 ascospore in the #1 QM6a/CBS999.97(MAT1-1) ascus | PacBio RSII | PRJNA386077 | CP021290-CP021296 |
| #1-2 | The #2 ascospore in the #1 QM6a/CBS999.97(MAT1-1) ascus | ILLUMINA NextSeq 500 | PRJNA433292 | SRR6884668 |
| #1-3 | The #3 ascospore in the #1 QM6a/CBS999.97(MAT1-1) ascus | ILLUMINA NextSeq 500 | PRJNA433292 | SRR6884667 |
| #1-3 | The #3 ascospore in the #1 QM6a/CBS999.97(MAT1-1) ascus | PacBio RSII | PRJNA386077 | CP021297-CP021303 |
| #1-4 | The #4 ascospore in the #1 QM6a/CBS999.97(MAT1-1) ascus | ILLUMINA NextSeq 500 | PRJNA433292 | SRR6884666 |
| #1-5 | The #5 ascospore in the #1 QM6a/CBS999.97(MAT1-1) ascus | ILLUMINA NextSeq 500 | PRJNA433292 | SRR6884673 |
| #1-6 | The #6 ascospore in the #1 QM6a/CBS999.97(MAT1-1) ascus | ILLUMINA NextSeq 500 | PRJNA433292 | SRR6884672 |
| #1-7 | The #7 ascospore in the #1 QM6a/CBS999.97(MAT1-1) ascus | ILLUMINA NextSeq 500 | PRJNA433292 | SRR6884671 |

| | | | | |
|---|---|---|---|---|
| #1-7 | The #7 ascospore in the #1 QM6a/CBS999.97(MAT1-1) ascus | PacBio RSII | PRJNA386077 | CP021304-CP021310 |
| #1-8 | The #8 ascospore in the #1 QM6a/CBS999.97(MAT1-1) ascus | ILLUMINA NextSeq 500 | PRJNA433292 | SRR6884670 |
| #1-9 | The #9 ascospore in the #1 QM6a/CBS999.97(MAT1-1) ascus | ILLUMINA NextSeq 500 | PRJNA433292 | SRR6884665 |
| #1-10 | The #10 ascospore in the #1 QM6a/CBS999.97(MAT1-1) ascus | ILLUMINA NextSeq 500 | PRJNA433292 | SRR6884664 |
| #1-11 | The #11 ascospore in the #1 QM6a/CBS999.97(MAT1-1) ascus | ILLUMINA NextSeq 500 | PRJNA433292 | SRR6884649 |
| #1-11 | The #11 ascospore in the #1 QM6a/CBS999.97(MAT1-1) ascus | PacBio RSII | PRJNA386077 | CP021311-CP021317 |
| #1-12 | The #12 ascospore in the #1 QM6a/CBS999.97(MAT1-1) ascus | ILLUMINA NextSeq 500 | PRJNA433292 | SRR6884648 |
| #1-13 | The #13 ascospore in the #1 QM6a/CBS999.97(MAT1-1) ascus | ILLUMINA NextSeq 500 | PRJNA433292 | SRR6884651 |
| #1-14 | The #14 ascospore in the #1 QM6a/CBS999.97(MAT1-1) ascus | ILLUMINA NextSeq 500 | PRJNA433292 | SRR6884650 |

| | | | | |
|---|---|---|---|---|
| #1-15 | The #15 ascospore in the #1 QM6a/CBS999.97(MAT1-1) ascus | ILLUMINA NextSeq 500 | PRJNA433292 | SRR6884653 |
| #1-16 | The #16 ascospore in the #1 QM6a/CBS999.97(MAT1-1) ascus | ILLUMINA NextSeq 500 | PRJNA433292 | SRR6884652 |
| #2-1 | The #1 ascospore in the #2 QM6a/CBS999.97(MAT1-1) ascus | ILLUMINA NextSeq 500 | PRJNA433292 | SRR6884655 |
| #2-5 | The #5 ascospore in the #2 QM6a/CBS999.97(MAT1-1) ascus | ILLUMINA NextSeq 500 | PRJNA433292 | SRR6884654 |
| #2-9 | The #9 ascospore in the #2 QM6a/CBS999.97(MAT1-1) ascus | ILLUMINA NextSeq 500 | PRJNA433292 | SRR6884647 |
| #2-13 | The #13 ascospore in the #2 QM6a/CBS999.97(MAT1-1) ascus | ILLUMINA NextSeq 500 | PRJNA433292 | SRR6884646 |
| #3-1 | The #1 ascospore in the #3 QM6a/CBS999.97(MAT1-1) ascus | ILLUMINA NextSeq 500 | PRJNA433292 | SRR6884642 |
| #3-5 | The #5 ascospore in the #3 QM6a/CBS999.97(MAT1-1) ascus | ILLUMINA NextSeq 500 | PRJNA433292 | SRR6884643 |
| #3-9 | The #9 ascospore in the #3 QM6a/CBS999.97(MAT1-1) ascus | ILLUMINA NextSeq 500 | PRJNA433292 | SRR6884644 |

| | | | | |
|---|---|---|---|---|
| #3-13 | The #13 ascospore in the #3 QM6a/CBS999.97(MAT1-1) ascus | ILLUMINA NextSeq 500 | PRJNA433292 | SRR6884645 |
| #4-1 | The #1 ascospore in the #4 QM6a *spo11*Δ/CBS999.97(MAT1-1) *spo11*Δ ascus | ILLUMINA NextSeq 500 | PRJNA433292 | SRR6884640 |
| #4-1 | The #1 ascospore in the #4 QM6a *spo11*Δ/CBS999.97(MAT1-1) *spo11*Δ ascus | PacBio Sequel | PRJNA433292 | CP040187-CP040193 |
| #4-2 | The #2 ascospore in the #4 QM6a *spo11*Δ/CBS999.97(MAT1-1) *spo11*Δ ascus | ILLUMINA NextSeq 500 | PRJNA433292 | SRR6884641 |
| #4-3 | The #3 ascospore in the #4 QM6a *spo11*Δ/CBS999.97(MAT1-1) *spo11*Δ ascus | ILLUMINA NextSeq 500 | PRJNA433292 | SRR6884636 |
| #4-4 | The #4 ascospore in the #4 QM6a *spo11*Δ/CBS999.97(MAT1-1) *spo11*Δ ascus | ILLUMINA NextSeq 500 | PRJNA433292 | SRR6884637 |
| #4-5 | The #5 ascospore in the #4 QM6a *spo11*Δ/CBS999.97(MAT1-1) *spo11*Δ ascus | ILLUMINA NextSeq 500 | PRJNA433292 | SRR6884629 |
| #4-5 | The #5 ascospore in the #4 QM6a *spo11*Δ/CBS999.97(MAT1-1) *spo11*Δ ascus | PacBio Sequel | PRJNA433292 | CP040194-CP040200 |
| #4-6 | The #6 ascospore in the #4 QM6a *spo11*Δ/CBS999.97(MAT1-1) *spo11*Δ ascus | ILLUMINA NextSeq 500 | PRJNA433292 | SRR6884628 |

| | | | | |
|---|---|---|---|---|
| #4-7 | The #7 ascospore in the #4 QM6a *spo11*Δ/CBS999.97(MAT1-1) *spo11*Δ ascus | ILLUMINA NextSeq 500 | PRJNA433292 | SRR6884627 |
| #4-8 | The #8 ascospore in the #4 QM6a *spo11*Δ/CBS999.97(MAT1-1) *spo11*Δ ascus | ILLUMINA NextSeq 500 | PRJNA433292 | SRR6884626 |
| #4-9 | The #9 ascospore in the #4 QM6a *spo11*Δ/CBS999.97(MAT1-1) *spo11*Δ ascus | ILLUMINA NextSeq 500 | PRJNA433292 | SRR6884633 |
| #4-9 | The #9 ascospore in the #4 QM6a *spo11*Δ/CBS999.97(MAT1-1) *spo11*Δ ascus | PacBio Sequel | PRJNA433292 | CP040201-CP040207 |
| #4-10 | The #10 ascospore in the #4 QM6a *spo11*Δ/CBS999.97(MAT1-1) *spo11*Δ ascus | ILLUMINA NextSeq 500 | PRJNA433292 | SRR6884632 |
| #4-11 | The #11 ascospore in the #4 QM6a *spo11*Δ/CBS999.97(MAT1-1) *spo11*Δ ascus | ILLUMINA NextSeq 500 | PRJNA433292 | SRR6884631 |
| #4-12 | The #12 ascospore in the #4 QM6a *spo11*Δ/CBS999.97(MAT1-1) *spo11*Δ ascus | ILLUMINA NextSeq 500 | PRJNA433292 | SRR6884630 |
| #4-13 | The #13 ascospore in the #4 QM6a *spo11*Δ/CBS999.97(MAT1-1) *spo11*Δ ascus | ILLUMINA NextSeq 500 | PRJNA433292 | SRR6884635 |
| #4-13 | The #13 ascospore in the #4 QM6a *spo11*Δ/CBS999.97(MAT1-1) *spo11*Δ ascus | PacBio Sequel | PRJNA433292 | CP040208-CP040214 |

| | | | | |
|---|---|---|---|---|
| #4-14 | The #14 ascospore in the #4 QM6a *spo11*Δ/CBS999.97(MAT1-1) *spo11*Δ ascus | ILLUMINA NextSeq 500 | PRJNA433292 | SRR6884634 |
| #4-15 | The #15 ascospore in the #4 QM6a *spo11*Δ/CBS999.97(MAT1-1) *spo11*Δ ascus | ILLUMINA NextSeq 500 | PRJNA433292 | SRR6884710 |
| #4-16 | The #16 ascospore in the #4 QM6a *spo11*Δ/CBS999.97(MAT1-1) *spo11*Δ ascus | ILLUMINA NextSeq 500 | PRJNA433292 | SRR6884711 |
| #5-1 | The #1 ascospore in the #5 QM6a *spo11*Δ/CBS999.97(MAT1-1) *spo11*Δ ascus | ILLUMINA NextSeq 500 | PRJNA433292 | SRR6884708 |
| #5-2 | The #2 ascospore in the #5 QM6a *spo11*Δ/CBS999.97(MAT1-1) *spo11*Δ ascus | ILLUMINA NextSeq 500 | PRJNA433292 | SRR6884709 |
| #5-3 | The #3 ascospore in the #5 QM6a *spo11*Δ/CBS999.97(MAT1-1) *spo11*Δ ascus | ILLUMINA NextSeq 500 | PRJNA433292 | SRR6884706 |
| #5-4 | The #4 ascospore in the #5 QM6a *spo11*Δ/CBS999.97(MAT1-1) *spo11*Δ ascus | ILLUMINA NextSeq 500 | PRJNA433292 | SRR6884707 |
| #5-5 | The #5 ascospore in the #5 QM6a *spo11*Δ/CBS999.97(MAT1-1) *spo11*Δ ascus | ILLUMINA NextSeq 500 | PRJNA433292 | SRR6884704 |
| #5-6 | The #6 ascospore in the #5 QM6a *spo11*Δ/CBS999.97(MAT1-1) *spo11*Δ ascus | ILLUMINA NextSeq 500 | PRJNA433292 | SRR6884705 |

| | | | | |
|---|---|---|---|---|
| #5-7 | The #7 ascospore in the #5 QM6a *spo11*Δ/CBS999.97(MAT1-1) *spo11*Δ ascus | ILLUMINA NextSeq 500 | PRJNA433292 | SRR6884712 |
| #5-8 | The #8 ascospore in the #5 QM6a *spo11*Δ/CBS999.97(MAT1-1) *spo11*Δ ascus | ILLUMINA NextSeq 500 | PRJNA433292 | SRR6884713 |
| #5-9 | The #9 ascospore in the #5 QM6a *spo11*Δ/CBS999.97(MAT1-1) *spo11*Δ ascus | ILLUMINA NextSeq 500 | PRJNA433292 | SRR6884699 |
| #5-10 | The #10 ascospore in the #5 QM6a *spo11*Δ/CBS999.97(MAT1-1) *spo11*Δ ascus | ILLUMINA NextSeq 500 | PRJNA433292 | SRR6884698 |
| #5-11 | The #11 ascospore in the #5 QM6a *spo11*Δ/CBS999.97(MAT1-1) *spo11*Δ ascus | ILLUMINA NextSeq 500 | PRJNA433292 | SRR6884701 |
| #5-12 | The #12 ascospore in the #5 QM6a *spo11*Δ/CBS999.97(MAT1-1) *spo11*Δ ascus | ILLUMINA NextSeq 500 | PRJNA433292 | SRR6884700 |
| #5-13 | The #13 ascospore in the #5 QM6a *spo11*Δ/CBS999.97(MAT1-1) *spo11*Δ ascus | ILLUMINA NextSeq 500 | PRJNA433292 | SRR6884695 |
| #5-14 | The #14 ascospore in the #5 QM6a *spo11*Δ/CBS999.97(MAT1-1) *spo11*Δ ascus | ILLUMINA NextSeq 500 | PRJNA433292 | SRR6884694 |
| #5-15 | The #15 ascospore in the #5 QM6a *spo11*Δ/CBS999.97(MAT1-1) *spo11*Δ ascus | ILLUMINA NextSeq 500 | PRJNA433292 | SRR6884697 |

| | | | | |
|---|---|---|---|---|
| #5-16 | The #16 ascospore in the #5 QM6a *spo11*Δ/CBS999.97(MAT1-1) *spo11*Δ ascus | ILLUMINA NextSeq 500 | PRJNA433292 | SRR6884696 |
| #6-1 | The #1 ascospore in the #6 QM6a *spo11*Δ/CBS999.97(MAT1-1) *spo11*Δ ascus | ILLUMINA NextSeq 500 | PRJNA433292 | SRR6884703 |
| #6-2 | The #2 ascospore in the #6 QM6a *spo11*Δ/CBS999.97(MAT1-1) *spo11*Δ ascus | ILLUMINA NextSeq 500 | PRJNA433292 | SRR6884702 |
| #6-3 | The #3 ascospore in the #6 QM6a *spo11*Δ/CBS999.97(MAT1-1) *spo11*Δ ascus | ILLUMINA NextSeq 500 | PRJNA433292 | SRR6884686 |
| #6-4 | The #4 ascospore in the #6 QM6a *spo11*Δ/CBS999.97(MAT1-1) *spo11*Δ ascus | ILLUMINA NextSeq 500 | PRJNA433292 | SRR6884687 |
| #6-5 | The #5 ascospore in the #6 QM6a *spo11*Δ/CBS999.97(MAT1-1) *spo11*Δ ascus | ILLUMINA NextSeq 500 | PRJNA433292 | SRR6884688 |
| #6-6 | The #6 ascospore in the #6 QM6a *spo11*Δ/CBS999.97(MAT1-1) *spo11*Δ ascus | ILLUMINA NextSeq 500 | PRJNA433292 | SRR6884689 |
| #6-7 | The #7 ascospore in the #6 QM6a *spo11*Δ/CBS999.97(MAT1-1) *spo11*Δ ascus | ILLUMINA NextSeq 500 | PRJNA433292 | SRR6884690 |
| #6-8 | The #8 ascospore in the #6 QM6a *spo11*Δ/CBS999.97(MAT1-1) *spo11*Δ ascus | ILLUMINA NextSeq 500 | PRJNA433292 | SRR6884691 |

| | | | | |
|---|---|---|---|---|
| #6-9 | The #9 ascospore in the #6 QM6a *spo11*Δ/CBS999.97(MAT1-1) *spo11*Δ ascus | ILLUMINA NextSeq 500 | PRJNA433292 | SRR6884692 |
| #6-10 | The #10 ascospore in the #6 QM6a *spo11*Δ/CBS999.97(MAT1-1) *spo11*Δ ascus | ILLUMINA NextSeq 500 | PRJNA433292 | SRR6884693 |
| #6-11 | The #11 ascospore in the #6 QM6a *spo11*Δ/CBS999.97(MAT1-1) *spo11*Δ ascus | ILLUMINA NextSeq 500 | PRJNA433292 | SRR6884684 |
| #6-12 | The #12 ascospore in the #6 QM6a *spo11*Δ/CBS999.97(MAT1-1) *spo11*Δ ascus | ILLUMINA NextSeq 500 | PRJNA433292 | SRR6884685 |
| #6-13 | The #13 ascospore in the #6 QM6a *spo11*Δ/CBS999.97(MAT1-1) *spo11*Δ ascus | ILLUMINA NextSeq 500 | PRJNA433292 | SRR6884683 |
| #6-14 | The #14 ascospore in the #6 QM6a *spo11*Δ/CBS999.97(MAT1-1) *spo11*Δ ascus | ILLUMINA NextSeq 500 | PRJNA433292 | SRR6884682 |
| #6-15 | The #15 ascospore in the #6 QM6a *spo11*Δ/CBS999.97(MAT1-1) *spo11*Δ ascus | ILLUMINA NextSeq 500 | PRJNA433292 | SRR6884681 |
| #6-16 | The #16 ascospore in the #6 QM6a *spo11*Δ/CBS999.97(MAT1-1) *spo11*Δ ascus | ILLUMINA NextSeq 500 | PRJNA433292 | SRR6884680 |
| #7-1 | The #1 ascospore in the #7 QM6a *spo11*Δ/CBS999.97(MAT1-1) *spo11*Δ ascus | ILLUMINA NextSeq 500 | PRJNA433292 | SRR6884677 |

| | | | | |
|---|---|---|---|---|
| #7-5 | The #5 ascospore in the #7 QM6a *spo11*Δ/CBS999.97(MAT1-1) *spo11*Δ ascus | ILLUMINA NextSeq 500 | PRJNA433292 | SRR6884676 |
| #7-9 | The #9 ascospore in the #7 QM6a *spo11*Δ/CBS999.97(MAT1-1) *spo11*Δ ascus | ILLUMINA NextSeq 500 | PRJNA433292 | SRR6884675 |
| #7-13 | The #13 ascospore in the #7 QM6a *spo11*Δ/CBS999.97(MAT1-1) *spo11*Δ ascus | ILLUMINA NextSeq 500 | PRJNA433292 | SRR6884674 |
| #8-1 | The #1 ascospore in the #8 QM6a *spo11*Δ/CBS999.97(MAT1-1) *spo11*Δ ascus | ILLUMINA NextSeq 500 | PRJNA433292 | SRR6884658 |
| #8-5 | The #5 ascospore in the #7 QM6a *spo11*Δ/CBS999.97(MAT1-1) *spo11*Δ ascus | ILLUMINA NextSeq 500 | PRJNA433292 | SRR6884659 |
| #8-9 | The #9 ascospore in the #8 QM6a *spo11*Δ/CBS999.97(MAT1-1) *spo11*Δ ascus | ILLUMINA NextSeq 500 | PRJNA433292 | SRR6884656 |
| #8-13 | The #13 ascospore in the #8 QM6a *spo11*Δ/CBS999.97(MAT1-1) *spo11*Δ ascus | ILLUMINA NextSeq 500 | PRJNA433292 | SRR6884657 |
| #9-1 | The #1 ascospore in the #9 QM6a *spo11*Δ/CBS999.97(MAT1-1) *spo11*Δ ascus | ILLUMINA NextSeq 500 | PRJNA433292 | SRR6884662 |
| #9-5 | The #5 ascospore in the #9 QM6a *spo11*Δ/CBS999.97(MAT1-1) *spo11*Δ ascus | ILLUMINA NextSeq 500 | PRJNA433292 | SRR6884663 |

| | | | | |
|---|---|---|---|---|
| #9-9 | The #9 ascospore in the #9 QM6a *spo11*Δ/CBS999.97(MAT1-1) *spo11*Δ ascus | ILLUMINA NextSeq 500 | PRJNA433292 | SRR6884660 |
| #9-13 | The #13 ascospore in the #9 QM6a *spo11*Δ/CBS999.97(MAT1-1) *spo11*Δ ascus | ILLUMINA NextSeq 500 | PRJNA433292 | SRR6884661 |

**Table S11. Summary of all CO products from the three asci (#1-#3) generated by crossing QM6a and CBS999.97(*MAT1-1*)**

| Asci number | Chromosome number | Chromatids | SNP_start_out | SNP_start_in | SNP_end_in | SNP_end_out | Position of CO | GC tract associated with CO | GC tract length (bp) | CO length (bp) |
|---|---|---|---|---|---|---|---|---|---|---|
| 1 | 1 | [1, 4] | 1376466 | 1376630 | 1376220 | 1376316 | 1376408 | yes | 280 | 280 |
| 1 | 1 | [2, 4] | 5942600 | 5942621 | 5942621 | 5942690 | 5942633 | yes | 45 | 45 |
| 1 | 1 | [1, 3] | 4850040 | 4850203 | 4850040 | 4850203 | 4850122 | no | 0 | 163 |
| 1 | 1 | [3, 4] | 2846520 | 2846712 | 2846520 | 2846712 | 2846616 | no | 0 | 192 |
| 1 | 1 | [1, 4] | 290360 | 290417 | 290660 | 290742 | 290545 | yes | 313 | 313 |
| 1 | 2 | [3, 4] | 1078135 | 1078229 | 1078313 | 1078580 | 1078314 | yes | 265 | 265 |
| 1 | 2 | [1, 2] | 3067988 | 3068061 | 3067677 | 3067793 | 3067880 | yes | 290 | 290 |
| 1 | 2 | [1, 2] | 4581140 | 4581413 | 4581140 | 4581413 | 4581277 | no | 0 | 273 |
| 1 | 3 | [1, 3] | 2230660 | 2231098 | 2230660 | 2231098 | 2230879 | no | 0 | 438 |
| 1 | 3 | [1, 4] | 389234 | 389285 | 389129 | 389234 | 389221 | yes | 78 | 78 |
| 1 | 3 | [2, 4] | 1639259 | 1639497 | 1639199 | 1639259 | 1639304 | yes | 149 | 149 |
| 1 | 4 | [1, 2] | 3937229 | 3937528 | 3937528 | 3937598 | 3937471 | yes | 185 | 185 |
| 1 | 4 | [1, 3] | 823701 | 823776 | 823701 | 823776 | 823739 | no | 0 | 75 |
| 1 | 4 | [1, 2] | 1922088 | 1922223 | 1921803 | 1921929 | 1922011 | yes | 290 | 290 |
| 1 | 5 | [3, 4] | 906203 | 906347 | 906347 | 906371 | 906317 | yes | 84 | 84 |

| | | | | | | | | | | |
|---|---|---|---|---|---|---|---|---|---|---|
| 1 | 5 | [3, 4] | 1651260 | 1651328 | 1651328 | 1651483 | 1651350 | yes | 112 | 112 |
| 1 | 5 | [2, 3] | 2640809 | 2641088 | 2640737 | 2640809 | 2640861 | yes | 176 | 176 |
| 1 | 5 | [1, 3] | 3964850 | 3965199 | 3964850 | 3965199 | 3965025 | no | 0 | 349 |
| 1 | 6 | [2, 4] | 2245618 | 2246094 | 2245618 | 2246094 | 2245856 | no | 0 | 476 |
| 1 | 6 | [1, 3] | 3265860 | 3266018 | 3266018 | 3266194 | 3266023 | yes | 167 | 167 |
| 1 | 6 | [1, 2] | 640643 | 640693 | 640864 | 641090 | 640823 | yes | 309 | 309 |
| 1 | 7 | [1, 2] | 2890900 | 2890971 | 2891551 | 2891642 | 2891266 | yes | 661 | 661 |
| 1 | 7 | [3, 4] | 1180585 | 1180958 | 1180585 | 1180958 | 1180772 | no | 0 | 373 |
| 1 | 7 | [3, 4] | 3411387 | 3411435 | 3411632 | 3411692 | 3411537 | yes | 251 | 251 |
| 2 | 1 | [2, 3] | 4251294 | 4251404 | 4251501 | 4251936 | 4251534 | yes | 370 | 370 |
| 2 | 1 | [1, 2] | 442558 | 442580 | 442608 | 442895 | 442660 | yes | 183 | 183 |
| 2 | 1 | [1, 2] | 5974725 | 5974980 | 5974725 | 5974980 | 5974853 | no | 0 | 255 |
| 2 | 1 | [1, 3] | 2084691 | 2084737 | 2084691 | 2084737 | 2084714 | no | 0 | 46 |
| 2 | 1 | [1, 2] | 1506474 | 1506720 | 1506745 | 1506803 | 1506686 | yes | 177 | 177 |
| 2 | 2 | [1, 2] | 4300767 | 4301025 | 4300364 | 4300510 | 4300667 | yes | 459 | 459 |
| 2 | 2 | [2, 4] | 1674268 | 1674949 | 1674268 | 1674949 | 1674609 | no | 0 | 681 |
| 2 | 2 | [2, 3] | 5020085 | 5020184 | 5019921 | 5020071 | 5020065 | yes | 139 | 139 |
| 2 | 2 | [2, 4] | 3360922 | 3361041 | 3361146 | 3361272 | 3361095 | yes | 228 | 228 |
| 2 | 3 | [3, 4] | 3159300 | 3159525 | 3159300 | 3159525 | 3159413 | no | 0 | 225 |
| 2 | 3 | [1, 3] | 1423196 | 1423241 | 1423196 | 1423241 | 1423219 | no | 0 | 45 |
| 2 | 3 | [1, 2] | 514260 | 514334 | 514359 | 514393 | 514337 | yes | 79 | 79 |
| 2 | 3 | [1, 3] | 2307852 | 2308078 | 2307631 | 2307774 | 2307834 | yes | 263 | 263 |

| | | | | | | | | | | |
|---|---|---|---|---|---|---|---|---|---|---|
| 2 | 4 | [2, 4] | 2627285 | 2627293 | 2627636 | 2627691 | 2627476 | yes | 375 | 375 |
| 2 | 4 | [2, 4] | 575839 | 575922 | 575839 | 575922 | 575881 | no | 0 | 83 |
| 2 | 4 | [2, 4] | 4085711 | 4085902 | 4085543 | 4085711 | 4085717 | yes | 180 | 180 |
| 2 | 5 | [1, 3] | 745152 | 745275 | 745152 | 745275 | 745214 | no | 0 | 123 |
| 2 | 5 | [1, 2] | 3666820 | 3666825 | 3666742 | 3666820 | 3666802 | yes | 42 | 42 |
| 2 | 5 | [1, 3] | 1724856 | 1725224 | 1724856 | 1725224 | 1725040 | no | 0 | 368 |
| 2 | 6 | [2, 4] | 3013333 | 3013564 | 3012895 | 3012955 | 3013187 | yes | 524 | 524 |
| 2 | 6 | [2, 3] | 833048 | 833157 | 832423 | 832694 | 832831 | yes | 544 | 544 |
| 2 | 7 | [3, 4] | 3111332 | 3111536 | 3110921 | 3111076 | 3111216 | yes | 436 | 436 |
| 2 | 7 | [2, 3] | 2211545 | 2211672 | 2211545 | 2211672 | 2211609 | no | 0 | 127 |
| 2 | 7 | [2, 3] | 1243420 | 1243724 | 1243241 | 1243413 | 1243450 | yes | 245 | 245 |
| 3 | 1 | [1, 3] | 2704304 | 2704336 | 2704501 | 2704569 | 2704428 | yes | 215 | 215 |
| 3 | 1 | [1, 2] | 801334 | 801348 | 801334 | 801348 | 801341 | no | 0 | 14 |
| 3 | 1 | [2, 4] | 5831517 | 5831622 | 5831664 | 5831925 | 5831682 | yes | 225 | 225 |
| 3 | 1 | [2, 3] | 250116 | 250612 | 250965 | 251305 | 250750 | yes | 771 | 771 |
| 3 | 2 | [2, 3] | 1673723 | 1673857 | 1673488 | 1673564 | 1673658 | yes | 264 | 264 |
| 3 | 2 | [2, 3] | 1529079 | 1529171 | 1518542 | 1518568 | 1523840 | yes | 10570 | 10570 |
| 3 | 2 | [1, 3] | 422994 | 423160 | 423367 | 423632 | 423288 | yes | 423 | 423 |
| 3 | 2 | [2, 3] | 3335826 | 3336217 | 3335826 | 3336217 | 3336022 | no | 0 | 391 |
| 3 | 2 | [2, 3] | 2537986 | 2538219 | 2537835 | 2537976 | 2538004 | yes | 197 | 197 |
| 3 | 2 | [2, 3] | 5223023 | 5223642 | 5223023 | 5223642 | 5223333 | no | 0 | 619 |
| 3 | 3 | [1, 2] | 537260 | 537458 | 537260 | 537458 | 537359 | no | 0 | 198 |

| | | | | | | | | | | |
|---|---|---|---|---|---|---|---|---|---|---|
| 3 | 3 | [2, 4] | 1395209 | 1395432 | 1395209 | 1395432 | 1395321 | no | 0 | 223 |
| 3 | 3 | [1, 3] | 2086346 | 2086547 | 2085628 | 2085849 | 2086093 | yes | 708 | 708 |
| 3 | 4 | [1, 4] | 557846 | 558182 | 557846 | 558182 | 558014 | no | 0 | 336 |
| 3 | 4 | [1, 2] | 2676737 | 2676891 | 2676737 | 2676891 | 2676814 | no | 0 | 154 |
| 3 | 4 | [1, 4] | 1902383 | 1902552 | 1901961 | 1902383 | 1902320 | yes | 296 | 296 |
| 3 | 4 | [1, 2] | 3558403 | 3558505 | 3557186 | 3557217 | 3557828 | yes | 1253 | 1253 |
| 3 | 5 | [1, 3] | 2923773 | 2923934 | 2923997 | 2924060 | 2923941 | yes | 175 | 175 |
| 3 | 5 | [2, 4] | 3952992 | 3953332 | 3952992 | 3953332 | 3953162 | no | 0 | 340 |
| 3 | 5 | [2, 3] | 776022 | 776148 | 775763 | 775830 | 775941 | yes | 289 | 289 |
| 3 | 5 | [2, 4] | 1672597 | 1672807 | 1672372 | 1672594 | 1672593 | yes | 219 | 219 |
| 3 | 6 | [2, 4] | 2581058 | 2581227 | 2581227 | 2581576 | 2581272 | yes | 259 | 259 |
| 3 | 6 | [2, 4] | 1444127 | 1444280 | 1444280 | 1444692 | 1444345 | yes | 283 | 283 |
| 3 | 6 | [2, 4] | 3091487 | 3091546 | 3090941 | 3091107 | 3091270 | yes | 493 | 493 |
| 3 | 6 | [2, 4] | 600880 | 600918 | 600933 | 601023 | 600939 | yes | 79 | 79 |
| 3 | 7 | [1, 3] | 1092936 | 1093155 | 1092699 | 1092936 | 1092932 | yes | 228 | 228 |
| 3 | 7 | [2, 3] | 3110158 | 3110271 | 3109955 | 3110158 | 3110136 | yes | 158 | 158 |

**Table S12. Summary of all NCO products from the three asci (#1-#3) generated by crossing QM6a and CBS999.97(*MAT1-1*)**

| Asci number | Chromosome number | Chromatids | SNP_start_out | SNP_start_in | SNP_end_in | SNP_end_out | Position of NCO (bp) | SNP number | NCO length |
|---|---|---|---|---|---|---|---|---|---|
| 1 | 1 | 4 | 2841732 | 2841773 | 2841798 | 2842041 | 2841836 | 2 | 167 |
| 1 | 2 | 1 | 1379241 | 1379363 | 1379405 | 1379528 | 1379384 | 3 | 165 |
| 1 | 3 | 3 | 1308011 | 1308088 | 1308375 | 1308402 | 1308219 | 22 | 339 |
| 1 | 3 | 2 | 1581077 | 1581212 | 1581214 | 1581307 | 1581203 | 2 | 116 |
| 1 | 3 | 2 | 3934647 | 3934821 | 3935580 | 3935970 | 3935255 | 13 | 1041 |
| 1 | 3 | 1 | 4101219 | 4101426 | 4101619 | 4101796 | 4101515 | 9 | 385 |
| 1 | 5 | 1 | 783355 | 783388 | 783475 | 783607 | 783456 | 2 | 170 |
| 1 | 5 | 1 | 2906100 | 2906222 | 2906235 | 2906367 | 2906231 | 3 | 140 |
| 2 | 1 | 3 | 1369629 | 1370048 | 1370056 | 1370333 | 1370017 | 2 | 356 |
| 2 | 1 | 2 | 3651786 | 3651791 | 3651837 | 3651856 | 3651818 | 3 | 58 |
| 2 | 1 | 4 | 3712241 | 3712365 | 3712549 | 3712728 | 3712471 | 12 | 336 |
| 2 | 2 | 1 | 1084839 | 1084945 | 1085158 | 1085167 | 1085027 | 8 | 271 |
| 2 | 2 | 4 | 4571606 | 4571666 | 4571815 | 4571994 | 4571770 | 9 | 269 |
| 2 | 2 | 3 | 5129211 | 5129412 | 5129467 | 5129695 | 5129446 | 5 | 270 |
| 2 | 3 | 4 | 1577457 | 1577642 | 1577672 | 1577695 | 1577617 | 2 | 134 |
| 2 | 3 | 4 | 2103630 | 2103799 | 2103989 | 2104176 | 2103899 | 11 | 368 |

| | | | | | | | | | |
|---|---|---|---|---|---|---|---|---|---|
| 2 | 3 | 2 | 2431447 | 2431638 | 2432263 | 2432791 | 2432035 | 12 | 985 |
| 2 | 4 | 2 | 1138845 | 1138905 | 1139077 | 1139160 | 1138997 | 21 | 244 |
| 2 | 4 | 4 | 1724988 | 1725687 | 1726330 | 1726778 | 1725946 | 5 | 1217 |
| 2 | 6 | 1 | 1406831 | 1406962 | 1409630 | 1409834 | 1408314 | 104 | 2836 |
| 2 | 7 | 1 | 2140902 | 2140949 | 2141212 | 2141509 | 2141143 | 6 | 435 |
| 3 | 1 | 3 | 1958151 | 1958389 | 1958402 | 1958440 | 1958346 | 4 | 151 |
| 3 | 1 | 4 | 250965 | 251305 | 258564 | 258641 | 254869 | 190 | 7468 |
| 3 | 2 | 2 | 276341 | 276357 | 276394 | 276455 | 276387 | 3 | 76 |
| 3 | 2 | 4 | 1081240 | 1081282 | 1081314 | 1081467 | 1081326 | 3 | 130 |
| 3 | 2 | 3 | 1515769 | 1515902 | 1517414 | 1517551 | 1516659 | 36 | 1647 |
| 3 | 2 | 1 | 4581140 | 4581413 | 4582518 | 4582923 | 4581999 | 13 | 1444 |
| 3 | 3 | 1 | 2083858 | 2084158 | 2084730 | 2084855 | 2084400 | 32 | 785 |
| 3 | 3 | 1 | 2086583 | 2086838 | 2086932 | 2087078 | 2086858 | 2 | 295 |
| 3 | 3 | 1 | 4830502 | 4831120 | 4831818 | 4832061 | 4831375 | 22 | 1129 |
| 3 | 5 | 2 | 162435 | 162635 | 166837 | 166915 | 164706 | 54 | 4341 |
| 3 | 6 | 1 | 1263014 | 1263092 | 1263164 | 1263243 | 1263128 | 2 | 151 |
| 3 | 6 | 4 | 2581995 | 2582082 | 2582161 | 2582218 | 2582114 | 4 | 151 |
| 3 | 7 | 3 | 532552 | 532597 | 532612 | 532631 | 532598 | 2 | 47 |
| 3 | 7 | 3 | 714501 | 714627 | 714682 | 714777 | 714647 | 4 | 166 |
| 3 | 7 | 3 | 2432818 | 2433100 | 2433281 | 2433337 | 2433134 | 9 | 350 |

**Table S13. PacBio sequencing and assembly results of the four representative F1 progeny in the #1 ascus and the #4 ascus**

| Strain[1] | #1-1 | #1-5 | #1-9 | #1-13 |
|---|---|---|---|---|
| Sequencing method | PacBio RSII | PacBio RSII | PacBio RSII | PacBio RSII |
| Total sequenced bases | 2,078,510,561 bp | 4,810,795,008 bp | 6,202,644,396 bp | 5,732,223,364 bp |
| Analysis application | Assembly (HGAP3) | Assembly (HGAP3) | Assembly (HGAP3) | Assembly (HGAP3) |
| Maximum of all assembled unitigs | 6,801,106 bp | 6,822,671 bp | 6,835,650 bp | 6,849,886 bp |
| $N_{50}$ of all assembled unitigs | 5,275,058 bp | 5,258,125 bp | 5,262,578 bp | 4,872,978 bp |
| Number reads | 170,751 | 370,345 | 636,376 | 386,388 |
| $N_{50}$ reads | 17,254 bp | 18,599 bp | 14,406 bp | 21,047 bp |
| Phred quality score | 48.8 | 48.1 | 48.7 | 48.8 |

| | | | | | | | | |
|---|---|---|---|---|---|---|---|---|
| Unitigs | 7 linear chromosomes | | 7 linear chromosomes | | 7 linear chromosomes | | 7 linear chromosomes | |
| Coverage | 55 X | | 93 X | | 127 X | | 113 X | |
| Genome size | 34,922,528 bp | | 34,319,199 bp | | 34,324,311 bp | | 34,434,741 bp | |
| Unidentified bases (N) | 0 bp | | 0 bp | | 0 bp | | 0 bp | |
| GC content | 51.1% | | 51.6% | | 51.6% | | 51.5% | |
| **[2]Strain** | **#4-1** | | **#4-5** | | **#4-9** | | **#4-13** | |
| Sequencing method | PacBio sequel | | PacBio sequel | | PacBio sequel | | PacBio sequel | |
| Total sequenced bases | 4,625,221,270 | | 4,373,478,251 | | 2,756,929,979 | | 4,253,956,984 | |
| Analysis application | Resequence | | Resequence | | Resequence | | Resequence | |
| Reference genomic sequences | QM6a | CBS999.97 (MAT1-1) | QM6a | CBS999.97 (MAT1-1) | QM6a | CBS999.97 (MAT1-1) | QM6a | CBS999.97 (MAT1-1) |
| Reference consensus concordance (mean) | 98.17% | 99% | 98.46% | 98.72% | 98.57% | 98.73% | 99.10% | 98.06% |

| | | | | | | | | |
|---|---|---|---|---|---|---|---|---|
| Mean concordance (mapped) | 87.7% | 88.4% | 88.1% | 88.4% | 86.2% | 86.2% | 88.5% | 87.5% |
| Number of subreads (mapped) | 578,256 | 587,540 | 568,283 | 570,155 | 292,937 | 293,313 | 540,750 | 525,680 |
| Number of subread bases (mapped) | 3,863,155,720 | 3,981,737,173 | 3,731,450,212 | 3,757,452,841 | 1,833,026,510 | 1,823,693,606 | 3,617,439,353 | 3,443,611,790 |
| Mean coverage | 109X | 114X | 105X | 107X | 50X | 51X | 101X | 97X |
| Genome size | 34,924,315 | 34,328,149 | 34926061 | 34333358 | 34,920,815 | 34,342,449 | 34922714 | 34344368 |

1. The genome of the four representative ascospores from ascus #1 were sequenced by the PacBio RSII sequencing system and then assembled by the HGAP3 assembly protocol (SMRTportal 2.2.0).
2. The genomic sequences of the four representative ascospores from ascus #4 were sequenced by the PacBio Sequel sequencing system and then assembled by the Resequencing application, which maps sequencing reads against two reference sequences: QM6a and CBS999.97(MAT1-1) following the user guide (https://www.pacb.com/wp-content/uploads/SMRT_Link_User_Guide_v600.pdf).

### Table S14. Distribution of CO and NCO products in the seven homologous chromosomes

| | QM6a X CBS99.97(*MAT1-1*) | | | | | | | QM6a *spo11*Δ X CBS99.97(*MAT1-1*) *spo11*Δ | | | | | | | QM6a *spo11*Δ X CBS99.97(*MAT1-1*) *spo11*Δ | | | | | | |
|---|---|---|---|---|---|---|---|---|---|---|---|---|---|---|---|---|---|---|---|---|---|
| Chromosome | SNP coverage | Asci #1 | | Asci #2 | | Asci #3 | | SNP coverage | Asci #4 | | Asci #5 | | Asci #6 | | SNP coverage | Asci #7 | | Asci #8 | | Asci #9 | |
| | | CO | NCO | CO | NCO | CO | NCO | | CO | NCO | CO | NCO | CO | NCO | | CO | NCO | CO | NCO | CO | NCO |
| I | 100% | 5 | 1 | 5 | 3 | 4 | 2 | 92.86% | 4 | 2 | 1 | 0 | 5 | 3 | 26.86% | 1 | 0 | 1 | 0 | 2 | 1 |
| II | 100% | 3 | 1 | 4 | 3 | 6 | 4 | 97.79% | 3 | 0 | 2 | 0 | 4 | 3 | 47.87% | 2 | 2 | 1 | 0 | 3 | 0 |
| III | 100% | 3 | 4 | 4 | 3 | 3 | 3 | 94.50% | 2 | 2 | 2 | 1 | 1 | 4 | 72.76% | 0 | 1 | 1 | 0 | 3 | 1 |
| IV | 100% | 3 | 0 | 3 | 2 | 4 | 0 | 100% | 3 | 1 | 2 | 0 | 5 | 1 | 81.08% | 0 | 2 | 3 | 0 | 4 | 1 |
| V | 100% | 4 | 2 | 3 | 0 | 4 | 1 | 100% | 4 | 1 | 2 | 0 | 2 | 0 | 28.13% | 1 | 1 | 0 | 0 | 2 | 0 |
| VI | 100% | 3 | 0 | 2 | 1 | 4 | 2 | 100% | 2 | 0 | 1 | 0 | 2 | 0 | 77.68% | 1 | 0 | 1 | 2 | 4 | 1 |
| VII | 100% | 3 | 0 | 3 | 1 | 2 | 3 | 77.34% | 1 | 1 | 2 | 0 | 2 | 2 | 25.65% | 0 | 0 | 0 | 2 | 1 | 1 |
| I-VII | 100% | 24 | 8 | 24 | 13 | 27 | 15 | 94.64% | 19 | 7 | 12 | 1 | 21 | 13 | 50.51% | 5 | 6 | 7 | 4 | 19 | 5 |
| Total | 75 COs and 36 NCOs | | | | | | | 52 COs and 21 NCOs | | | | | | | 31 COs and 15 NCOs | | | | | | |

**Table S15. Summary of all CO products from the three asci (#4-#6) generated by crossing the first pair of QM6a *spo11*Δ and CBS999.97(*MAT1-1*) *spo11*Δ mutants**

| Asci number | Chromosome number | Chromatids | SNP_start_out | SNP_start_in | SNP_end_in | SNP_end_out | Position of CO | GC tract associated with CO | GC tract length (bp) | CO length (bp) |
|---|---|---|---|---|---|---|---|---|---|---|
| 1 | 1 | [1, 4] | 2241842 | 2241952 | 2241769 | 2241842 | 2241851 | yes | 92 | 92 |
| 1 | 1 | [1, 2] | 6409743 | 6409839 | 6409743 | 6409839 | 6409791 | no | 0 | 96 |
| 1 | 1 | [2, 3] | 4763716 | 4763778 | 4763564 | 4763668 | 4763682 | yes | 131 | 131 |
| 1 | 1 | [1, 2] | 1769653 | 1769855 | 1769653 | 1769855 | 1769754 | no | 0 | 202 |
| 1 | 2 | [2, 4] | 2897929 | 2898274 | 2897761 | 2897929 | 2897973 | yes | 257 | 257 |
| 1 | 2 | [2, 4] | 2156378 | 2156697 | 2156378 | 2156697 | 2156538 | no | 0 | 319 |
| 1 | 2 | [1, 3] | 1103549 | 1104399 | 1103485 | 1103534 | 1103742 | yes | 465 | 465 |
| 1 | 3 | [2, 3] | 992357 | 992518 | 991973 | 992075 | 992231 | yes | 414 | 414 |
| 1 | 3 | [2, 4] | 2827908 | 2828034 | 2827908 | 2828034 | 2827971 | no | 0 | 126 |
| 1 | 4 | [2, 4] | 4020005 | 4020052 | 4020052 | 4020124 | 4020058 | yes | 60 | 60 |
| 1 | 4 | [1, 3] | 1242923 | 1242937 | 1242957 | 1243008 | 1242956 | yes | 53 | 53 |
| 1 | 4 | [1, 3] | 3095802 | 3095844 | 3095682 | 3095802 | 3095783 | yes | 81 | 81 |
| 1 | 5 | [1, 2] | 2588226 | 2588267 | 2588267 | 2588388 | 2588287 | yes | 81 | 95 |
| 1 | 5 | [1, 4] | 519154 | 519238 | 519239 | 519377 | 519252 | yes | 112 | 112 |
| 1 | 5 | [3, 4] | 4088627 | 4088927 | 4088627 | 4088927 | 4088777 | no | 0 | 300 |

| | | | | | | | | | | |
|---|---|---|---|---|---|---|---|---|---|---|
| 1 | 5 | [2, 4] | 1690104 | 1690199 | 1690104 | 1690199 | 1690152 | no | 0 | 95 |
| 1 | 6 | [2, 3] | 3269980 | 3270115 | 3270124 | 3270220 | 3270110 | yes | 125 | 125 |
| 1 | 6 | [1, 3] | 244261 | 244399 | 244261 | 244399 | 244330 | no | 0 | 138 |
| 1 | 7 | [1, 3] | 1283745 | 1283835 | 1283464 | 1283586 | 1283658 | yes | 265 | 265 |
| 2 | 1 | [1, 3] | 2735360 | 2735689 | 2735360 | 2735689 | 2735525 | no | 0 | 329 |
| 2 | 2 | [1, 3] | 2539850 | 2540457 | 2539850 | 2540457 | 2540154 | no | 0 | 607 |
| 2 | 2 | [3, 4] | 3265637 | 3403422 | 3265637 | 3403422 | 3334530 | no | 0 | 137785 |
| 2 | 3 | [2, 4] | 4879455 | 4879922 | 4879455 | 4879922 | 4879689 | no | 0 | 467 |
| 2 | 3 | [1, 3] | 1482361 | 1482496 | 1482361 | 1482496 | 1482429 | no | 0 | 135 |
| 2 | 4 | [1, 4] | 1240981 | 1241185 | 1241191 | 1241259 | 1241154 | yes | 142 | 142 |
| 2 | 4 | [2, 3] | 3145084 | 3145281 | 3145281 | 3145308 | 3145239 | yes | 112 | 112 |
| 2 | 5 | [1, 4] | 3164269 | 3164357 | 3163568 | 3163905 | 3164025 | yes | 577 | 577 |
| 2 | 5 | [2, 3] | 1802626 | 1803383 | 1802626 | 1803383 | 1803005 | no | 0 | 757 |
| 2 | 6 | [1, 3] | 1761028 | 1761172 | 1761355 | 1761396 | 1761238 | yes | 276 | 276 |
| 2 | 7 | [1, 3] | 2221582 | 2221752 | 2221930 | 2221964 | 2221807 | yes | 280 | 280 |
| 2 | 7 | [2, 4] | 2858910 | 2859253 | 2858910 | 2859253 | 2859082 | no | 0 | 343 |
| 3 | 1 | [2, 3] | 1774685 | 1774775 | 1774685 | 1774775 | 1774730 | no | 0 | 90 |
| 3 | 1 | [1, 4] | 5629095 | 5629308 | 5629313 | 5629496 | 5629303 | yes | 203 | 203 |
| 3 | 1 | [2, 3] | 2566063 | 2566316 | 2566652 | 2566878 | 2566477 | yes | 576 | 576 |
| 3 | 1 | [1, 4] | 6704547 | 6705209 | 6704547 | 6705209 | 6704878 | no | 0 | 662 |
| 3 | 1 | [1, 3] | 513739 | 513863 | 513681 | 513718 | 513750 | yes | 102 | 102 |
| 3 | 2 | [2, 3] | 2609281 | 2609462 | 2609095 | 2609281 | 2609280 | yes | 184 | 184 |

| | | | | | | | | | | |
|---|---|---|---|---|---|---|---|---|---|---|
| 3 | 2 | [3, 4] | 3901065 | 3901144 | 3901065 | 3901144 | 3901105 | no | 0 | 79 |
| 3 | 2 | [2, 3] | 1870797 | 1870942 | 1870942 | 1871041 | 1870931 | yes | 122 | 122 |
| 3 | 2 | [1, 2] | 233783 | 233903 | 233330 | 233344 | 233590 | yes | 506 | 506 |
| 3 | 3 | [2, 3] | 1073712 | 1073823 | 1074212 | 1074273 | 1074005 | yes | 475 | 475 |
| 3 | 4 | [2, 4] | 4144762 | 4144780 | 4144713 | 4144756 | 4144753 | yes | 37 | 37 |
| 3 | 4 | [2, 4] | 1965377 | 1965613 | 1964912 | 1965131 | 1965258 | yes | 474 | 474 |
| 3 | 4 | [1, 3] | 2658234 | 2658268 | 2657912 | 2658234 | 2658162 | yes | 178 | 178 |
| 3 | 4 | [2, 4] | 1188245 | 1188529 | 1188529 | 1188823 | 1188532 | yes | 289 | 289 |
| 3 | 4 | [2, 4] | 560791 | 560980 | 560791 | 560980 | 560886 | no | 0 | 189 |
| 3 | 5 | [1, 3] | 2591380 | 2591967 | 2596510 | 2596533 | 2594043 | yes | 4739 | 4739 |
| 3 | 5 | [2, 4] | 1894350 | 1894573 | 1894648 | 1894746 | 1894579 | yes | 236 | 236 |
| 3 | 6 | [1, 4] | 1586463 | 1586839 | 1586463 | 1586839 | 1586651 | no | 0 | 376 |
| 3 | 6 | [1, 2] | 642933 | 643765 | 643765 | 643892 | 643589 | yes | 480 | 480 |

**Table S16. Summary of all NCO products from the three asci (#4-#6) generated by crossing the first pair of QM6a *spo11*Δ and CBS999.97(*MAT1-1*) *spo11*Δ mutants**

| Asci number | Chromosome number | Chromatids | SNP_start_out | SNP_start_in | SNP_end_in | SNP_end_out | Position of NCO (bp) | SNP number | NCO length |
|---|---|---|---|---|---|---|---|---|---|
| 1 | 1 | 2 | 2853921 | 2854012 | 2854363 | 2854674 | 2854243 | 9 | 552 |
| 1 | 1 | 2 | 2960385 | 2960429 | 2960596 | 2960629 | 2960510 | 3 | 206 |
| 1 | 3 | 2 | 983870 | 984114 | 984116 | 984259 | 984090 | 2 | 196 |
| 1 | 3 | 1 | 2985663 | 2985838 | 2986713 | 2986852 | 2986267 | 13 | 1032 |
| 1 | 4 | 3 | 2770039 | 2770097 | 2770405 | 2771687 | 2770557 | 2 | 978 |
| 1 | 5 | 4 | 3154517 | 3154897 | 3156725 | 3156962 | 3155775 | 15 | 2137 |
| 1 | 7 | 4 | 2773139 | 2773571 | 2774433 | 2774456 | 2773900 | 23 | 1090 |
| 2 | 3 | 1 | 1482118 | 1482166 | 1482181 | 1482295 | 1482190 | 5 | 96 |
| 3 | 1 | 1 | 2041689 | 2041745 | 2042085 | 2042163 | 2041921 | 10 | 407 |
| 3 | 1 | 1 | 6123851 | 6124970 | 6124971 | 6125477 | 6124817 | 2 | 814 |
| 3 | 1 | 0 | 1472641 | 1472833 | 1472882 | 1472935 | 1472823 | 2 | 172 |
| 3 | 2 | 3 | 3912210 | 3912246 | 3912495 | 3912700 | 3912413 | 4 | 370 |
| 3 | 2 | 3 | 4548348 | 4548441 | 4548989 | 4549146 | 4548731 | 26 | 673 |
| 3 | 2 | 1 | 5637286 | 5637342 | 5637471 | 5637536 | 5637409 | 7 | 190 |
| 3 | 3 | 2 | 188381 | 188392 | 189196 | 189284 | 188813 | 28 | 854 |

| | | | | | | | | | |
|---|---|---|---|---|---|---|---|---|---|
| 3 | 3 | 3 | 1577409 | 1577457 | 1577695 | 1578150 | 1577678 | 4 | 490 |
| 3 | 3 | 4 | 2266423 | 2266718 | 2267648 | 2267736 | 2267131 | 76 | 1122 |
| 3 | 3 | 3 | 5158688 | 5159280 | 5159514 | 5159572 | 5159264 | 17 | 559 |
| 3 | 4 | 1 | 1457320 | 1457535 | 1457752 | 1457782 | 1457597 | 3 | 340 |
| 3 | 7 | 2 | 1664266 | 1664336 | 1664493 | 1664760 | 1664464 | 4 | 326 |
| 3 | 7 | 1 | 1795858 | 1797947 | 1798009 | 1798012 | 1797457 | 2 | 1108 |

**Table S17. Summary of all CO products from the three asci (#7-#9) generated by crossing the second pair of QM6a *spo11*Δ and CBS999.97(*MAT1-1*) *spo11*Δ mutants**

| Asci number | Chromosome number | Chromatids | SNP_start_out | SNP_start_in | SNP_end_in | SNP_end_out | Position of CO | GC tract associated with CO | GC tract length (bp) | CO length (bp) |
|---|---|---|---|---|---|---|---|---|---|---|
| 1 | 1 | [1/2, 4] | 1154888 | 1154970 | 1155297 | 1155354 | 1155127 | yes | 396.5 | 396.5 |
| 1 | 2 | [1, 4] | 917515 | 917675 | 918079 | 918313 | 917895.5 | yes | 601 | 601 |
| 1 | 2 | [1, 4] | 2176457 | 2176490 | 2176690 | 2176843 | 2176620 | yes | 293 | 293 |
| 1 | 5 | [2, 4] | 334710 | 334879 | 334915 | 335023 | 334881.8 | yes | 174.5 | 174.5 |
| 1 | 6 | [2, 4] | 1754113 | 1754223 | 1754113 | 1754223 | 1754168 | no | 0 | 110 |
| 2 | 1 | [2, 4] | 843695 | 844019 | 844019 | 844055 | 843947 | yes | 180 | 180 |
| 2 | 2 | [1, 4] | 2659364 | 2659535 | 2659589 | 2659603 | 2659523 | yes | 54 | 54 |
| 2 | 3 | [2, 3] | 2885922 | 2886135 | 2886189 | 2886386 | 2886158 | yes | 259 | 259 |
| 2 | 4 | [1, 4] | 676208 | 676495 | 676495 | 676587 | 676446.3 | yes | 189.5 | 189.5 |
| 2 | 4 | [2, 3] | 2536572 | 2536589 | 2536662 | 2536712 | 2536634 | yes | 106.5 | 106.5 |
| 2 | 4 | [1, 4] | 3247755 | 3248165 | 3247755 | 3248165 | 3247960 | no | 0 | 410 |
| 2 | 6 | [1, 3] | 1760954 | 1761028 | 1761093 | 1761136 | 1761053 | yes | 123.5 | 123.5 |
| 3 | 1 | [3, 4] | 1110333 | 1110538 | 1110614 | 1110839 | 1110581 | yes | 291 | 291 |
| 3 | 1 | [1, 4] | 5104165 | 5104180 | 5104268 | 5104306 | 5104230 | yes | 114.5 | 114.5 |
| 3 | 2 | [3, 4] | 1161153 | 1161369 | 1161420 | 1161582 | 1161381 | yes | 51 | 51 |

| | | | | | | | | | | |
|---|---|---|---|---|---|---|---|---|---|---|
| 3 | 2 | [2, 3] | 1583376 | 1583477 | 1583711 | 1583878 | 1583611 | yes | 234 | 234 |
| 3 | 2 | [2, 3] | 2350646 | 2351290 | 2350646 | 2351290 | 2350968 | no | 0 | 644 |
| 3 | 3 | [1, 3] | 2391079 | 2391271 | 2391395 | 2391875 | 2391405 | yes | 460 | 460 |
| 3 | 3 | [3, 4] | 4105417 | 4105631 | 4105631 | 4105730 | 4105602 | yes | 156.5 | 156.5 |
| 3 | 3 | [3, 4] | 4937795 | 4937885 | 4939804 | 4939958 | 4938861 | yes | 2041 | 2041 |
| 3 | 4 | [2, 3] | 2351665 | 2352304 | 2351665 | 2352304 | 2351985 | no | 0 | 639 |
| 3 | 4 | [2, 3] | 2617141 | 2617366 | 2620221 | 2620272 | 2618750 | yes | 2855 | 2855 |
| 3 | 4 | [2, 4] | 3310394 | 3310628 | 3310639 | 3310820 | 3310620 | yes | 218.5 | 218.5 |
| 3 | 4 | [3, 4] | 3936546 | 3936753 | 3936753 | 3936905 | 3936739 | yes | 179.5 | 179.5 |
| 3 | 5 | [2, 3] | 2588083 | 2588213 | 2588470 | 2588642 | 2588352 | yes | 257 | 257 |
| 3 | 5 | [1, 4] | 2748079 | 2748280 | 2748079 | 2748280 | 2748180 | no | 0 | 201 |
| 3 | 6 | [1, 2] | 486396 | 486451 | 486462 | 486580 | 486472.3 | yes | 97.5 | 97.5 |
| 3 | 6 | [1, 4] | 1713933 | 1714456 | 1714804 | 1714955 | 1714537 | yes | 685 | 685 |
| 3 | 6 | [1, 4] | 2761238 | 2761264 | 2761294 | 2761477 | 2761318 | yes | 134.5 | 134.5 |
| 3 | 6 | [2, 3] | 3257425 | 3257597 | 3258180 | 3258283 | 3257871 | yes | 720.5 | 720.5 |
| 3 | 7 | [3, 4] | 2619520 | 2619612 | 2622527 | 2622634 | 2621073 | yes | 3014.5 | 3014.5 |

**Table S18. Summary of all NCO products from the three asci (#7-#9) generated by crossing the 2nd pair of QM6a *spo11*Δ and CBS999.97(*MAT1-1*) *spo11*Δ mutants**

| Asci number | Chromosome number | Chromatids | SNP_start | SNP_end | Position of NCO (bp) | SNP number | NCO length (bp) |
|---|---|---|---|---|---|---|---|
| 7 | 2 | 1 | 924224 | 927097 | 925661 | 91 | 2873 |
| 7 | 2 | 1 | 1063734 | 1064152 | 1063943 | 6 | 418 |
| 7 | 3 | 3 | 4858574 | 4858852 | 4858713 | 3 | 278 |
| 7 | 4 | 1 | 4500742 | 4501971 | 4501357 | 9 | 1229 |
| 7 | 4 | 1 | 4502224 | 4502609 | 4502417 | 5 | 385 |
| 7 | 5 | 2 | 333621 | 333648 | 333635 | 2 | 27 |
| 8 | 6 | 1 | 1358929 | 1360969 | 1359949 | 58 | 2040 |
| 8 | 6 | 1 | 1361143 | 1361666 | 1361405 | 25 | 523 |
| 8 | 7 | 4 | 2757097 | 2758429 | 2757763 | 3 | 1332 |
| 8 | 7 | 3 | 3020104 | 3030476 | 3025290 | 9 | 10372 |
| 9 | 1 | 1 | 536146 | 536244 | 536195 | 11 | 98 |
| 9 | 3 | 2 | 4557879 | 4557929 | 4557904 | 3 | 50 |
| 9 | 4 | 2 | 2616194 | 2616303 | 2616249 | 3 | 109 |
| 9 | 6 | 3 | 1943827 | 1943975 | 1943901 | 5 | 148 |
| 9 | 7 | 3 | 2630426 | 2631625 | 2631026 | 22 | 1199 |

**Table S19. Summary of pairwise SNP calling between the 5th chromosomes# in all 16 F1 progeny of the #1 ascus generated by crossing QM6a and CBS999.97(*MAT1-1*)**

| ChV | 1-1 | 1-2 | 1-14 | 1-15 | 1-3 | 1-4 | 1-5 | 1-6 | 1-7 | 1-8 | 1-9 | 1-10 | 1-11 | 1-12 | 1-15 | 1-16 |
|---|---|---|---|---|---|---|---|---|---|---|---|---|---|---|---|---|
| 1-1 | | 1602 | 1494 | 1581 | 64973 | 64405 | 64319 | 64330 | 51865 | 51928 | 51830 | 51903 | 82379 | 82277 | 82195 | 82218 |
| 1-2 | | | 1476 | 1761 | 65039 | 64403 | 64407 | 64347 | 52001 | 51981 | 51960 | 52020 | 82459 | 82314 | 82294 | 82282 |
| 1-14 | | | | 1613 | 64894 | 64275 | 64281 | 64244 | 51783 | 51793 | 51767 | 51883 | 82511 | 82233 | 82246 | 82224 |
| 1-15 | | | | | 65100 | 64594 | 64485 | 64421 | 52103 | 52144 | 52091 | 52172 | 82583 | 82458 | 82351 | 82404 |
| 1-3 | | | | | | 2359 | 2307 | 2117 | 82953 | 83024 | 82972 | 83010 | 52755 | 52632 | 52680 | 52732 |
| 1-4 | | | | | | | 1796 | 1651 | 82446 | 82417 | 82435 | 82427 | 52200 | 51970 | 51975 | 52026 |
| 1-5 | | | | | | | | 1607 | 82291 | 82362 | 82336 | 82363 | 52200 | 51958 | 51918 | 52029 |
| 1-6 | | | | | | | | | 82416 | 82440 | 82380 | 82420 | 52184 | 52060 | 51911 | 52027 |
| 1-7 | | | | | | | | | | 1377 | 1467 | 1538 | 64414 | 64071 | 64051 | 64116 |
| 1-8 | | | | | | | | | | | 1487 | 1557 | 64408 | 64105 | 64034 | 64166 |
| 1-9 | | | | | | | | | | | | 1580 | 64362 | 64017 | 64077 | 64092 |
| 1-10 | | | | | | | | | | | | | 64457 | 64076 | 64109 | 64189 |
| 1-11 | | | | | | | | | | | | | | 1624 | 1629 | 1718 |
| 1-12 | | | | | | | | | | | | | | | 1347 | 1402 |
| 1-15 | | | | | | | | | | | | | | | | 1297 |
| 1-16 | | | | | | | | | | | | | | | | |

### The overall number of SNPs in the 5th chromosome is 115,253.

**Table S20. Summary of pairwise SNP calling between the 6th chromosomes# in all 16 F1 progeny of the #1 ascus generated by crossing QM6a and CBS999.97(*MAT1-1*)**

| ChVI | 1-1 | 1-2 | 1-14 | 1-15 | 1-3 | 1-4 | 1-5 | 1-6 | 1-7 | 1-8 | 1-9 | 1-10 | 1-11 | 1-12 | 1-15 | 1-16 |
|---|---|---|---|---|---|---|---|---|---|---|---|---|---|---|---|---|
| 1-1 | | 1588 | 1603 | 1248 | 67232 | 66976 | 67025 | 67152 | 75985 | 76043 | 76225 | 76190 | 36107 | 36171 | 36166 | 36105 |
| 1-2 | | | 1164 | 1468 | 67237 | 66972 | 66999 | 67066 | 76249 | 76297 | 75953 | 76354 | 35942 | 35968 | 35950 | 35994 |
| 1-14 | | | | 1511 | 67224 | 67009 | 66948 | 67100 | 76168 | 76208 | 75945 | 76259 | 35995 | 35987 | 35956 | 36004 |
| 1-15 | | | | | 67246 | 66954 | 67032 | 67124 | 76078 | 76132 | 76284 | 76264 | 36103 | 36024 | 36054 | 36029 |
| 1-3 | | | | | | 1535 | 1559 | 1599 | 36105 | 36194 | 37015 | 36170 | 76590 | 76545 | 76474 | 76470 |
| 1-4 | | | | | | | 1362 | 1335 | 35875 | 35846 | 36575 | 35980 | 76465 | 76350 | 76274 | 76319 |
| 1-5 | | | | | | | | 1601 | 35971 | 35936 | 36726 | 36034 | 76303 | 76327 | 76257 | 76288 |
| 1-6 | | | | | | | | | 35940 | 35852 | 36587 | 35929 | 76371 | 76325 | 76245 | 76286 |
| 1-7 | | | | | | | | | | 1165 | 2220 | 1236 | 67059 | 66966 | 66886 | 66896 |
| 1-8 | | | | | | | | | | | 2137 | 1023 | 67108 | 66987 | 66927 | 66896 |
| 1-9 | | | | | | | | | | | | 2180 | 66991 | 66861 | 66926 | 66898 |
| 1-10 | | | | | | | | | | | | | 67106 | 67040 | 67023 | 67018 |
| 1-11 | | | | | | | | | | | | | | 1552 | 1454 | 1506 |
| 1-12 | | | | | | | | | | | | | | | 1268 | 1232 |
| 1-15 | | | | | | | | | | | | | | | | 1189 |
| 1-16 | | | | | | | | | | | | | | | | |

### The overall number of SNPs in the 6th chromosome is 104,773.

**Table S21. Summary of pairwise SNP calling between the 5th chromosomes[#] in all 16 F1 progeny of the #4 ascus generated by crossing QM6a *spo11*Δ and CBS999.97(*MAT1-1*) *spo111*Δ**

| ChV | 4-1 | 4-2 | 4-3 | 4-4 | 4-5 | 4-6 | 4-7 | 4-8 | 4-9 | 4-10 | 4-11 | 4-12 | 4-13 | 4-14 | 4-15 | 4-16 |
|---|---|---|---|---|---|---|---|---|---|---|---|---|---|---|---|---|
| 4-1 | | 2198 | 2188 | 2131 | 71585 | 71329 | 71954 | 71388 | 49937 | 50269 | 49813 | 49831 | 75196 | 75906 | 75357 | 75328 |
| 4-2 | | | 2339 | 2362 | 71592 | 71362 | 71971 | 71409 | 50079 | 50533 | 50045 | 50097 | 75260 | 75920 | 75442 | 75440 |
| 4-3 | | | | 2343 | 71631 | 71393 | 72036 | 71413 | 50061 | 50425 | 49909 | 49937 | 75231 | 75928 | 75457 | 75316 |
| 4-4 | | | | | 71609 | 71423 | 71969 | 71475 | 50123 | 50408 | 50146 | 50100 | 75462 | 76052 | 75594 | 75490 |
| 4-5 | | | | | | 2408 | 2827 | 2539 | 75737 | 75808 | 75579 | 75726 | 50144 | 50720 | 50153 | 50213 |
| 4-6 | | | | | | | 2912 | 2514 | 75402 | 75547 | 75252 | 75287 | 49998 | 50598 | 50104 | 49962 |
| 4-7 | | | | | | | | 3233 | 76060 | 76016 | 75996 | 76091 | 50556 | 50833 | 50679 | 50616 |
| 4-8 | | | | | | | | | 75349 | 75505 | 75063 | 75086 | 49797 | 50639 | 50016 | 49944 |
| 4-9 | | | | | | | | | | 2569 | 2363 | 2412 | 71210 | 71753 | 71241 | 71223 |
| 4-10 | | | | | | | | | | | 2556 | 2725 | 71693 | 71480 | 71681 | 71608 |
| 4-11 | | | | | | | | | | | | 2023 | 71226 | 71945 | 71371 | 71351 |
| 4-12 | | | | | | | | | | | | | 71267 | 72070 | 71371 | 71383 |
| 4-13 | | | | | | | | | | | | | | 3128 | 2398 | 2369 |
| 4-14 | | | | | | | | | | | | | | | 3081 | 3161 |
| 4-15 | | | | | | | | | | | | | | | | 2579 |
| 4-16 | | | | | | | | | | | | | | | | |

### The overall number of SNPs in the 5th chromosome is 115,278.

**Table S22. Summary of pairwise SNP calling between the 6th chromosomes# in all 16 F1 progeny of the #4 ascus generated by crossing QM6a *spo11*Δ and CBS999.97(*MAT1-1*) *spo11*Δ**

| ChVI | 4-1 | 4-2 | 4-3 | 4-4 | 4-5 | 4-6 | 4-7 | 4-8 | 4-9 | 4-10 | 4-11 | 4-12 | 4-13 | 4-14 | 4-15 | 4-16 |
|---|---|---|---|---|---|---|---|---|---|---|---|---|---|---|---|---|
| 4-1 | | 1684 | 1690 | 1672 | 20247 | 20332 | 20791 | 20198 | 71741 | 71958 | 71796 | 71830 | 85355 | 85773 | 85596 | 85328 |
| 4-2 | | | 1568 | 1735 | 20270 | 20279 | 20639 | 20146 | 71664 | 71864 | 71742 | 71693 | 85243 | 85662 | 85484 | 85243 |
| 4-3 | | | | 1717 | 20296 | 20260 | 20726 | 20092 | 71694 | 71945 | 71771 | 71735 | 85259 | 85686 | 85504 | 85279 |
| 4-4 | | | | | 20224 | 20223 | 20658 | 20059 | 71702 | 71969 | 71707 | 71693 | 85252 | 85648 | 85499 | 85240 |
| 4-5 | | | | | | 1910 | 2255 | 2116 | 85471 | 85666 | 85468 | 85489 | 71535 | 71946 | 71954 | 71544 |
| 4-6 | | | | | | | 2187 | 1948 | 85294 | 85547 | 85357 | 85359 | 71446 | 71869 | 71855 | 71503 |
| 4-7 | | | | | | | | 2409 | 85695 | 85843 | 85712 | 85769 | 71819 | 72059 | 72208 | 71852 |
| 4-8 | | | | | | | | | 85232 | 85497 | 85238 | 85291 | 71423 | 71829 | 71803 | 71410 |
| 4-9 | | | | | | | | | | 2221 | 2182 | 2259 | 20357 | 20684 | 20613 | 20665 |
| 4-10 | | | | | | | | | | | 2612 | 2676 | 20645 | 20684 | 20895 | 20939 |
| 4-11 | | | | | | | | | | | | 1949 | 20540 | 20933 | 20283 | 20307 |
| 4-12 | | | | | | | | | | | | | 20630 | 21055 | 20501 | 20544 |
| 4-13 | | | | | | | | | | | | | | 2296 | 2324 | 2192 |
| 4-14 | | | | | | | | | | | | | | | 2769 | 2710 |
| 4-15 | | | | | | | | | | | | | | | | 2114 |
| 4-16 | | | | | | | | | | | | | | | | |

### The overall number of SNPs in the 6th chromosome is 105,116.

**Table S23. Distances in the 6 asci for all COs to the two neighboring AT-rich blocks.**

| Asci number | Chromosome number | CO position (bp) | Position of the 5' neighboring AT-rich block | Length of 5' neighboring AT-rich block (bp) | Distance of CO to the 5' neighboring AT-rich block# | Position of the 3' neighboring AT-rich block | Length of 3' neighboring AT-rich block (bp) | Distance of CO to the 3' neighboring AT-rich bloc# |
|---|---|---|---|---|---|---|---|---|
| 1 | 1 | 1376408 | 1361500-1362000 | 500 | 14408 | 1402000-1402500 | 500 | 25592 |
| 1 | 1 | 5942633 | 5929000-5930000 | 1000 | 12633 | 6007500-6009000 | 1500 | 64867 |
| 1 | 1 | 4850122 | 4846000-4847000 | 1000 | 3122 | 4860500-4861500 | 1000 | 10379 |
| 1 | 1 | 2846616 | 2823000-2823500 | 500 | 23116 | 2856000-2856500 | 500 | 9384 |
| 1 | 1 | 290545 | 262500-263000 | 500 | 27545 | 291000-291500 | 500 | 455 |
| 1 | 2 | 1078314 | 1066500-1067000 | 500 | 11314 | 1082000-1082500 | 500 | 3686 |
| 1 | 2 | 3067880 | 3047000-3051000 | 4000 | 16880 | 3068500-3069000 | 500 | 620 |
| 1 | 2 | 4581277 | 4576000-4576500 | 500 | 4777 | 4593500-4594000 | 500 | 12224 |
| 1 | 3 | 2230879 | 2198500-2199000 | 500 | 31879 | 2238500-2239500 | 1000 | 7621 |
| 1 | 3 | 389221 | 387000-388000 | 1000 | 1221 | 394500-395000 | 500 | 5280 |
| 1 | 3 | 1639304 | 1630000-1632500 | 2500 | 6804 | 1639500-1640000 | 500 | 197 |
| 1 | 4 | 3937471 | 3919500-3920000 | 500 | 17471 | 3965500-3966000 | 500 | 28029 |
| 1 | 4 | 823739 | 792500-793000 | 500 | 30739 | 885000-888500 | 3500 | 61262 |
| 1 | 4 | 1922011 | 1903000-1903500 | 500 | 18511 | 1934000-1935000 | 1000 | 11989 |
| 1 | 5 | 906317 | 906000-907000 | 1000 | 0 | 906000-907000 | 1000 | 0 |

| | | | | | | | | |
|---|---|---|---|---|---|---|---|---|
| 1 | 5 | 1651350 | 1590500-1591000 | 500 | 60350 | 1655000-1655500 | 500 | 3650 |
| 1 | 5 | 2640861 | 2640000-2640500 | 500 | 361 | 2652500-2653000 | 500 | 11639 |
| 1 | 5 | 3965025 | 3941500-3942000 | 500 | 23025 | 3985000-3985500 | 500 | 19976 |
| 1 | 6 | 2245856 | 2244500-2245500 | 1000 | 356 | 2251500-2252000 | 500 | 5644 |
| 1 | 6 | 3266023 | 3265000-3265500 | 500 | 523 | 3280500-3281000 | 500 | 14478 |
| 1 | 6 | 640823 | 635500-636000 | 500 | 4823 | 663000-663500 | 500 | 22178 |
| 1 | 7 | 2891266 | 2867500-2879000 | 11500 | 12266 | 2898000-2898500 | 500 | 6734 |
| 1 | 7 | 1180772 | 1125500-1126000 | 500 | 54772 | 1207000-1207500 | 500 | 26229 |
| 1 | 7 | 3411537 | 3401000-3401500 | 500 | 10037 | 3418000-3418500 | 500 | 6464 |
| 2 | 1 | 4251534 | 4242000-4242500 | 500 | 9034 | 4253000-4256000 | 3000 | 1466 |
| 2 | 1 | 442660 | 439000-439500 | 500 | 3160 | 449000-449500 | 500 | 6340 |
| 2 | 1 | 5974853 | 5929000-5930000 | 1000 | 44853 | 6007500-6009000 | 1500 | 32648 |
| 2 | 1 | 2084714 | 2083500-2084000 | 500 | 714 | 2103500-2104000 | 500 | 18786 |
| 2 | 1 | 1506686 | 1504000-1504500 | 500 | 2186 | 1524000-1524500 | 500 | 17315 |
| 2 | 2 | 4300667 | 4295000-4295500 | 500 | 5167 | 4335500-4336000 | 500 | 34834 |
| 2 | 2 | 1674609 | 1660000-1660500 | 500 | 14109 | 1685500-1686500 | 1000 | 10892 |
| 2 | 2 | 5020065 | 4988500-5008000 | 19500 | 12065 | 5029000-5029500 | 500 | 8935 |
| 2 | 2 | 3361095 | 3358000-3360000 | 2000 | 1095 | 3368000-3368500 | 500 | 6905 |
| 2 | 3 | 3159413 | 3139000-3139500 | 500 | 19913 | 3193500-3194500 | 1000 | 34088 |
| 2 | 3 | 1423219 | 1411500-1412000 | 500 | 11219 | 1452000-1452500 | 500 | 28782 |
| 2 | 3 | 514337 | 489500-498500 | 9000 | 15837 | 523000-524000 | 1000 | 8664 |
| 2 | 3 | 2307834 | 2296500-2298000 | 1500 | 9834 | 2308500-2309000 | 500 | 666 |

| | | | | | | | | |
|---|---|---|---|---|---|---|---|---|
| 2 | 4 | 2627476 | 2624500-2625500 | 1000 | 1976 | 2654500-2655500 | 1000 | 27024 |
| 2 | 4 | 575881 | 559500-560000 | 500 | 15881 | 579000-579500 | 500 | 3120 |
| 2 | 4 | 4085717 | 4072500-4073000 | 500 | 12717 | 4101500-4102000 | 500 | 15783 |
| 2 | 5 | 745214 | 743000-744000 | 1000 | 1214 | 752000-752500 | 500 | 6787 |
| 2 | 5 | 3666802 | 3663000-3663500 | 500 | 3302 | 3669500-3671500 | 2000 | 2698 |
| 2 | 5 | 1725040 | 1712500-1713000 | 500 | 12040 | 1734000-1734500 | 500 | 8960 |
| 2 | 6 | 3013187 | 2994000-2994500 | 500 | 18687 | 3035000-3036000 | 1000 | 21813 |
| 2 | 6 | 832831 | 802500-803000 | 500 | 29831 | 849500-850000 | 500 | 16670 |
| 2 | 7 | 3111216 | 3106500-3107000 | 500 | 4216 | 3153500-3154500 | 1000 | 42284 |
| 2 | 7 | 2211609 | 2194500-2195500 | 1000 | 16109 | 2222500-2223000 | 500 | 10892 |
| 2 | 7 | 1243450 | 1238000-1238500 | 500 | 4950 | 1251000-1251500 | 500 | 7551 |
| 3 | 1 | 2704428 | 2654500-2655500 | 1000 | 48928 | 2711500-2713000 | 1500 | 7073 |
| 3 | 1 | 801341 | 792000-793000 | 1000 | 8341 | 801500-802000 | 500 | 159 |
| 3 | 1 | 5831682 | 5812000-5813000 | 1000 | 18682 | 5832500-5833000 | 500 | 818 |
| 3 | 1 | 250750 | 228500-229000 | 500 | 21750 | 262500-263000 | 500 | 11751 |
| 3 | 2 | 1673658 | 1660000-1660500 | 500 | 13158 | 1685500-1686500 | 1000 | 11842 |
| 3 | 2 | 1523840 | 1521500-1522000 | 500 | 1840 | 1538000-1538500 | 500 | 14160 |
| 3 | 2 | 423288 | 420500-421000 | 500 | 2288 | 426500-427000 | 500 | 3212 |
| 3 | 2 | 3336022 | 3323000-3326500 | 3500 | 9522 | 3337500-3338000 | 500 | 1479 |
| 3 | 2 | 2538004 | 2534500-2535000 | 500 | 3004 | 2556000-2556500 | 500 | 17996 |
| 3 | 2 | 5223333 | 5208000-5208500 | 500 | 14833 | 5224000-5224500 | 500 | 668 |
| 3 | 3 | 537359 | 527000-527500 | 500 | 9859 | 539500-540000 | 500 | 2141 |

| | | | | | | | | |
|---|---|---|---|---|---|---|---|---|
| 3 | 3 | 1395321 | 1392000-1393000 | 1000 | 2321 | 1399000-1399500 | 500 | 3680 |
| 3 | 3 | 2086093 | 2069000-2069500 | 500 | 16593 | 2091000-2091500 | 500 | 4908 |
| 3 | 4 | 558014 | 546500-547000 | 500 | 11014 | 559500-560000 | 500 | 1486 |
| 3 | 4 | 2676814 | 2654500-2655500 | 1000 | 21314 | 2680500-2681000 | 500 | 3686 |
| 3 | 4 | 1902320 | 1889500-1890000 | 500 | 12320 | 1903000-1903500 | 500 | 680 |
| 3 | 4 | 3557828 | 3540000-3540500 | 500 | 17328 | 3558000-3558500 | 500 | 172 |
| 3 | 5 | 2923941 | 2923000-2923500 | 500 | 441 | 2926500-2927000 | 500 | 2559 |
| 3 | 5 | 3953162 | 3941500-3942000 | 500 | 11162 | 3985000-3985500 | 500 | 31838 |
| 3 | 5 | 775941 | 752000-752500 | 500 | 23441 | 784000-788000 | 4000 | 8059 |
| 3 | 5 | 1672593 | 1655000-1655500 | 500 | 17093 | 1690500-1699000 | 8500 | 17908 |
| 3 | 6 | 2581272 | 2580000-2580500 | 500 | 772 | 2597500-2598000 | 500 | 16228 |
| 3 | 6 | 1444345 | 1436000-1436500 | 500 | 7845 | 1460000-1460500 | 500 | 15655 |
| 3 | 6 | 3091270 | 3090000-3090500 | 500 | 770 | 3106000-3106500 | 500 | 14730 |
| 3 | 6 | 600939 | 590500-591000 | 500 | 9939 | 612500-613000 | 500 | 11562 |
| 3 | 7 | 1092932 | 1090000-1090500 | 500 | 2432 | 1121000-1124000 | 3000 | 28069 |
| 3 | 7 | 3110136 | 3106500-3107000 | 500 | 3136 | 3153500-3154500 | 1000 | 43365 |
| 4 | 1 | 2241851 | 2241000-2241500 | 500 | 351 | 2280500-2286500 | 6000 | 38649 |
| 4 | 1 | 6409791 | 6385500-6386000 | 500 | 23791 | 6416000-6416500 | 500 | 6209 |
| 4 | 1 | 4763682 | 4761500-4762000 | 500 | 1682 | 4773000-4773500 | 500 | 9319 |
| 4 | 1 | 1769754 | 1764500-1765000 | 500 | 4754 | 1775000-1775500 | 500 | 5246 |
| 4 | 2 | 2897973 | 2896500-2897000 | 500 | 973 | 2931500-2932000 | 500 | 33527 |
| 4 | 2 | 2156538 | 2156500-2157000 | 500 | 0 | 2156500-2157000 | 500 | 0 |

| | | | | | | | | |
|---|---|---|---|---|---|---|---|---|
| 4 | 2 | 1103742 | 1102000-1102500 | 500 | 1242 | 1148500-1149000 | 500 | 44758 |
| 4 | 3 | 992231 | 991500-992500 | 1000 | 0 | 991500-992500 | 1000 | 0 |
| 4 | 3 | 2827971 | 2820000-2823500 | 3500 | 4471 | 2829500-2830000 | 500 | 1529 |
| 4 | 4 | 4020058 | 4012500-4013000 | 500 | 7058 | 4065000-4065500 | 500 | 44942 |
| 4 | 4 | 1242956 | 1236500-1237000 | 500 | 5956 | 1271000-1274000 | 3000 | 28044 |
| 4 | 4 | 3095783 | 3093500-3094000 | 500 | 1783 | 3101500-3102000 | 500 | 5718 |
| 4 | 5 | 2588287 | 2588000-2588500 | 500 | 0 | 2588000-2588500 | 500 | 0 |
| 4 | 5 | 519252 | 517500-518000 | 500 | 1252 | 534000-534500 | 500 | 14748 |
| 4 | 5 | 4088777 | 4087500-4088000 | 500 | 777 | 4089000-4089500 | 500 | 223 |
| 4 | 5 | 1690152 | 1655000-1655500 | 500 | 34652 | 1690500-1699000 | 8500 | 349 |
| 4 | 6 | 3270110 | 3265000-3265500 | 500 | 4610 | 3280500-3281000 | 500 | 10390 |
| 4 | 6 | 244330 | 235000-238000 | 3000 | 6330 | 276500-277000 | 500 | 32170 |
| 4 | 7 | 1283658 | 1283000-1284000 | 1000 | 0 | 1283000-1284000 | 1000 | 0 |
| 5 | 1 | 2735525 | 2734500-2735000 | 500 | 525 | 2749000-2749500 | 500 | 13476 |
| 5 | 2 | 2540154 | 2534500-2535000 | 500 | 5154 | 2556000-2556500 | 500 | 15847 |
| 5 | 2 | 3334530 | 3323000-3326500 | 3500 | 8030 | 3337500-3338000 | 500 | 2971 |
| 5 | 3 | 4879689 | 4864000-4867500 | 3500 | 12189 | 4887500-4888000 | 500 | 7812 |
| 5 | 3 | 1482429 | 1457000-1457500 | 500 | 24929 | 1497500-1498000 | 500 | 15072 |
| 5 | 4 | 1241154 | 1236500-1237000 | 500 | 4154 | 1271000-1274000 | 3000 | 29846 |
| 5 | 4 | 3145239 | 3134500-3135000 | 500 | 10239 | 3167000-3167500 | 500 | 21762 |
| 5 | 5 | 3164025 | 3163000-3163500 | 500 | 525 | 3174000-3174500 | 500 | 9975 |
| 5 | 5 | 1803005 | 1779500-1780000 | 500 | 23005 | 1803500-1804000 | 500 | 496 |

| | | | | | | | | |
|---|---|---|---|---|---|---|---|---|
| 5 | 6 | 1761238 | 1760500-1761000 | 500 | 238 | 1769000-1769500 | 500 | 7762 |
| 5 | 7 | 2221807 | 2194500-2195500 | 1000 | 26307 | 2222500-2223000 | 500 | 693 |
| 5 | 7 | 2859082 | 2848000-2848500 | 500 | 10582 | 2867500-2879000 | 11500 | 8419 |
| 6 | 1 | 1774730 | 1764500-1765000 | 500 | 9730 | 1775000-1775500 | 500 | 270 |
| 6 | 1 | 5629303 | 5625000-5626000 | 1000 | 3303 | 5646500-5647000 | 500 | 17197 |
| 6 | 1 | 2566477 | 2553500-2554000 | 500 | 12477 | 2588000-2589000 | 1000 | 21523 |
| 6 | 1 | 6704878 | 6697500-6698000 | 500 | 6878 | 6709500-6710000 | 500 | 4622 |
| 6 | 1 | 513750 | 497000-497500 | 500 | 16250 | 525000-525500 | 500 | 11250 |
| 6 | 2 | 2609280 | 2604000-2604500 | 500 | 4780 | 2631000-2631500 | 500 | 21720 |
| 6 | 2 | 3901105 | 3900500-3901500 | 1000 | 0 | 3900500-3901500 | 1000 | 0 |
| 6 | 2 | 1870931 | 1870000-1870500 | 500 | 431 | 1940500-2103000 | 162500 | 69570 |
| 6 | 2 | 233590 | 233500-234000 | 500 | 0 | 233500-234000 | 500 | 0 |
| 6 | 3 | 1074005 | 1065500-1066000 | 500 | 8005 | 1075500-1076500 | 1000 | 1495 |
| 6 | 4 | 4144753 | 4124000-4124500 | 500 | 20253 | 4165500-4181500 | 16000 | 20747 |
| 6 | 4 | 1965258 | 1934000-1935000 | 1000 | 30258 | 1986000-1986500 | 500 | 20742 |
| 6 | 4 | 2658162 | 2654500-2655500 | 1000 | 2662 | 2680500-2681000 | 500 | 22338 |
| 6 | 4 | 1188532 | 1187000-1187500 | 500 | 1032 | 1199500-1200000 | 500 | 10969 |
| 6 | 4 | 560886 | 559500-560000 | 500 | 886 | 579000-579500 | 500 | 18115 |
| 6 | 5 | 2594043 | 2588000-2588500 | 500 | 5543 | 2598500-2599000 | 500 | 4457 |
| 6 | 5 | 1894579 | 1876500-1877000 | 500 | 17579 | 1901500-1902000 | 500 | 6921 |
| 6 | 6 | 1586651 | 1569000-1569500 | 500 | 17151 | 1595500-1596000 | 500 | 8849 |
| 6 | 6 | 643589 | 635500-636000 | 500 | 7589 | 663000-663500 | 500 | 19411 |

| | | | | | | | | |
|---|---|---|---|---|---|---|---|---|
| 6 | 7 | 1603731 | 1601000-1601500 | 500 | 2231 | 1665000-1665500 | 500 | 61270 |
| 6 | 7 | 3372988 | 3357000-3357500 | 500 | 15488 | 3388500-3389000 | 500 | 15512 |

#The COs locate within or overlap with to the AT-rich blocks (i.e., distance = 0) are marked in blue. There are 7 COs overlapping with short AT-rich blocks (≤1000 bp in length).

#### Table S24. Distances in the 6 asci for all NCOs to the two neighboring AT-rich blocks.

| Asci number | Chromosome number | CO position (bp) | Position of the 5' neighboring AT-rich block | Length of 5' neighboring AT-rich block (bp) | Distance of CO to the 5' neighboring AT-rich block# | Position of the 3' neighboring AT-rich block | Length of 3' neighboring AT-rich block (bp) | Distance of CO to the 3' neighboring AT-rich bloc# |
|---|---|---|---|---|---|---|---|---|
| 1 | 1 | 2841836 | 2823000-2823500 | 500 | 18336 | 2856000-2856500 | 500 | 14164 |
| 1 | 2 | 1379384 | 1355500-1356000 | 500 | 23384 | 1396500-1397000 | 500 | 17116 |
| 1 | 3 | 1308219 | 1281500-1282000 | 500 | 26219 | 1312500-1313000 | 500 | 4281 |
| 1 | 3 | 1581203 | 1571000-1571500 | 500 | 9703 | 1607000-1607500 | 500 | 25798 |
| 1 | 3 | 3935255 | 3929000-3929500 | 500 | 5755 | 3935500-3936000 | 500 | 246 |
| 1 | 3 | 4101515 | 4003500-4004000 | 500 | 97515 | 4108500-4109000 | 500 | 6985 |
| 1 | 5 | 783456 | 752000-752500 | 500 | 30956 | 784000-788000 | 4000 | 544 |
| 1 | 5 | 2906231 | 2895000-2895500 | 500 | 10731 | 2923000-2923500 | 500 | 16769 |
| 2 | 1 | 1370017 | 1361500-1362000 | 500 | 8017 | 1402000-1402500 | 500 | 31984 |
| 2 | 1 | 3651818 | 3645000-3645500 | 500 | 6318 | 3677000-3677500 | 500 | 25183 |
| 2 | 1 | 3712471 | 3712000-3712500 | 500 | 0 | 3712000-3712500 | 500 | 4529 |
| 2 | 2 | 1085027 | 1082000-1082500 | 500 | 2527 | 1092000-1092500 | 500 | 6973 |
| 2 | 2 | 4571770 | 4568500-4569000 | 500 | 2770 | 4576000-4576500 | 500 | 4230 |
| 2 | 2 | 5129446 | 5102000-5102500 | 500 | 26946 | 5146000-5146500 | 500 | 16554 |
| 2 | 3 | 1577617 | 1571000-1571500 | 500 | 6117 | 1607000-1607500 | 500 | 29384 |
| 2 | 3 | 2103899 | 2091000-2091500 | 500 | 12399 | 2107000-2107500 | 500 | 3102 |

| | | | | | | | | |
|---|---|---|---|---|---|---|---|---|
| 2 | 3 | 2432035 | 2431000-2431500 | 500 | 535 | 2433000-2434000 | 1000 | 965 |
| 2 | 4 | 1138997 | 1138500-1139000 | 500 | 0 | 1138500-1139000 | 500 | 21503 |
| 2 | 4 | 1725946 | 1718500-1722000 | 3500 | 3946 | 1736500-1737000 | 500 | 10554 |
| 2 | 6 | 1408314 | 1397000-1397500 | 500 | 10814 | 1422000-1422500 | 500 | 13686 |
| 2 | 7 | 2141143 | 2138500-2139000 | 500 | 2143 | 2141500-2142500 | 1000 | 357 |
| 3 | 1 | 1958346 | 1912500-1913000 | 500 | 45346 | 1973000-1973500 | 500 | 14655 |
| 3 | 1 | 254869 | 228500-229000 | 500 | 25869 | 262500-263000 | 500 | 7631 |
| 3 | 2 | 276387 | 258500-259000 | 500 | 17387 | 303000-303500 | 500 | 26613 |
| 3 | 2 | 1081326 | 1066500-1067000 | 500 | 14326 | 1082000-1082500 | 500 | 674 |
| 3 | 2 | 1516659 | 1477000-1477500 | 500 | 39159 | 1521500-1522000 | 500 | 4841 |
| 3 | 2 | 4581999 | 4576000-4576500 | 500 | 5499 | 4593500-4594000 | 500 | 11502 |
| 3 | 3 | 2084400 | 2069000-2069500 | 500 | 14900 | 2091000-2091500 | 500 | 6600 |
| 3 | 3 | 2086858 | 2069000-2069500 | 500 | 17358 | 2091000-2091500 | 500 | 4142 |
| 3 | 3 | 4831375 | 4811000-4814000 | 3000 | 17375 | 4864000-4867500 | 3500 | 32625 |
| 3 | 5 | 164706 | 151000-151500 | 500 | 13206 | 175500-176000 | 500 | 10795 |
| 3 | 6 | 1263128 | 1260500-1261000 | 500 | 2128 | 1265500-1266000 | 500 | 2372 |
| 3 | 6 | 2582114 | 2580000-2580500 | 500 | 1614 | 2597500-2598000 | 500 | 15386 |
| 3 | 7 | 532598 | 529000-530000 | 1000 | 2598 | 535500-536500 | 1000 | 2902 |
| 3 | 7 | 714647 | 691500-692000 | 500 | 22647 | 720000-720500 | 500 | 5353 |
| 3 | 7 | 2433134 | 2426500-2427000 | 500 | 6134 | 2450500-2451000 | 500 | 17366 |
| 4 | 1 | 2854242.5 | 2823000-2823500 | 500 | 30742.5 | 2856000-2856500 | 500 | 1757.5 |
| 4 | 1 | 2960509.75 | 2946500-2947000 | 500 | 13509.75 | 2964000-2964500 | 500 | 3490.25 |

| | | | | | | | | |
|---|---|---|---|---|---|---|---|---|
| 4 | 3 | 984089.75 | 982500-983000 | 500 | 1089.75 | 991500-992500 | 1000 | 7410.25 |
| 4 | 3 | 2986266.5 | 2985500-2986000 | 500 | 266.5 | 2998500-2999000 | 500 | 12233.5 |
| 4 | 4 | 2770557 | 2765500-2766000 | 500 | 4557 | 2777500-2778500 | 1000 | 6943 |
| 4 | 5 | 3155775.25 | 3153500-3154000 | 500 | 1775.25 | 3163000-3163500 | 500 | 7224.75 |
| 4 | 7 | 2773899.75 | 2762000-2763000 | 1000 | 10899.75 | 2774000-2774500 | 500 | 100.25 |
| 5 | 3 | 1482190 | 1457000-1457500 | 500 | 24690 | 1497500-1498000 | 500 | 15310 |
| 6 | 1 | 2041920.5 | 2041500-2042000 | 500 | 0 | 2041500-2042000 | 500 | 14079.5 |
| 6 | 1 | 6124817.25 | 6114000-6114500 | 500 | 10317.25 | 6135000-6136000 | 1000 | 10182.75 |
| 6 | 1 | 1472822.75 | 1471000-1471500 | 500 | 1322.75 | 1485000-1486000 | 1000 | 12177.25 |
| 6 | 2 | 3912412.75 | 3900500-3901500 | 1000 | 10912.75 | 3929500-3930000 | 500 | 17087.25 |
| 6 | 2 | 4548731 | 4537000-4537500 | 500 | 11231 | 4568500-4569000 | 500 | 19769 |
| 6 | 2 | 5637408.75 | 5627500-5628500 | 1000 | 8908.75 | 5644000-5644500 | 500 | 6591.25 |
| 6 | 3 | 188813.25 | 172000-172500 | 500 | 16313.25 | 194500-195000 | 500 | 5686.75 |
| 6 | 3 | 1577677.75 | 1571000-1571500 | 500 | 6177.75 | 1607000-1607500 | 500 | 29322.25 |
| 6 | 3 | 2267131.25 | 2238500-2239500 | 1000 | 27631.25 | 2275500-2276500 | 1000 | 8368.75 |
| 6 | 3 | 5159263.5 | 5157000-5157500 | 500 | 1763.5 | 5213000-5213500 | 500 | 53736.5 |
| 6 | 4 | 1457597.25 | 1442500-1443000 | 500 | 14597.25 | 1460000-1460500 | 500 | 2402.75 |
| 6 | 7 | 1664463.75 | 1601000-1601500 | 500 | 62963.75 | 1665000-1665500 | 500 | 536.25 |
| 6 | 7 | 1797456.5 | 1741000-1914000 | 173000 | 0 | 1741000-1914000 | 173000 | 0 |

#The NCOs locate within or overlap with to the AT-rich blocks (i.e., distance = 0) are marked in blue. There are 5 NCOs overlapping with short AT-rich blocks (≤1000 bp in length).

**Table S25. Coefficient of coincidence and interference for all neighboring interhomolog products in the three asci (#1-#3) generated by crossing QM6a and CBS999.97(*MAT1-1*)**

| Chromosome number | Asci number | 5' position of interhomolog products | 3' position of interhomolog products | Distance (bp) between two neighboring interhomolog | Actual double interhomolog products (cM) | coefficient of coincidence (c.o.c) | Interference (I = 1- c.o.c) |
|---|---|---|---|---|---|---|---|
| 1 | 1 | 290545 | 1376408 | 1085863 | 0.39 | 0.78 | 0.22 |
| 1 | 1 | 1376408 | 2841836 | 1465428 | 0.52 | 1.05 | -0.05 |
| 1 | 1 | 2841836 | 2846616 | 4780 | 0.00 | 0.00 | 1.00 |
| 1 | 1 | 2846616 | 4850122 | 2003506 | 0.72 | 1.43 | -0.43 |
| 1 | 1 | 4850122 | 5942633 | 1092512 | 0.39 | 0.78 | 0.22 |
| 1 | 2 | 1078314 | 1379384 | 301070 | 0.11 | 0.22 | 0.78 |
| 1 | 2 | 1379384 | 3067880 | 1688496 | 0.60 | 1.21 | -0.21 |
| 1 | 2 | 3067880 | 4581277 | 1513397 | 0.54 | 1.08 | -0.08 |
| 1 | 3 | 389221 | 1308219 | 918999 | 0.33 | 0.66 | 0.34 |
| 1 | 3 | 1308219 | 1581203 | 272984 | 0.10 | 0.20 | 0.80 |
| 1 | 3 | 1581203 | 1639304 | 58101 | 0.02 | 0.04 | 0.96 |
| 1 | 3 | 1639304 | 2230879 | 591576 | 0.21 | 0.42 | 0.58 |
| 1 | 3 | 2230879 | 3935255 | 1704376 | 0.61 | 1.22 | -0.22 |
| 1 | 3 | 3935255 | 4101515 | 166261 | 0.06 | 0.12 | 0.88 |
| 1 | 4 | 823739 | 1922011 | 1098272 | 0.39 | 0.79 | 0.21 |

| | | | | | | | |
|---|---|---|---|---|---|---|---|
| 1 | 4 | 1922011 | 3937471 | 2015460 | 0.72 | 1.44 | -0.44 |
| 1 | 5 | 783456 | 906317 | 122861 | 0.04 | 0.09 | 0.91 |
| 1 | 5 | 906317 | 1651350 | 745033 | 0.27 | 0.53 | 0.47 |
| 1 | 5 | 1651350 | 2640861 | 989511 | 0.35 | 0.71 | 0.29 |
| 1 | 5 | 2640861 | 2906231 | 265370 | 0.09 | 0.19 | 0.81 |
| 1 | 5 | 2906231 | 3965025 | 1058794 | 0.38 | 0.76 | 0.24 |
| 1 | 6 | 640823 | 2245856 | 1605034 | 0.57 | 1.15 | -0.15 |
| 1 | 6 | 2245856 | 3266023 | 1020167 | 0.37 | 0.73 | 0.27 |
| 1 | 7 | 1180772 | 2891266 | 1710495 | 0.61 | 1.22 | -0.22 |
| 1 | 7 | 2891266 | 3411537 | 520271 | 0.19 | 0.37 | 0.63 |
| 2 | 1 | 442660 | 1370017 | 927356 | 0.33 | 0.66 | 0.34 |
| 2 | 1 | 1370017 | 1506686 | 136669 | 0.05 | 0.10 | 0.90 |
| 2 | 1 | 1506686 | 2084714 | 578029 | 0.21 | 0.41 | 0.59 |
| 2 | 1 | 2084714 | 3651818 | 1567104 | 0.56 | 1.12 | -0.12 |
| 2 | 1 | 3651818 | 3712471 | 60653 | 0.02 | 0.04 | 0.96 |
| 2 | 1 | 3712471 | 4251534 | 539063 | 0.19 | 0.39 | 0.61 |
| 2 | 1 | 4251534 | 5974853 | 1723319 | 0.62 | 1.23 | -0.23 |
| 2 | 2 | 1085027 | 1674609 | 589581 | 0.21 | 0.42 | 0.58 |
| 2 | 2 | 1674609 | 3361095 | 1686487 | 0.60 | 1.21 | -0.21 |
| 2 | 2 | 3361095 | 4300667 | 939571 | 0.34 | 0.67 | 0.33 |
| 2 | 2 | 4300667 | 4571770 | 271104 | 0.10 | 0.19 | 0.81 |
| 2 | 2 | 4571770 | 5020065 | 448295 | 0.16 | 0.32 | 0.68 |

| | | | | | | | |
|---|---|---|---|---|---|---|---|
| 2 | 2 | 5020065 | 5129446 | 109381 | 0.04 | 0.08 | 0.92 |
| 2 | 3 | 514337 | 1423219 | 908882 | 0.33 | 0.65 | 0.35 |
| 2 | 3 | 1423219 | 1577617 | 154398 | 0.06 | 0.11 | 0.89 |
| 2 | 3 | 1577617 | 2103899 | 526282 | 0.19 | 0.38 | 0.62 |
| 2 | 3 | 2103899 | 2307834 | 203935 | 0.07 | 0.15 | 0.85 |
| 2 | 3 | 2307834 | 2432035 | 124201 | 0.04 | 0.09 | 0.91 |
| 2 | 3 | 2432035 | 3159413 | 727378 | 0.26 | 0.52 | 0.48 |
| 2 | 4 | 575881 | 1138997 | 563116 | 0.20 | 0.40 | 0.60 |
| 2 | 4 | 1138997 | 1725946 | 586949 | 0.21 | 0.42 | 0.58 |
| 2 | 4 | 1725946 | 2627476 | 901531 | 0.32 | 0.65 | 0.35 |
| 2 | 4 | 2627476 | 4085717 | 1458241 | 0.52 | 1.04 | -0.04 |
| 2 | 5 | 745214 | 1725040 | 979827 | 0.35 | 0.70 | 0.30 |
| 2 | 5 | 1725040 | 3666802 | 1941762 | 0.70 | 1.39 | -0.39 |
| 2 | 6 | 832831 | 1408314 | 575484 | 0.21 | 0.41 | 0.59 |
| 2 | 6 | 1408314 | 3013187 | 1604873 | 0.57 | 1.15 | -0.15 |
| 2 | 7 | 1243450 | 2141143 | 897694 | 0.32 | 0.64 | 0.36 |
| 2 | 7 | 2141143 | 2211609 | 70466 | 0.03 | 0.05 | 0.95 |
| 2 | 7 | 2211609 | 3111216 | 899608 | 0.32 | 0.64 | 0.36 |
| 3 | 1 | 250750 | 254869 | 4119 | 0.00 | 0.00 | 1.00 |
| 3 | 1 | 254869 | 801341 | 546472 | 0.20 | 0.39 | 0.61 |
| 3 | 1 | 801341 | 1958346 | 1157005 | 0.41 | 0.83 | 0.17 |
| 3 | 1 | 1958346 | 2704428 | 746082 | 0.27 | 0.53 | 0.47 |

| | | | | | | | |
|---|---|---|---|---|---|---|---|
| 3 | 1 | 2704428 | 5831682 | 3127255 | 1.12 | 2.24 | -1.24 |
| 3 | 2 | 276387 | 423288 | 146902 | 0.05 | 0.11 | 0.89 |
| 3 | 2 | 423288 | 1081326 | 658038 | 0.24 | 0.47 | 0.53 |
| 3 | 2 | 1081326 | 1516659 | 435333 | 0.16 | 0.31 | 0.69 |
| 3 | 2 | 1516659 | 1523840 | 7181 | 0.00 | 0.01 | 0.99 |
| 3 | 2 | 1523840 | 1673658 | 149818 | 0.05 | 0.11 | 0.89 |
| 3 | 2 | 1673658 | 2538004 | 864346 | 0.31 | 0.62 | 0.38 |
| 3 | 2 | 2538004 | 3336022 | 798018 | 0.29 | 0.57 | 0.43 |
| 3 | 2 | 3336022 | 4581999 | 1245977 | 0.45 | 0.89 | 0.11 |
| 3 | 2 | 4581999 | 5223333 | 641334 | 0.23 | 0.46 | 0.54 |
| 3 | 3 | 537359 | 1395321 | 857962 | 0.31 | 0.61 | 0.39 |
| 3 | 3 | 1395321 | 2084400 | 689080 | 0.25 | 0.49 | 0.51 |
| 3 | 3 | 2084400 | 2086093 | 1692 | 0.00 | 0.00 | 1.00 |
| 3 | 3 | 2086093 | 2086858 | 765 | 0.00 | 0.00 | 1.00 |
| 3 | 3 | 2086858 | 4831375 | 2744518 | 0.98 | 1.96 | -0.96 |
| 3 | 4 | 558014 | 1902320 | 1344306 | 0.48 | 0.96 | 0.04 |
| 3 | 4 | 1902320 | 2676814 | 774494 | 0.28 | 0.55 | 0.45 |
| 3 | 4 | 2676814 | 3557828 | 881014 | 0.32 | 0.63 | 0.37 |
| 3 | 5 | 164706 | 775941 | 611235 | 0.22 | 0.44 | 0.56 |
| 3 | 5 | 775941 | 1672593 | 896652 | 0.32 | 0.64 | 0.36 |
| 3 | 5 | 1672593 | 2923941 | 1251349 | 0.45 | 0.90 | 0.10 |
| 3 | 5 | 2923941 | 3953162 | 1029221 | 0.37 | 0.74 | 0.26 |

| | | | | | | | |
|---|---|---|---|---|---|---|---|
| 3 | 6 | 600939 | 1263128 | 662190 | 0.24 | 0.47 | 0.53 |
| 3 | 6 | 1263128 | 1444345 | 181217 | 0.06 | 0.13 | 0.87 |
| 3 | 6 | 1444345 | 2581272 | 1136927 | 0.41 | 0.81 | 0.19 |
| 3 | 6 | 2581272 | 2582114 | 842 | 0.00 | 0.00 | 1.00 |
| 3 | 6 | 2582114 | 3091270 | 509156 | 0.18 | 0.36 | 0.64 |
| 3 | 7 | 532598 | 714647 | 182049 | 0.07 | 0.13 | 0.87 |
| 3 | 7 | 714647 | 1092932 | 378285 | 0.14 | 0.27 | 0.73 |
| 3 | 7 | 1092932 | 2433134 | 1340203 | 0.48 | 0.96 | 0.04 |
| 3 | 7 | 2433134 | 3110136 | 677002 | 0.24 | 0.48 | 0.52 |

**Table S26. Coefficient of coincidence and interference for all neighboring interhomolog products (COs and NCOs) in the three asci (#4-#6) generated by crossing the first pair of QM6a *spo11*Δ and CBS999.97(*MAT1-1*) *spo11*Δ mutants**

| Chromosome number | Asci number | 5' position of interhomolog products | 3' position of interhomolog products | Distance (bp) between two neighboring interhomolog products | Actual double interhomolog products (cM) | coefficient of coincidence (c.o.c) | Interference (I = 1- c.o.c) |
|---|---|---|---|---|---|---|---|
| 1 | 1 | 1769754 | 2241851 | 472097 | 0.12 | 0.25 | 0.75 |
| 1 | 1 | 2241851 | 2854243 | 612391 | 0.16 | 0.32 | 0.68 |
| 1 | 1 | 2854243 | 2960510 | 106267 | 0.03 | 0.06 | 0.94 |
| 1 | 1 | 2960510 | 4763682 | 1803172 | 0.47 | 0.95 | 0.05 |
| 1 | 1 | 4763682 | 6409791 | 1646110 | 0.43 | 0.86 | 0.14 |
| 1 | 2 | 1103742 | 2156538 | 1052796 | 0.28 | 0.55 | 0.45 |
| 1 | 2 | 2156538 | 2897973 | 741436 | 0.19 | 0.39 | 0.61 |
| 1 | 3 | 984090 | 992231 | 8141 | 0.00 | 0.00 | 1.00 |
| 1 | 3 | 992231 | 2827971 | 1835740 | 0.48 | 0.96 | 0.04 |
| 1 | 3 | 2827971 | 2986267 | 158296 | 0.04 | 0.08 | 0.92 |

| | | | | | | | |
|---|---|---|---|---|---|---|---|
| 1 | 4 | 1242956 | 2770557 | 1527601 | 0.40 | 0.80 | 0.20 |
| 1 | 4 | 2770557 | 3095783 | 325226 | 0.09 | 0.17 | 0.83 |
| 1 | 4 | 3095783 | 4020058 | 924276 | 0.24 | 0.49 | 0.51 |
| 1 | 5 | 519252 | 1690152 | 1170900 | 0.31 | 0.61 | 0.39 |
| 1 | 5 | 1690152 | 2588287 | 898136 | 0.24 | 0.47 | 0.53 |
| 1 | 5 | 2588287 | 3155775 | 567488 | 0.15 | 0.30 | 0.70 |
| 1 | 5 | 3155775 | 4088777 | 933002 | 0.24 | 0.49 | 0.51 |
| 1 | 6 | 244330 | 3270110 | 3025780 | 0.79 | 1.59 | -0.59 |
| 1 | 7 | 1283658 | 2773900 | 1490242 | 0.39 | 0.78 | 0.22 |
| 2 | 2 | 2540154 | 3334530 | 794376 | 0.21 | 0.42 | 0.58 |
| 2 | 3 | 1482190 | 1482429 | 239 | 0.00 | 0.00 | 1.00 |
| 2 | 3 | 1482429 | 4879689 | 3397260 | 0.89 | 1.78 | -0.78 |
| 2 | 4 | 1241154 | 3145239 | 1904085 | 0.50 | 1.00 | 0.00 |
| 2 | 5 | 1803005 | 3164025 | 1361020 | 0.36 | 0.71 | 0.29 |
| 2 | 7 | 2221807 | 2859082 | 637275 | 0.17 | 0.33 | 0.67 |
| 3 | 1 | 513750 | 1472823 | 959073 | 0.25 | 0.50 | 0.50 |
| 3 | 1 | 1472823 | 1774730 | 301907 | 0.08 | 0.16 | 0.84 |
| 3 | 1 | 1774730 | 2041921 | 267191 | 0.07 | 0.14 | 0.86 |
| 3 | 1 | 2041921 | 2566477 | 524557 | 0.14 | 0.28 | 0.72 |
| 3 | 1 | 2566477 | 5629303 | 3062826 | 0.80 | 1.61 | -0.61 |
| 3 | 1 | 5629303 | 6124817 | 495514 | 0.13 | 0.26 | 0.74 |
| 3 | 1 | 6124817 | 6704878 | 580061 | 0.15 | 0.30 | 0.70 |

| | | | | | | | |
|---|---|---|---|---|---|---|---|
| 3 | 2 | 233590 | 1870931 | 1637341 | 0.43 | 0.86 | 0.14 |
| 3 | 2 | 1870931 | 2609280 | 738349 | 0.19 | 0.39 | 0.61 |
| 3 | 2 | 2609280 | 3901105 | 1291825 | 0.34 | 0.68 | 0.32 |
| 3 | 2 | 3901105 | 3912413 | 11308 | 0.00 | 0.01 | 0.99 |
| 3 | 2 | 3912413 | 4548731 | 636318 | 0.17 | 0.33 | 0.67 |
| 3 | 2 | 4548731 | 5637409 | 1088678 | 0.29 | 0.57 | 0.43 |
| 3 | 3 | 188813 | 1074005 | 885192 | 0.23 | 0.46 | 0.54 |
| 3 | 3 | 1074005 | 1577678 | 503673 | 0.13 | 0.26 | 0.74 |
| 3 | 3 | 1577678 | 2267131 | 689454 | 0.18 | 0.36 | 0.64 |
| 3 | 3 | 2267131 | 5159264 | 2892132 | 0.76 | 1.52 | -0.52 |
| 3 | 4 | 560886 | 1188532 | 627646 | 0.16 | 0.33 | 0.67 |
| 3 | 4 | 1188532 | 1457597 | 269066 | 0.07 | 0.14 | 0.86 |
| 3 | 4 | 1457597 | 1965258 | 507661 | 0.13 | 0.27 | 0.73 |
| 3 | 4 | 1965258 | 2658162 | 692904 | 0.18 | 0.36 | 0.64 |
| 3 | 4 | 2658162 | 4144753 | 1486591 | 0.39 | 0.78 | 0.22 |
| 3 | 5 | 1894579 | 2594043 | 699464 | 0.18 | 0.37 | 0.63 |
| 3 | 6 | 643589 | 1586651 | 943062 | 0.25 | 0.50 | 0.50 |
| 3 | 7 | 1603731 | 1664464 | 60733 | 0.02 | 0.03 | 0.97 |
| 3 | 7 | 1664464 | 1797457 | 132993 | 0.03 | 0.07 | 0.93 |
| 3 | 7 | 1797457 | 3372988 | 1575531 | 0.41 | 0.83 | 0.17 |

#### Table S27. Locations of interhomolog recombination products in the six asci (#1-#6)

| Ascus number | Chromosome | Recombination product | start | stop | Median position of recombination product | Located at intragenic or intergenic region | Gene start | Gene stop | distance to start codon | Gene start | Gene stop | distance to stop codon | near to 5' or 3' region |
|---|---|---|---|---|---|---|---|---|---|---|---|---|---|
| 1 | 1 | CO | 1376220 | 1376630 | 1376425 | Intragenic | 1377443 | 1372411 | 1018 | 1372368 | 1375382 | 1043 | - |
| 1 | 1 | CO | 5942621 | 5942621 | 5942621 | Intragenic | 5942255 | 5945364 | 366 | 5945213 | 5944128 | 1507 | 5' |
| 1 | 1 | CO | 4850040 | 4850203 | 4850122 | Intergenic | 4849259 | 4847117 | 863 | 4854470 | 4850258 | 137 | 3' |
| 1 | 1 | CO | 2846520 | 2846712 | 2846616 | Intragenic | 2846989 | 2845699 | 373 | 2841852 | 2846736 | 120 | 3' |
| 1 | 1 | CO | 290660 | 290417 | 290539 | Intragenic | 291274 | 289262 | 736 | 292888 | 291404 | 866 | 5' |
| 1 | 2 | CO | 1078313 | 1078229 | 1078271 | Intragenic | 1077130 | 1079666 | 1141 | 1080906 | 1078315 | 44 | 3' |
| 1 | 2 | CO | 3067677 | 3068061 | 3067869 | Intragenic | 3068400 | 3066163 | 531 | 3071357 | 3068514 | 645 | 5' |
| 1 | 2 | CO | 4581140 | 4581413 | 4581277 | Intragenic | 4581424 | 4577735 | 148 | 4580524 | 4581101 | 176 | 5' |
| 1 | 3 | CO | 2230660 | 2231098 | 2230879 | Intragenic | 2226325 | 2235510 | 4554 | 2226325 | 2235510 | 4631 | - |
| 1 | 3 | CO | 389129 | 389285 | 389207 | Intragenic | 387705 | 390723 | 1502 | 391304 | 388765 | 442 | 3' |
| 1 | 3 | CO | 1639199 | 1639497 | 1639348 | Intragenic | 1638733 | 1640301 | 615 | 1641128 | 1639096 | 252 | 3' |
| 1 | 4 | CO | 3937528 | 3937528 | 3937528 | Intragenic | 3937927 | 3935826 | 399 | 3939913 | 3938737 | 1209 | 5' |
| 1 | 4 | CO | 823701 | 823776 | 823739 | Intragenic | 819743 | 824900 | 3996 | 819743 | 824900 | 1162 | - |
| 1 | 4 | CO | 1921803 | 1922223 | 1922013 | Intragenic | 1923571 | 1927984 | 1558 | 1917870 | 1922829 | 816 | 3' |
| 1 | 5 | CO | 906347 | 906347 | 906347 | Intergenic | 911884 | 916092 | 5537 | 911985 | 908090 | 1743 | - |
| 1 | 5 | CO | 1651328 | 1651328 | 1651328 | Intragenic | 1653968 | 1648062 | 2640 | 1647992 | 1653968 | 2640 | - |

| | | | | | | | | | | | | | |
|---|---|---|---|---|---|---|---|---|---|---|---|---|---|
| 1 | 5 | CO | 2640737 | 2641088 | 2640913 | Intragenic | 2641744 | 2639555 | 832 | 2636189 | 2641198 | 286 | 3' |
| 1 | 5 | CO | 3964850 | 3965199 | 3965025 | Intragenic | 3965705 | 3961990 | 681 | 3961960 | 3965071 | 47 | 3' |
| 1 | 6 | CO | 2245618 | 2246094 | 2245856 | Intragenic | 2247952 | 2241089 | 2096 | 2241809 | 2244343 | 1513 | - |
| 1 | 6 | CO | 3266018 | 3266018 | 3266018 | Intragenic | 3268432 | 3265865 | 2414 | 3268432 | 3265865 | 153 | 3' |
| 1 | 6 | CO | 640864 | 640693 | 640779 | Intergenic | 644583 | 646889 | 3805 | 636675 | 638981 | 1798 | - |
| 1 | 7 | CO | 2891551 | 2890971 | 2891261 | Intragenic | 2894254 | 2887053 | 2993 | 2884587 | 2890494 | 767 | 3' |
| 1 | 7 | CO | 1180585 | 1180958 | 1180772 | Intragenic | 1180277 | 1184063 | 495 | 1180277 | 1184063 | 3292 | 5' |
| 1 | 7 | CO | 3411632 | 3411435 | 3411534 | Intragenic | 3409336 | 3413548 | 2198 | 3409336 | 3413548 | 2015 | - |
| 2 | 1 | CO | 4251501 | 4251404 | 4251453 | Intergenic | 4245080 | 4241893 | 6373 | 4239321 | 4243718 | 7735 | - |
| 2 | 1 | CO | 442608 | 442580 | 442594 | Intragenic | 443210 | 438353 | 616 | 444456 | 443545 | 951 | 5' |
| 2 | 1 | CO | 5974725 | 5974980 | 5974853 | Intragenic | 5976068 | 5974382 | 1216 | 5976068 | 5974382 | 471 | 3' |
| 2 | 1 | CO | 2084691 | 2084737 | 2084714 | Intergenic | 2084170 | 2084460 | 544 | 2084170 | 2084460 | 254 | 3' |
| 2 | 1 | CO | 1506745 | 1506720 | 1506733 | Intragenic | 1504562 | 1506899 | 2171 | 1504562 | 1506899 | 167 | 3' |
| 2 | 2 | CO | 4300364 | 4301025 | 4300695 | Intragenic | 4300329 | 4302172 | 366 | 4300329 | 4302172 | 1478 | 5' |
| 2 | 2 | CO | 1674268 | 1674949 | 1674609 | Intragenic | 1675682 | 1674523 | 1074 | 1675682 | 1674523 | 86 | 3' |
| 2 | 2 | CO | 5019921 | 5020184 | 5020053 | Intergenic | 5016932 | 5012493 | 3121 | 5026222 | 5023499 | 3447 | - |
| 2 | 2 | CO | 3361146 | 3361041 | 3361094 | Intragenic | 3362217 | 3362876 | 1124 | 3356057 | 3362151 | 1058 | - |
| 2 | 3 | CO | 3159300 | 3159525 | 3159413 | Intragenic | 3159873 | 3155865 | 461 | 3155805 | 3159496 | 84 | 3' |
| 2 | 3 | CO | 1423196 | 1423241 | 1423219 | Intragenic | 1423955 | 1426766 | 737 | 1418747 | 1423673 | 455 | 3' |
| 2 | 3 | CO | 514359 | 514334 | 514347 | Intragenic | 514460 | 513466 | 114 | 512519 | 513918 | 429 | 5' |
| 2 | 3 | CO | 2307631 | 2308078 | 2307855 | Intragenic | 2309490 | 2311534 | 1636 | 2298881 | 2308429 | 575 | 3' |
| 2 | 4 | CO | 2627636 | 2627293 | 2627465 | Intergenic | 2630531 | 2629760 | 3067 | 2630531 | 2629760 | 2296 | - |

| | | | | | | | | | | | | | |
|---|---|---|---|---|---|---|---|---|---|---|---|---|---|
| 2 | 4 | CO | 575839 | 575922 | 575881 | Intergenic | 571977 | 571615 | 3904 | 571977 | 571615 | 4266 | - |
| 2 | 4 | CO | 4085543 | 4085902 | 4085723 | Intragenic | 4084287 | 4087552 | 1436 | 4084287 | 4087552 | 1830 | - |
| 2 | 5 | CO | 745152 | 745275 | 745214 | Intragenic | 744612 | 746329 | 602 | 744612 | 746329 | 1116 | 5' |
| 2 | 5 | CO | 3666742 | 3666825 | 3666784 | Intergenic | 3668372 | 3667791 | 1589 | 3668372 | 3667791 | 1008 | - |
| 2 | 5 | CO | 1724856 | 1725224 | 1725040 | Intragenic | 1724511 | 1727794 | 529 | 1727684 | 1724511 | 529 | 5' |
| 2 | 6 | CO | 3012895 | 3013564 | 3013230 | Intragenic | 3012385 | 3016039 | 845 | 3015928 | 3015156 | 1927 | 5' |
| 2 | 6 | CO | 832423 | 833157 | 832790 | Intragenic | 833332 | 831220 | 542 | 830103 | 832578 | 212 | 3' |
| 2 | 7 | CO | 3110921 | 3111536 | 3111229 | Intergenic | 3111312 | 3113756 | 84 | 3110700 | 3111055 | 174 | 5' |
| 2 | 7 | CO | 2211545 | 2211672 | 2211609 | Intragenic | 2213400 | 2208426 | 1792 | 2208578 | 2212707 | 1099 | - |
| 2 | 7 | CO | 1243241 | 1243724 | 1243483 | Intergenic | 1244372 | 1246462 | 890 | 1244372 | 1246462 | 2980 | 5' |
| 3 | 1 | CO | 2704501 | 2704336 | 2704419 | Intragenic | 2702469 | 2704433 | 1950 | 2702469 | 2704433 | 15 | 3' |
| 3 | 1 | CO | 801334 | 801348 | 801341 | Intragenic | 799436 | 803103 | 1905 | 799436 | 803103 | 1762 | - |
| 3 | 1 | CO | 5831664 | 5831622 | 5831643 | Intragenic | 5830266 | 5832204 | 1377 | 5830266 | 5832204 | 561 | 3' |
| 3 | 1 | CO | 250965 | 250612 | 250789 | Intragenic | 249805 | 254132 | 984 | 254812 | 253268 | 2480 | 5' |
| 3 | 2 | CO | 1673488 | 1673857 | 1673673 | Intergenic | 1671664 | 1669794 | 2009 | 1675682 | 1674523 | 851 | 3' |
| 3 | 2 | CO | 1518542 | 1529171 | 1523857 | Intragenic | 1524596 | 1523333 | 740 | 1524596 | 1523333 | 524 | 3' |
| 3 | 2 | CO | 423367 | 423160 | 423264 | Intergenic | 422803 | 426624 | 461 | 426530 | 425256 | 1993 | 5' |
| 3 | 2 | CO | 3335826 | 3336217 | 3336022 | Intragenic | 3337401 | 3335711 | 1380 | 3337401 | 3335711 | 311 | 3' |
| 3 | 2 | CO | 2537835 | 2538219 | 2538027 | Intragenic | 2536891 | 2539464 | 1136 | 2536891 | 2539464 | 1437 | - |
| 3 | 2 | CO | 5223023 | 5223642 | 5223333 | Intragenic | 5223548 | 5223255 | 216 | 5223548 | 5223255 | 78 | 3' |
| 3 | 3 | CO | 537260 | 537458 | 537359 | Intragenic | 536233 | 538661 | 1126 | 536233 | 538661 | 1302 | - |
| 3 | 3 | CO | 1395209 | 1395432 | 1395321 | Intergenic | 1394936 | 1392987 | 385 | 1394936 | 1392987 | 2334 | 5' |

| | | | | | | | | | | | | | |
|---|---|---|---|---|---|---|---|---|---|---|---|---|---|
| 3 | 3 | CO | 2085628 | 2086547 | 2086088 | Intergenic | 2088183 | 2091283 | 2096 | 2082977 | 2083939 | 2149 | - |
| 3 | 4 | CO | 557846 | 558182 | 558014 | Intragenic | 558715 | 556426 | 701 | 558715 | 556426 | 1588 | 5' |
| 3 | 4 | CO | 2676737 | 2676891 | 2676814 | Intergenic | 2678402 | 2679638 | 1588 | 2685503 | 2677437 | 623 | 3' |
| 3 | 4 | CO | 1901961 | 1902552 | 1902257 | Intragenic | 1901016 | 1902548 | 1241 | 1901016 | 1902548 | 292 | 3' |
| 3 | 4 | CO | 3557186 | 3558505 | 3557846 | Intragenic | 3556302 | 3558266 | 1544 | 3556302 | 3558266 | 421 | 3' |
| 3 | 5 | CO | 2923997 | 2923934 | 2923966 | Intragenic | 2925852 | 2921609 | 1887 | 2921334 | 2922584 | 1382 | - |
| 3 | 5 | CO | 3952992 | 3953332 | 3953162 | Intergenic | 3951087 | 3944811 | 2075 | 3957018 | 3954864 | 1702 | - |
| 3 | 5 | CO | 775763 | 776148 | 775956 | Intergenic | 781684 | 781938 | 5729 | 781684 | 781938 | 5983 | - |
| 3 | 5 | CO | 1672372 | 1672807 | 1672590 | Intergenic | 1672910 | 1677163 | 321 | 1672910 | 1677163 | 4574 | 5' |
| 3 | 6 | CO | 2581227 | 2581227 | 2581227 | Intergenic | 2581187 | 2580555 | 40 | 2581187 | 2580555 | 672 | 5' |
| 3 | 6 | CO | 1444280 | 1444280 | 1444280 | Intragenic | 1444411 | 1439333 | 131 | 1444762 | 1447446 | 3166 | 5' |
| 3 | 6 | CO | 3090941 | 3091546 | 3091244 | Intergenic | 3088109 | 3089725 | 3135 | 3088109 | 3089725 | 1519 | - |
| 3 | 6 | CO | 600933 | 600918 | 600926 | Intragenic | 596362 | 604446 | 4564 | 596362 | 604446 | 3521 | - |
| 3 | 7 | CO | 1092699 | 1093155 | 1092927 | Intergenic | 1091455 | 1090004 | 1472 | 1094841 | 1093708 | 781 | 3' |
| 3 | 7 | CO | 3109955 | 3110271 | 3110113 | Intragenic | 3110700 | 3111055 | 587 | 3107533 | 3110564 | 451 | 3' |
| 1 | 1 | NCO | 2841732 | 2842041 | 2841887 | Intragenic | 2841852 | 2846736 | 35 | 2846989 | 2845699 | 3813 | 5' |
| 1 | 2 | NCO | 1379241 | 1379528 | 1379385 | Intergenic | 1379620 | 1382922 | 236 | 1379144 | 1377849 | 1536 | 5' |
| 1 | 3 | NCO | 1308011 | 1308402 | 1308207 | Intergenic | 1308395 | 1313960 | 189 | 1304668 | 1307178 | 1029 | 5' |
| 1 | 3 | NCO | 1581077 | 1581307 | 1581192 | Intergenic | 1581384 | 1584653 | 192 | 1579418 | 1580063 | 1129 | 5' |
| 1 | 3 | NCO | 3934647 | 3935970 | 3935309 | Intergenic | 3934549 | 3929226 | 760 | 3937982 | 3936714 | 1406 | 5' |
| 1 | 3 | NCO | 4101219 | 4101796 | 4101508 | Intragenic | 4105247 | 4108761 | 3740 | 4096861 | 4102598 | 1091 | - |
| 1 | 5 | NCO | 783355 | 783607 | 783481 | Intergenic | 782524 | 782871 | 957 | 782524 | 782871 | 610 | 3' |

| | | | | | | | | | | | | | |
|---|---|---|---|---|---|---|---|---|---|---|---|---|---|
| 1 | 5 | NCO | 2906100 | 2906367 | 2906234 | Intragenic | 2907740 | 2901315 | 1507 | 2915194 | 2909645 | 3412 | - |
| 2 | 1 | NCO | 1369629 | 1370333 | 1369981 | Intergenic | 1369194 | 1364715 | 787 | 1377443 | 1372411 | 2430 | 5' |
| 2 | 1 | NCO | 3651786 | 3651856 | 3651821 | Intragenic | 3653250 | 3657324 | 1429 | 3650304 | 3651929 | 108 | 3' |
| 2 | 1 | NCO | 3712241 | 3712728 | 3712485 | Intragenic | 3713111 | 3710542 | 627 | 3701753 | 3712204 | 281 | 3' |
| 2 | 2 | NCO | 1084839 | 1085167 | 1085003 | Intergenic | 1086126 | 1085197 | 1123 | 1086126 | 1085197 | 194 | 3' |
| 2 | 2 | NCO | 4571606 | 4571994 | 4571800 | Intragenic | 4571993 | 4565179 | 193 | 4572506 | 4573306 | 1506 | 5' |
| 2 | 2 | NCO | 5129211 | 5129695 | 5129453 | Intragenic | 5129030 | 5132462 | 423 | 5133070 | 5129486 | 33 | 3' |
| 2 | 3 | NCO | 1577457 | 1577695 | 1577576 | Intergenic | 1578905 | 1577709 | 1329 | 1578905 | 1577709 | 133 | 3' |
| 2 | 3 | NCO | 2103630 | 2104176 | 2103903 | Intergenic | 2103658 | 2102234 | 245 | 2103658 | 2102234 | 1669 | 5' |
| 2 | 3 | NCO | 2431447 | 2432791 | 2432119 | Intragenic | 2431758 | 2432989 | 361 | 2431758 | 2432989 | 870 | 5' |
| 2 | 4 | NCO | 1138845 | 1139160 | 1139003 | Intergenic | 1139421 | 1140686 | 419 | 1136858 | 1138270 | 733 | 5' |
| 2 | 4 | NCO | 1724988 | 1726778 | 1725883 | Intergenic | 1725616 | 1723557 | 267 | 1732616 | 1726189 | 306 | 5' |
| 2 | 6 | NCO | 1406831 | 1409834 | 1408333 | Intergenic | 1408569 | 1409813 | 237 | 1412087 | 1408660 | 328 | 5' |
| 2 | 7 | NCO | 2140902 | 2141509 | 2141206 | Intragenic | 2141975 | 2144008 | 770 | 2140148 | 2141462 | 257 | 3' |
| 3 | 1 | NCO | 1958151 | 1958440 | 1958296 | Intragenic | 1955683 | 1958348 | 2613 | 1955683 | 1958348 | 53 | 3' |
| 3 | 1 | NCO | 250965 | 258641 | 254803 | Intragenic | 254812 | 253268 | 9 | 249805 | 254132 | 671 | 5' |
| 3 | 2 | NCO | 276341 | 276455 | 276398 | Intergenic | 276192 | 274672 | 206 | 279641 | 276700 | 302 | 5' |
| 3 | 2 | NCO | 1081240 | 1081467 | 1081354 | Intergenic | 1080906 | 1078315 | 448 | 1082865 | 1082524 | 1171 | 5' |
| 3 | 2 | NCO | 1515769 | 1517551 | 1516660 | Intergenic | 1513903 | 1515837 | 2757 | 1513903 | 1515837 | 823 | 3' |
| 3 | 2 | NCO | 4581140 | 4582923 | 4582032 | Intergenic | 4581530 | 4586355 | 502 | 4586205 | 4584061 | 2030 | 5' |
| 3 | 3 | NCO | 2083858 | 2084855 | 2084357 | Intergenic | 2082977 | 2083939 | 1380 | 2082977 | 2083939 | 418 | 3' |
| 3 | 3 | NCO | 2086583 | 2087078 | 2086831 | Intergenic | 2088183 | 2091283 | 1353 | 2082977 | 2083939 | 2892 | - |

| | | | | | | | | | | | | | |
|---|---|---|---|---|---|---|---|---|---|---|---|---|---|
| 3 | 3 | NCO | 4830502 | 4832061 | 4831282 | Intergenic | 4837382 | 4838545 | 6101 | 4825154 | 4828646 | 2636 | - |
| 3 | 5 | NCO | 162435 | 166915 | 164675 | Intragenic | 165783 | 162555 | 1108 | 162796 | 163791 | 884 | 3' |
| 3 | 6 | NCO | 1263014 | 1263243 | 1263129 | Intergenic | 1263683 | 1263130 | 555 | 1263683 | 1263130 | 2 | 3' |
| 3 | 6 | NCO | 2581995 | 2582218 | 2582107 | Intergenic | 2582554 | 2584560 | 448 | 2585848 | 2582554 | 448 | 5' |
| 3 | 7 | NCO | 532552 | 532631 | 532592 | Intergenic | 532044 | 529757 | 548 | 532044 | 529757 | 2835 | 5' |
| 3 | 7 | NCO | 714501 | 714777 | 714639 | Intergenic | 713491 | 711372 | 1148 | 717385 | 715567 | 928 | 3' |
| 3 | 7 | NCO | 2432818 | 2433337 | 2433078 | Intergenic | 2432874 | 2430332 | 204 | 2436318 | 2433522 | 445 | 5' |
| 4 | 1 | CO | 2241769 | 2241952 | 2241861 | Intergenic | 2243719 | 2242366 | 1859 | 2243719 | 2242366 | 506 | 3' |
| 4 | 1 | CO | 6409743 | 6409839 | 6409791 | Intragenic | 6409380 | 6411031 | 411 | 6416598 | 6409709 | 82 | 3' |
| 4 | 1 | CO | 4763564 | 4763778 | 4763671 | Intergenic | 4763335 | 4762198 | 336 | 4764972 | 4764226 | 555 | 5' |
| 4 | 1 | CO | 1769653 | 1769855 | 1769754 | Intergenic | 1771217 | 1774581 | 1463 | 1763019 | 1769007 | 747 | 3' |
| 4 | 2 | CO | 2897761 | 2898274 | 2898018 | Intragenic | 2897202 | 2899812 | 816 | 2900346 | 2898748 | 731 | 3' |
| 4 | 2 | CO | 2156378 | 2156697 | 2156538 | Intragenic | 2156933 | 2159825 | 396 | 2154499 | 2156796 | 259 | 3' |
| 4 | 2 | CO | 1103485 | 1104399 | 1103942 | Intragenic | 1104491 | 1101578 | 549 | 1097079 | 1104711 | 769 | 5' |
| 4 | 3 | CO | 991973 | 992518 | 992246 | Intergenic | 987104 | 991660 | 5142 | 987104 | 991660 | 586 | 3' |
| 4 | 3 | CO | 2827908 | 2828034 | 2827971 | Intragenic | 2829037 | 2828414 | 1066 | 2826696 | 2828091 | 120 | 3' |
| 4 | 4 | CO | 4020052 | 4020052 | 4020052 | Intragenic | 4022373 | 4018979 | 2321 | 4022373 | 4018979 | 1073 | - |
| 4 | 4 | CO | 1242957 | 1242937 | 1242947 | Intragenic | 1243989 | 1242213 | 1042 | 1243989 | 1242213 | 734 | 3' |
| 4 | 4 | CO | 3095682 | 3095844 | 3095763 | Intragenic | 3096833 | 3095256 | 1070 | 3096833 | 3095256 | 507 | 3' |
| 4 | 5 | CO | 2588267 | 2588267 | 2588267 | Intragenic | 2590651 | 2586799 | 2384 | 2590651 | 2586799 | 1468 | - |
| 4 | 5 | CO | 519239 | 519238 | 519239 | Intragenic | 519690 | 519068 | 452 | 519690 | 519068 | 171 | 3' |
| 4 | 5 | CO | 4088627 | 4088927 | 4088777 | Intragenic | 4088888 | 4083052 | 111 | 4085873 | 4088516 | 261 | 5' |

| | | | | | | | | | | | | | |
|---|---|---|---|---|---|---|---|---|---|---|---|---|---|
| 4 | 5 | CO | 1690104 | 1690199 | 1690152 | Intergenic | 1686464 | 1689112 | 3688 | 1686464 | 1689112 | 1040 | - |
| 4 | 6 | CO | 3270124 | 3270115 | 3270120 | Intragenic | 3268921 | 3275655 | 1199 | 3275655 | 3268921 | 1199 | - |
| 4 | 6 | CO | 244261 | 244399 | 244330 | Intragenic | 232404 | 234437 | 11926 | 256364 | 241700 | 2630 | - |
| 4 | 7 | CO | 1283464 | 1283835 | 1283650 | Intergenic | 1280064 | 1283137 | 3586 | 1280064 | 1283137 | 513 | 3' |
| 5 | 1 | CO | 2735360 | 2735689 | 2735525 | Intragenic | 2736794 | 2733269 | 1270 | 2729715 | 2734694 | 831 | 3' |
| 5 | 2 | CO | 2539850 | 2540457 | 2540154 | Intragenic | 2541240 | 2539574 | 1087 | 2541240 | 2539574 | 580 | 3' |
| 5 | 2 | CO | 3265637 | 3403422 | 3334530 | Intragenic | 3335019 | 3334243 | 490 | 3335019 | 3334243 | 287 | 3' |
| 5 | 3 | CO | 4879455 | 4879922 | 4879689 | Intragenic | 4880712 | 4875548 | 1024 | 4876160 | 4878151 | 1538 | - |
| 5 | 3 | CO | 1482361 | 1482496 | 1482429 | Intragenic | 1480265 | 1484912 | 2164 | 1486865 | 1483233 | 805 | 3' |
| 5 | 4 | CO | 1241191 | 1241185 | 1241188 | Intragenic | 1241617 | 1239555 | 429 | 1240541 | 1241502 | 314 | 3' |
| 5 | 4 | CO | 3145281 | 3145281 | 3145281 | Intragenic | 3144232 | 3145383 | 1049 | 3144232 | 3145383 | 102 | 3' |
| 5 | 5 | CO | 3163568 | 3164357 | 3163963 | Intergenic | 3165976 | 3164677 | 2014 | 3165976 | 3164677 | 715 | 3' |
| 5 | 5 | CO | 1802626 | 1803383 | 1803005 | Intergenic | 1805512 | 1803198 | 2508 | 1805512 | 1803198 | 194 | 3' |
| 5 | 6 | CO | 1761355 | 1761172 | 1761264 | Intragenic | 1763531 | 1760877 | 2268 | 1763531 | 1760877 | 387 | 3' |
| 5 | 7 | CO | 2221930 | 2221752 | 2221841 | Intergenic | 2219283 | 2221720 | 2558 | 2219283 | 2221720 | 121 | 3' |
| 5 | 7 | CO | 2858910 | 2859253 | 2859082 | Intragenic | 2859613 | 2861625 | 532 | 2857948 | 2859360 | 279 | 3' |
| 6 | 1 | CO | 1774685 | 1774775 | 1774730 | Intergenic | 1774121 | 1771753 | 609 | 1771217 | 1774581 | 149 | 3' |
| 6 | 1 | CO | 5629313 | 5629308 | 5629311 | Intragenic | 5629062 | 5631539 | 249 | 5629062 | 5631539 | 2229 | 5' |
| 6 | 1 | CO | 2566652 | 2566316 | 2566484 | Intragenic | 2565821 | 2569888 | 663 | 2565821 | 2569888 | 3404 | 5' |
| 6 | 1 | CO | 6704547 | 6705209 | 6704878 | Intragenic | 6700909 | 6704051 | 3969 | 6709167 | 6704373 | 505 | 3' |
| 6 | 1 | CO | 513681 | 513863 | 513772 | Intergenic | 511932 | 513363 | 1840 | 511932 | 513363 | 409 | 3' |
| 6 | 2 | CO | 2609095 | 2609462 | 2609279 | Intragenic | 2609109 | 2608382 | 170 | 2608232 | 2609461 | 183 | 5' |

| | | | | | | | | | | | | | |
|---|---|---|---|---|---|---|---|---|---|---|---|---|---|
| 6 | 2 | CO | 3901065 | 3901144 | 3901105 | Intragenic | 3898236 | 3902464 | 2869 | 3907654 | 3900293 | 812 | 3' |
| 6 | 2 | CO | 1870942 | 1870942 | 1870942 | Intergenic | 1873389 | 1871254 | 2447 | 1873389 | 1871254 | 312 | 3' |
| 6 | 2 | CO | 233330 | 233903 | 233617 | Intergenic | 231202 | 233393 | 2415 | 231202 | 233393 | 224 | 3' |
| 6 | 3 | CO | 1074212 | 1073823 | 1074018 | Intergenic | 1074557 | 1075246 | 540 | 1074557 | 1075246 | 1229 | 5' |
| 6 | 4 | CO | 4144713 | 4144780 | 4144747 | Intragenic | 4144475 | 4144879 | 272 | 4144475 | 4144879 | 133 | 3' |
| 6 | 4 | CO | 1964912 | 1965613 | 1965263 | Intragenic | 1965922 | 1964957 | 660 | 1965922 | 1964957 | 306 | 3' |
| 6 | 4 | CO | 2657912 | 2658268 | 2658090 | Intragenic | 2655675 | 2659774 | 2415 | 2662633 | 2658702 | 612 | 3' |
| 6 | 4 | CO | 1188529 | 1188529 | 1188529 | Intragenic | 1187147 | 1194489 | 1382 | 1192127 | 1191114 | 2585 | - |
| 6 | 4 | CO | 560791 | 560980 | 560886 | Intragenic | 560715 | 561631 | 171 | 560715 | 561631 | 746 | 5' |
| 6 | 5 | CO | 2596510 | 2591967 | 2594239 | Intergenic | 2593778 | 2591667 | 461 | 2598360 | 2596426 | 2188 | 5' |
| 6 | 5 | CO | 1894648 | 1894573 | 1894611 | Intragenic | 1892792 | 1895345 | 1819 | 1892792 | 1895345 | 735 | 3' |
| 6 | 6 | CO | 1586463 | 1586839 | 1586651 | Intergenic | 1588789 | 1587943 | 2138 | 1583859 | 1586364 | 287 | 3' |
| 6 | 6 | CO | 643765 | 643765 | 643765 | Intergenic | 644583 | 646889 | 818 | 644583 | 646889 | 3124 | 5' |
| 6 | 7 | CO | 1603662 | 1603799 | 1603731 | Intragenic | 1602571 | 1604896 | 1160 | 1602571 | 1604896 | 1166 | - |
| 6 | 7 | CO | 3372959 | 3372959 | 3372959 | Intragenic | 3372326 | 3373289 | 633 | 3372326 | 3373289 | 330 | 3' |
| 4 | 1 | NCO | 2853921 | 2854674 | 2854298 | Intragenic | 2854145 | 2856391 | 153 | 2859907 | 2855965 | 1668 | 5' |
| 4 | 1 | NCO | 2960385 | 2960629 | 2960507 | Intergenic | 2960861 | 2962204 | 354 | 2960861 | 2962204 | 1697 | 5' |
| 4 | 3 | NCO | 983870 | 984259 | 984065 | Intragenic | 985148 | 982841 | 1084 | 985148 | 982841 | 1224 | - |
| 4 | 3 | NCO | 2985663 | 2986852 | 2986258 | Intragenic | 2986992 | 2986189 | 735 | 2986992 | 2986189 | 69 | 3' |
| 4 | 4 | NCO | 2770039 | 2771687 | 2770863 | Intragenic | 2770298 | 2772386 | 565 | 2775463 | 2770315 | 548 | 3' |
| 4 | 5 | NCO | 3154517 | 3156962 | 3155740 | Intragenic | 3158022 | 3154433 | 2283 | 3150248 | 3154919 | 821 | 3' |
| 4 | 7 | NCO | 2773139 | 2774456 | 2773798 | Intragenic | 2773415 | 2772534 | 383 | 2769297 | 2774201 | 404 | 5' |

| | | | | | | | | | | | | | |
|---|---|---|---|---|---|---|---|---|---|---|---|---|---|
| 5 | 3 | NCO | 1482118 | 1482295 | 1482207 | Intragenic | 1480265 | 1484912 | 1942 | 1486865 | 1483233 | 1027 | - |
| 6 | 1 | NCO | 2041689 | 2042163 | 2041926 | Intragenic | 2045644 | 2041576 | 3718 | 2045644 | 2041576 | 350 | 3' |
| 6 | 1 | NCO | 6123851 | 6125477 | 6124664 | Intergenic | 6123075 | 6122609 | 1589 | 6123075 | 6122609 | 2055 | - |
| 6 | 1 | NCO | 1472641 | 1472935 | 1472788 | Intragenic | 1472961 | 1466603 | 173 | 1468050 | 1472546 | 242 | 5' |
| 6 | 2 | NCO | 3912210 | 3912700 | 3912455 | Intergenic | 3916364 | 3933723 | 3909 | 3925959 | 3916427 | 3972 | - |
| 6 | 2 | NCO | 4548348 | 4549146 | 4548747 | Intergenic | 4548351 | 4547389 | 396 | 4548351 | 4547389 | 1358 | 5' |
| 6 | 2 | NCO | 5637286 | 5637536 | 5637411 | Intragenic | 5637379 | 5639868 | 32 | 5637379 | 5639868 | 2457 | 5' |
| 6 | 3 | NCO | 188381 | 189284 | 188833 | Intragenic | 191345 | 188587 | 2513 | 185649 | 188736 | 97 | 3' |
| 6 | 3 | NCO | 1577409 | 1578150 | 1577780 | Intragenic | 1578905 | 1577709 | 1126 | 1578905 | 1577709 | 71 | 3' |
| 6 | 3 | NCO | 2266423 | 2267736 | 2267080 | Intragenic | 2268310 | 2272073 | 1231 | 2256651 | 2268107 | 1028 | - |
| 6 | 3 | NCO | 5158688 | 5159572 | 5159130 | Intergenic | 5158357 | 5157443 | 773 | 5162299 | 5159467 | 337 | 3' |
| 6 | 4 | NCO | 1457320 | 1457782 | 1457551 | Intragenic | 1459984 | 1458078 | 2433 | 1454185 | 1458015 | 464 | 3' |
| 6 | 7 | NCO | 1664266 | 1664760 | 1664513 | Intergenic | 1663898 | 1661320 | 615 | 1667080 | 1666223 | 1710 | 5' |
| 6 | 7 | NCO | 1795858 | 1798012 | 1796935 | Intergenic | 1741012 | 1739687 | 55923 | 1738173 | 1741012 | 55923 | - |
